# IFITM3 blocks influenza virus entry by sorting lipids and stabilizing hemifusion

**DOI:** 10.1101/2022.09.01.506185

**Authors:** Steffen Klein, Gonen Golani, Fabio Lolicato, Carmen Lahr, Daniel Beyer, Alexia Herrmann, Moritz Wachsmuth-Melm, Nina Reddmann, Romy Brecht, Mehdi Hosseinzadeh, Androniki Kolovou, Jana Makroczyova, Sarah Peterl, Martin Schorb, Yannick Schwab, Britta Brügger, Walter Nickel, Ulrich S. Schwarz, Petr Chlanda

## Abstract

Interferon-induced transmembrane protein 3 (IFITM3) inhibits the entry of numerous viruses through undefined molecular mechanisms. IFITM3 localizes in the endosomal-lysosomal system and specifically impacts virus fusion with target cell membranes. We found that IFITM3 induces local lipid sorting, resulting in an increased concentration of lipids disfavoring viral fusion at the hemifusion site. This increases the energy barrier for fusion pore formation and the hemifusion dwell time, promoting viral degradation in lysosomes. *In situ* cryo-electron tomography captured IFITM3-mediated arrest of influenza A virus membrane fusion. Observation of hemifusion diaphragms between viral particles and late endosomal membranes confirmed hemifusion stabilization as a molecular mechanism of IFITM3. The presence of the influenza fusion protein hemagglutinin in post-fusion conformation close to hemifusion sites further indicated that IFITM3 does not interfere with the viral fusion machinery. Collectively, these findings show that IFITM3 induces lipid sorting to stabilize hemifusion and prevent virus entry into target cells.

## INTRODUCTION

Perhaps the most decisive step during infection is the release of the viral genome into the cytoplasm, for which membrane enveloped viruses rely on their fusion machinery. Many enveloped viruses such as influenza A virus (IAV) are important human pathogens and blocking viral entry is one of the most efficient countermeasures the host cell can take. An IAV infection starts with the virus attaching to the host cell by binding the trimeric surface glycoprotein hemagglutinin (HA) to sialic acid residues on the cell surface, followed by clathrin-mediated endocytosis^1^ or macropinocytosis^2^. The acidic environment in late endosomes induces complex conformational changes in the HA glycoprotein and triggers the disassembly of the viral matrix 1 (M1) layer. The pre-fusion HA structure undergoes extensive refolding with a loop-to-helix transition of the HA2 domain that leads to the formation of an extended HA intermediate^3, 4^, which anchors the N-terminal fusion peptide into the late endosomal membrane. The extended intermediate folds back into an energetically more stable coiled-coil post-fusion state. The HA refolding provides the mechanical work bringing opposing membranes to proximity below 1–2 nm^5^ and forming a hemifusion stalk^6–8^, which subsequently expands to a hemifusion diaphragm. The diaphragm is a mechanically stressed lipid structure, and transient membrane pores tend to open within it. The magnitude of the energy barrier for pore expansion depends on the lipid composition and determines the stability and dwell time of the hemifusion state. Fusion-pore opening is promoted by lipids with positive, spontaneous curvature (e.g., lysophosphatidylcholine (LPC)) and inhibited by lipids with negative spontaneous curvature (e.g., cholesterol) in the virus and host distal monolayers^9^. While HA-mediated membrane fusion has been extensively studied *in vitro* and *in silico*^10^, little is known about how this process is controlled in cells and how host cell restriction factors interfere with this process on the molecular level.

Interferon-induced transmembrane protein 3 (IFITM3) has evolved as a broad-spectrum cellular restriction factor that blocks cytosolic entry of IAV^11^ and various late penetrating viruses^12^ by interfering with the fusion process. IFITM3 is a small (15 kDa) single-domain transmembrane protein^13^ localized in the endosomal-lysosomal system^14–16^. It comprises two amphipathic helices facing the cytosolic side of the endosomal membrane^17^, which are essential for its antiviral properties^18^. IFITM3 is post-translationally modified by S-palmitoylation^19^, essential for the functional incorporation of IFITM3 into the endosomal membrane and its antiviral activity^20, 21^. Fluorescence microscopy-based live-cell imaging revealed that while IFITM3 effectively blocks the release of the viral genome into the cytoplasm, lipid mixing between the viral and endosomal membrane could still be observed^22^. Based on these data, two alternative models were proposed. In the so-called ‘hemifusion stabilization’ model, IFITM3 interferes with the fusion pore formation in a direct or indirect mode of action, whereas, in the ‘fusion decoy’ model, IFITM3 increases the number of endosomal intraluminal vesicles (ILVs) which may serve as a decoy target for virus membrane fusion^22^. *In vitro* experiments suggest that IFITM3 may function by altering the biophysical properties of the membrane^23^ or by changing the lipid composition and cholesterol levels of the endosomal membranes^24–26^. However, despite many studies on IFITM3, the exact mode of action remains to be clarified. It has not yet been addressed on the molecular level and directly inside the infected host cells.

## RESULTS

### IFITM3 does not alter the number of intraluminal vesicles in the late endosomal lumen

While IFITM3 expression can be induced by type I interferon (Figure 1A, Figure S1A-C), interferon treatment leads to the induction of additional restriction factors that may inhibit viral entry. Hence we established an A549 cell line stably overexpressing non-tagged IFITM3 (A549-IFITM3), which showed a significant 3.9–fold increase in IFITM3 expression level compared to A549 cells induced with FNβ-1b (Figure 1A, Figure S1A and B). The stable overexpression of IFITM3 did not induce the expression of IFITM1 or IFITM2 (Figure S1C). Fluorescent confocal microscopy showed that IFITM3 partially colocalizes with the late endosomal marker Rab7 (Pearson’s *r*=0.51, SD=0.08, n=9) and the lysosomal marker LAMP1 (Pearson’s *r*=0.64, SD=0.14, n=10) (Figure S2), consistent with previous studies on endogenous IFITM3^14–16, 27^, showing that the cellular localization of IFITM3 in the A549-IFITM3 cell line is unaltered. As expected, both IFNβ-1b treatment and stable IFITM3 expression increased the number of IFITM3-positive cellular organelles in A549 cells (Figure 1B–H). Three-dimensional segmentation (Figure 1C, 1E, and 1G) revealed a 3.1–fold increase in the number of organelles (Figure 1H) and a moderate but significant 1.3–fold increase in average organelle volume of 0.39 µm^3^ (SD=0.41 µm^3^, n=2,402) (Figure S1D) when compared to the IFNβ-1b treated A549 cell line. To characterize the antiviral effect of the A549-IFITM3 cell line, cells were infected and immunolabelled against M2 (Figure 1I–P). While A549 cells showed an infection rate of 98.8% (n=1,136) (Figure 1O), the infection rate in A549-IFITM3 was dramatically decreased (11.5–fold) as only 8.2% (n=2,009) of the cells were infected (Figure 1P). These data agree with the previously reported antiviral properties of IFITM3^11^. To further validate the antiviral properties of the A549-IFITM3 cell line, we performed a β-lactamase (Blam) cell entry assay utilizing influenza A virus-like particles (VLPs) expressing M1-Blam^28^. Our data show an IFNβ-1b concentration-dependent decrease in relative viral entry with an up to 8.7–fold decrease. In comparison, A549-IFITM3 showed a relative viral entry inhibition of 10.7–fold (Figure 1Q and Figure S1I). Overall these data are consistent with previous studies^14, 22^ and show that viral-induced membrane fusion in late endosomes is inhibited in A549-IFITM3 cells.

**Figure 1.**
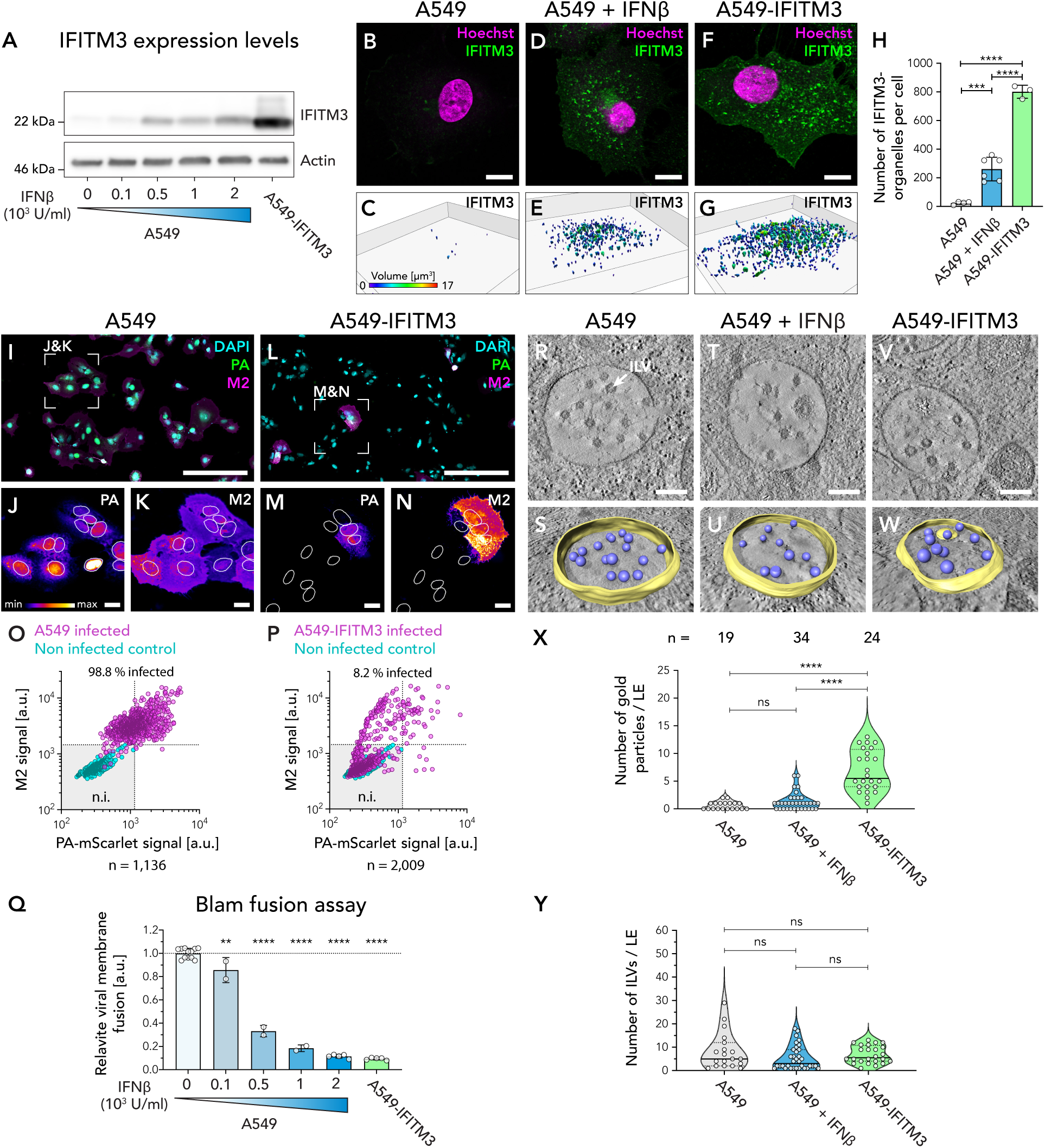
IFITM3 inhibits IAV infection by blocking viral membrane fusion without altering the number of ILVs in the late endosomal lumen. **(A)** Immunoblot analysis of IFNβ-1b-treated A549 cells and A549-IFITM3 cells. See Figure S1A for the expression quantification and Figure S1B for the full blot. (B–G) Confocal microscopy analysis of A549 cells **(B and C)**, A549 cells treated with 1×10^3^ units/ml IFNβ-1b **(D and E)**, and A549-IFITM3 cells **(F and G)**. Cells were immunolabeled against IFITM3 (green), and the nucleus was stained with Hoechst33342 (magenta). The top row **(B, D, F)** shows a composite central slice of a representative cell. The bottom row **(C, E, G)** shows the three-dimensional segmentation of the IFITM3 signal. Individual objects are color-labeled according to their volume. (H) The number of individual IFITM3 organelles per cell, as quantified by three-dimensional IFITM3 signal segmentation (C, E, G). The bar graph shows the mean value with standard deviation (SD). An unpaired t-test was performed. **(I–N)** IAV infection assay for A549 cells (I–K) and A549-IFITM3 cells (L–N). Cells were infected with the fluorescent reporter virus A/WSN/1933-PA-mScarlet (MOI = 3) (green) and fixed 24 hours post-infection (hpi). The viral protein M2 (magenta) was labeled by immunofluorescence, and nuclei were labeled with DAPI (cyan). Composite overview images for both conditions are shown (I and L). For each condition, a magnified area (white squares) for the signals of PA-mScarlet and M2 are shown in the bottom row (J, K, M, N) using a fire lookup table. Individual nuclei were segmented (indicated with white outlines), allowing for measuring the average signal intensity for PA-mScarlet and M2 in the segmented area. **(O and P)** Quantification of infection rate for A549 cells (O) and A549-IFITM3 cells (P). For each segmented nucleus, the signal intensity for PA-mScarlet and M2 is shown as a scatter plot for infected cells (magenta) and non-infected control (cyan). Two thresholds of M1≤1,150 a.u. and PA-mScarlet≤1,450 a.u. (dotted lines) were defined for non-infected (n.i.) cells (shown as a grey square), and accordingly, all other cells are defined as infected. See Figure S1E–H for immunofluorescence images and scatter plots of the non-infected control only. **(Q)** Blam-based cell entry assay for IFNβ-1b-treated A549 and A549-IFITM3 cells infected with M1-Blam influenza A VLPs. An unpaired t-test was performed. See Figure S1I for fluorescence microscopy images of all samples. **(R, T, V)** Tomographic slices of A549 cells, IFNβ-1b-treated A549 cells (T), and A549-IFITM3 cells (V). A representative late endosome was selected for each sample with the typical multivesicular morphology. One ILV is indicated (white arrow In R). 10 slices were averaged (10.9 nm), and low pass filtered (3.09 nm). See Figure S3 for a gallery of late endosomes for each condition. **(S, U, W)** Three-dimensional segmentation of the late endosomes is shown in (R, T, V). The late endosomal membrane (yellow) and individual ILVs (purple) were segmented. **(X)** Quantification of anti-IFITM3 immunogold particles per late endosome. See Figure S3 for a gallery of immune-gold-labeled late endosomes. **(Y)** Quantification of the number of ILVs per endosome. Statistical significance: * for p<0.05, ** for p<0.01, *** for p<0.001 **** for p<0.0001. Scale bars: (B–F) 10 µm, (I and L) 200 µm, (J, K, M, N) 20 µm, (R, T, V) 200 nm.

To test the ‘fusion decoy’ hypothesis^22^, we determined morphological changes of late endosomes in A549-IFITM3 cells by electron tomography (Figure 1R–Y). As the ‘fusion decoy’ hypothesis suggested an increase in ILVs, we analyzed late endosomes in a total of 77 tomograms of A549 cells (n=19), A549 cells treated with 2×10^3^ units/ml IFNβ-1b (n=34), and A549-IFITM3 cells (n=24). The endosomal membrane and ILVs were 3D segmented (Figure 1S, 1U, 1W) to quantify the endosomal volume (Figure S1J). Consistent with confocal microscopy data (Figure S1D), late endosomal volume in A549-IFITM3 cells was modestly (1.6–fold) but significantly increased compared to non-treated A549 cells (Figure S1J). Immuno-gold-labeling against IFITM3 showed that late endosomes in A549-IFITM3 cells displayed a significantly higher (5.3–fold) IFITM3 labeling than A549 cells (Figure 1X and Figure S3). This data is consistent with the increased IFITM3 expression levels of A549-IFITM3 cells (Figure 1A) and shows that IFITM3 is specifically localized to multivesicular organelles. Importantly, our data revealed no significant difference in the number of ILVs per endosome between A549 cells, IFNβ-1b-treated A549 cells, or A549-IFITM3 cells (Figure 1Y), hence contradicting the assumptions of the ‘fusion decoy’ model.

### IFITM3 modulates the local lipid composition of the endosomal membrane

We next analyzed the impact of IFITM3 on the lipid composition of these organelles. It has been previously reported that IFITM3 alters the cholesterol level in endosomes^24^. To determine whether IFITM3 directly interacts with cholesterol, we performed UV cross-linking using a photo-reactive and clickable cholesterol analog (pacChol) as a probe to map cholesterol-IFITM3 interactions in living cells. Western blot analysis of immunoprecipitated pacChol showed that IFITM3 and cholesterol directly interact (Figure 2A), as reported recently^26^.

**Figure 2.**
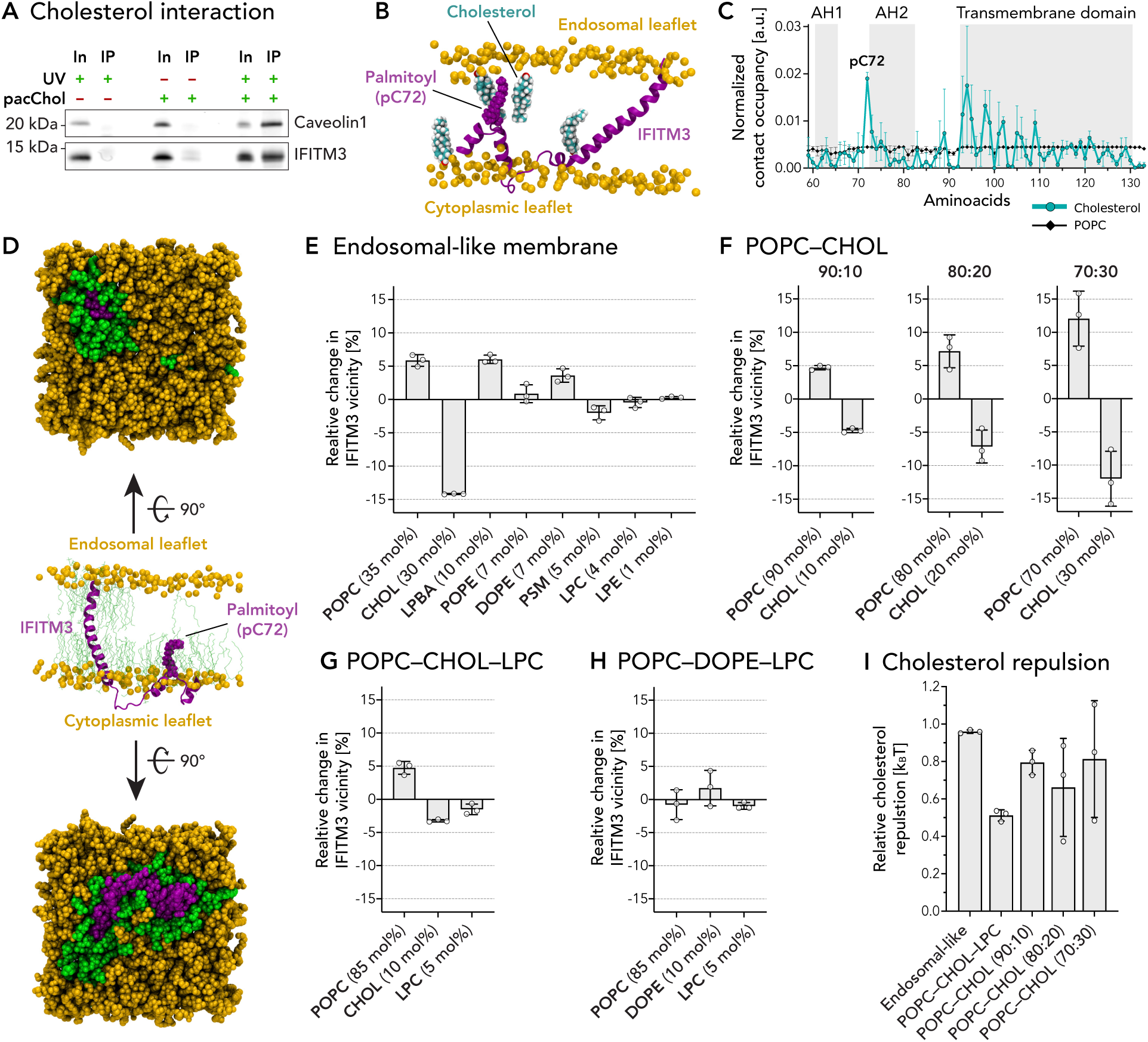
Molecular dynamics simulations predict IFITM3-induced lipid sorting by selective lipid binding. **(A)** Analysis of IFIMT3-cholesterol interactions in A549-IFITM3 cells. Cells were treated with pacChol, UV cross-linked, and immunoblot was performed against IFITM3 after immunoprecipitation (IP). Caveolin1 was used as a positive control. Full blots can be found in Figure S4A and S4B. **(B)** A representative snapshot of the interaction between cholesterol molecules and IFITM3 in a binary POPC–cholesterol (90:10) system. IFITM3 is represented as a ribbon (magenta), and the palmitoyl group of Cys72 is represented as an atom sphere model, having a van der Waals (vdW) radius of their respective atoms. Phosphorus atoms of the POPC head groups are represented as an atom sphere model (orange). POPC tails are not shown. Cholesterol molecules in contact (distance < 0.4 nm) with IFITM3 are represented as an atom sphere model (cyan), with oxygen atoms in red and hydrogen atoms in white. Non-interacting cholesterol molecules are not shown. **(C)** Normalized contact occupancy between IFITM3 and POPC (black graph) or cholesterol (cyan graph) in a binary POPC–cholesterol (90:10) model membrane as shown in (B). A contact is defined if the distance between any atoms of IFITM3 and POPC (or cholesterol) is less than 0.4 nm. Data are normalized based on the total number of lipids of each lipid type. The sequence sections for the two amphipathic helix domains (AH1 and AH2) and the transmembrane domain are indicated. Error bars are shown as standard deviation (SD). **(D)** A representative snapshot of IFITM3 (magenta), with palmitoylated Cys^72^, embedded in a late endosomal-like model membrane (orange). The central side view shows IFITM3 as a ribbon model, with the palmitoyl-group at Cys72 as an atom sphere model. Lipid tails of lipids in close vicinity to IFITM3 (< 0.4 nm) are shown as stick model (green), and all other lipid tails are hidden. The top views of both the endosomal (top) and cytoplasmic (bottom) leaflets are shown. All molecules are shown as an atom sphere model, with IFITM3 in magenta, lipids in close vicinity to IFITM3 (< 0.4 nm) in green, and all other lipids in orange. **(E–H)** Relative changes of lipid concentrations in the vicinity of IFITM3 (< 0.4 nm) for different lipid systems: Late-endosomal like lipid system (E); binary POPC–cholesterol lipid system with cholesterol concentrations of 10, 20, and 30 mol% IFITM3 (F); ternary POPC–cholesterol–LPC lipid system (G); and ternary POPC–DOPE–LPC lipid system (H). **(I)** Cholesterol repulsion from IFITM3 for all studied membrane systems. All plots in (E–I) show the mean of three independent MD simulation runs. Individual values are indicated. The error bar represents the standard deviation (SD).

To further characterize the IFITM3-cholesterol interaction, we employed atomistic molecular dynamics (MD) simulations of IFITM3 embedded in different lipid compositions. As palmitoylated Cys72 is essential for IFITM3’s antiviral properties^21^, this modification was incorporated in all MD simulations. First, we analyzed the contact occupancy for each amino acid of IFITM3 in a binary POPC–cholesterol (90:10) model membrane (Figure 2B). Our data revealed that while most IFITM3 amino acids showed a low probability of interactions with cholesterol, there is an increased interaction probability in the cytosolic region of the transmembrane domain, and the palmitoylated Cys72 displayed the strongest cholesterol-interaction probability of 1.9×10^-2^ (SD=1.3×10^-3^, n=3) (Figure 2C). Next, we investigated the IFITM3-cholesterol interactions in a complex membrane environment by simulating IFITM3 in a late endosomal-like membrane^29, 30^ composed of eight different lipids (performed three times for a cumulative time of 18 µs). The IFITM3–lipid interaction landscape was investigated by comparing the molar fraction of lipids surrounding IFITM3 with the lipid bulk concentrations at a radial distance of 0.4 nm around IFITM3 (Figure 2D, green lipids). Our data highlight a statistically significant (p<0.000001, Figure S4C) depletion of cholesterol from IFITM3’s vicinity by 14.2% (SD=0.1%, n=3) (Figure 2E). The repelled cholesterol is mainly replaced by the two lipids POPC and LBPA (Figure 2E), which showed a statistically significant (Figure S4C) enrichment in IFITM3’s vicinity of 5.9% and 6.0%, respectively. To understand if the cholesterol levels in the membrane influence this IFITM3-induced lipid-sorting, we analyzed binary POPC–cholesterol lipid systems at cholesterol concentrations of 10, 20, and 30 mol%. We found that the IFITM3-induced cholesterol depletion activity can be observed in all tested cholesterol concentrations in a linear concentration-dependent manner (Figure 2F). To further analyze whether the IFITM3-induced cholesterol depletion is due to cholesterol’s intrinsic negative curvature, we simulated a POPC–cholesterol–LPC (80:10:5) system where cholesterol depletion but no LPC accumulation was observed (Figure 2G). Next, we replaced cholesterol in this system with DOPE, a lipid with an intrinsic negative curvature comparable to cholesterol. Surprisingly, DOPE is not depleted in the IFITM3 vicinity (Figure 2H), suggesting that the IFITM3-mediated cholesterol depletion activity is not dependent on the intrinsic curvature of the lipid but rather cholesterol specific. Finally, we calculated the relative repulsion energy of cholesterol from IFITM3 compared to all of the other lipids in the system based on the de-mixing entropy of cholesterol depletion near IFITM3 (Figure 2D, green lipids) and its accumulation in bulk (Figure 2D, orange lipids). We found the repulsion strength in the endosomal-like lipid composition to be 0.96 kBT (SD=0.01 kBT, n=3). Repulsions calculated from membrane models with different lipid compositions were similar (Figure 2I), indicating that the repulsion is independent of the cholesterol concentration.

The observed selective cholesterol binding to the transmembrane domain and the palmitoylated Cys72 might be important for the specific localization of IFITM3 to cholesterol-rich late endosomal membranes. However, the local cholesterol repulsion ability of IFTM3 leads to local lipid sorting in the late endosomal membrane.

### IFITM3-induced lipid sorting increases the energy barrier for fusion pore formation

Previous experimental studies revealed that IFITM3 does not affect the initiation of fusion and lipid mixing, but it blocks membrane fusion pore formation^22, 23, 31^. We suggest that the observed cholesterol repulsion from the IFITM3 vicinity (Figure 2E-I) and its accumulation in the virus and hemifusion diaphragm membranes are the primary force behind the fusion inhibition. The hemifusion diaphragm rim is a three-way junction between the endosomal and viral membranes (Figure 3A), characterized by a strong lipid tilt and splay, which decays at a 3–4 nm distance from the rim^32–34^. As a result, the transmembrane domain of IFITM3 is repelled from it and cannot pass the rim to the diaphragm and virus membranes (Figure 3A, TM domain repulsion region). Since the amphipathic helices and the Cys72 domains of IFITM3 span about 2.5 nm from the transmembrane domain (Figure 3A), it is unlikely that any part of the IFITM3 protein can reach the diaphragm. In such a setting, the hemifusion diaphragm rim and its surroundings act as a semipermeable barrier between the compartments, allowing only lipids to pass but not IFITM3. As a result of IFITM3-induced cholesterol repulsion, cholesterol is likely to be enriched in IFITM3-free regions of the hemifusion diaphragm and virus membranes and be replaced by more positively curved lipids (Figures 3B), causing local lipid-sorting. Thus, the mean intrinsic curvature of the hemifusion diaphragm monolayers shifts to more negative values leading to a decrease in mechanical stress in the diaphragm^32, 34, 35^ and an increase in the membrane pore line tension^36–38^, which in turn increases the energy barrier of fusion pore expansion and inhibits fusion.

**Figure 3.**
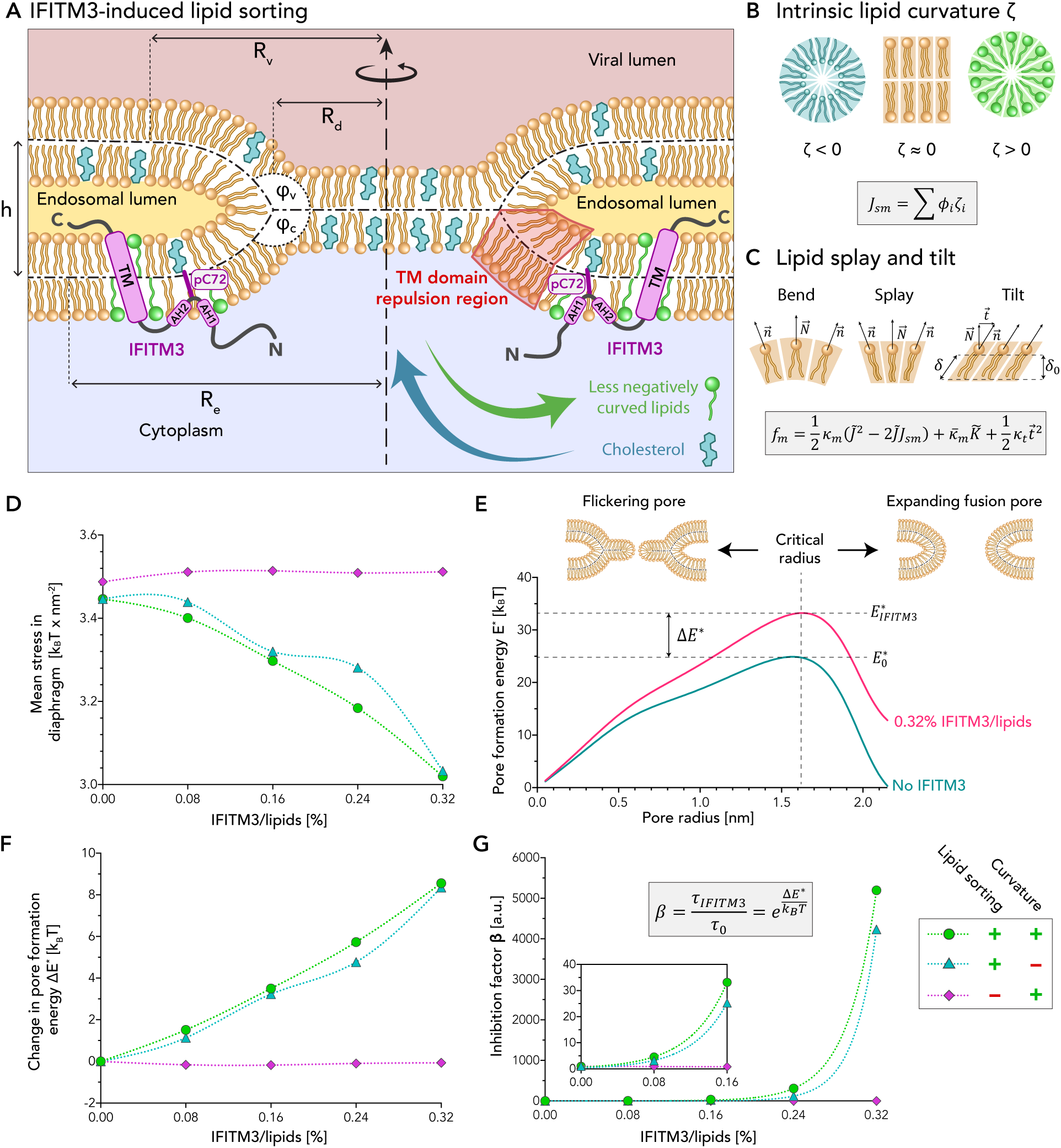
Continuum membrane modeling predicts that IFITM3-induced lipid sorting increases the energy barrier for fusion pore formation and thus stabilizes the hemifusion state. **(A)** Schematic representation of a hemifusion diaphragm during viral membrane fusion at the late endosome. The viral lumen (red), endosomal lumen (yellow), and cytoplasmic lumen (blue) are indicated. The geometry of the fusion site is defined by: The hemifusion diaphragm radius, *R_d_*; the distance between the mid-planes of fusing membranes, ℎ; and the inner angles in the cytosol and virus lumen side, *φ_c_* and *R_d_*, respectively. The radii of the fusion site, *R_v_* and *R_e_*, are defined as the distance from the center to where tilt and splay deformation vanish. The geometry of the fusion site is not fixed and is subjected to energy minimization. The main structural features of IFITM3 are shown in purple: The transmembrane domain I, the two amphipathic helixes (AH1 and AH2), and the palmitoylated cystine 72 (pC72), which interacts with a cholesterol molecule. The ‘TM domain repulsion region’ (dark red) represents the area near the diaphragm rim with a very strong tilt and splay. The TM domain of the IFITM3 is repelled from this region. IFITM3-induced lipid sorting is indicated with the blue and green arrows. **(B)** Schematic models of different intrinsic lipid curvatures, ζ_%_. The monolayer spontaneous curvature, *J_sm_*, is the averaged sum of all the lipid components with Φ_%_ being the mole fraction of each component (Equation 5, METHODS). **(C)** Theory of lipid splay-tilt and lipid monolayer energy density, *f_m_* (Equation 4, METHODS). Schematic representations of lipid bend, splay, and tilt are shown, with the lipid director, 푛)⃗; the normal to the mid-plane director, N)⃗; the lipid tilt director, t; the length of lipid tails, δ; and relaxed lipid tail length, δ_(_. Monolayer bending rigidity, *K_m_*; lipid splay, *j*; monolayer spontaneous-curvature, *j_m_*; monolayer saddle- splay modulus, *K_m_*; lipid saddle-splay, *K*3; tilt modulus, *K*_)_. (D–G) Results of the continuous membrane modeling. The following parameters were used in all simulations: *K_m_* = 14.4 k_*_T, *K*_m_ = −7.2 k_*_T, *K*_)_ = 40 mN/m, δ_(_ = 1.5 nm, ζ_"+,-_ = −0.5 nm^./^, ζ_(_ = −0.1nm^./^, Φ_"+,-_ = 0.3 and *λ*_(_ = 15 pN. The lipid distribution scenarios used in (D, F, and G) are indicated in the legend. **(D)** Mean stress due to splay and tilt deformations in the diaphragm as a function of IFITM3 concentration. **(E)** Pore formation energy, *K_pore_*, as a function of the pore radius in a model without IFITM3 (magenta graph) and with a 0.32% IFITM3/lipid ratio (blue-green graph). The ‘with sorting; with curvature’ scenario was used here. The critical radius represents the transition from flickering-pore to expanding fusion-pore. **(F)** Change in fusion- pore formation energy, Δ^∗^ = *E^*^*^∗^(Φ_121345_) − *E^*^*^∗^, as a function of IFITM3 concentration, with *E^*^*^∗^ being the energy barrier without IFITM3 and *E^*^*^∗^(Φ_121345_) being the energy barrier with the IFITM3 mole fraction, Φ_121345_. **(G)** Inhibition factor, *β*, of IFITM3 (Equation 18, METHODS), which describes the ratio between the time of pore opening with IFITM3, *t_IFITM3_*, and without IFITM3, *t_o_*, as a function of IFITM3 concentration.

To test this hypothesis, we used continuum modeling of the fusion site^32–34^, based on the theory of lipid membrane elasticity (Figure 3C)^39, 40^, and calculated the mechanical stress and energy barrier of membrane fusion pore expansion. We used the relative repulsion between cholesterol and IFITM3 as the ‘late endosome membrane’ composition of 0.96 k_B_T derived from the MD simulations (Figure 2I) and determined the lipid sorting between the endosome and diaphragm membrane as a function of the IFITM3 mole fraction. We found that cholesterol is enriched in hemifusion diaphragms at 8.3%/(mol_IFITM3_ %) beyond the baseline value of 30%, leading to the reduction of the monolayer spontaneous-curvature by 0.031 nm^- 1^/(mol_IFITM3_ %) (Figure S5A), and an increase of membrane-pore rim line tension by 6 pN/(mol_IFITM3_ %) (Figure S5A). Next, we simulated the fusion site shape (Figure S5B) and the lipid tilt and splay-related stresses in the diaphragm as a function of the IFITM3 mole fraction (Figure S5C). We found that the mean mechanical stress in the diaphragm decreases by 1.28 kBT/(nm^2^·mol_IFITM3_ %) (Figure 3D). The energy barrier for pore expansion increases by 26 k_B_T/(mol_IFITM3_ %) (Figure 3F, sensitivity to cholesterol-IFITM3 relative repulsion presented in Figure S5D and to the cholesterol concentration in Figure S5E). Hence, IFITM3 can inhibit fusion pore expansion due to the IFITM3-cholesterol sorting effect. Considering that cholesterol can undergo a spontaneous flip-flop in the absence of a dedicated flippase^41^, we assume that the cholesterol distribution in the hemifusion diaphragm is symmetric.

A recent study reported that the amphipathic helix domain of IFITM3 induces a strong negative membrane curvature^23^, which may influence the outcome of membrane fusion in addition to the effect of lipid sorting. Thus, we next calculated the changes in the mean mechanical stress and energy barrier of fusion-pore expansion when IFITM3-induced negative membrane curvature alone or combined with IFITM3-induced lipid sorting. Overall, IFITM3- induced negative curvature slightly increased the energy barrier of fusion-pore expansion in the presence of lipid sorting (from 26 to 27 k_B_T/(mol_IFITM3_ %) (Figure 3F). However, in the absence of lipid sorting, IFITM3 had a negligible effect on the energy barrier for pore expansion (Figure 3F). We also tested the possibility that the amphipathic helix domain of IFITM3 penetrates the diaphragm. In such a scenario, IFITM3 cannot induce lipid sorting between the diaphragm and endosomal membrane as it is, on average, equally distributed between the two. We found that the fusion-pore expansion energy barrier increases only by 0.45 k_B_T/(mol_IFITM3_ %) due to the negative intrinsic curvature of the IFITM3 amphipathic helix. Hence, our findings show that IFITM3-induced lipid sorting is the main effector of fusion pore expansion inhibition. This important lipid sorting activity of IFITM3 might be partially caused by the ability of IFITM3 to induce negative membrane curvature^23^.

As IFITM3 increases the energy barrier of fusion pore expansion, the time required to complete viral membrane fusion in late endosomes, hereafter termed ‘dwell time’, increases. We quantified this increase by calculating the ratio between the dwell time in the presence of IFITM3, τ_IFITM3_, to the dwell time without IFITM3, τ_0_ (Figure 3G). Our data showed that even a modest concentration of 0.16% IFITM3 (one IFITM3 molecule per 625 lipids) increases the dwell time by a factor of 33 if we consider both the sorting and direct mechanical effect of IFITM3. In summary, continuum membrane modeling results predict that IFITM3-induced lipid sorting leads to an increased energy barrier for fusion pore formation and, thus, to a prolonged dwell time in hemifusion.

### IFITM3 stabilizes the hemifusion state in late endosomes

To validate our theoretical model, we utilized *in situ* cryo-ET, which allows direct visualization of membrane-protein interactions inside cells in native conditions. A549-IFITM3 cells were infected with a fluorescently labeled virus and were plunge-frozen one hour post-infection (hpi). Using *in situ* cryo-correlative light and electron microscopy (cryo-CLEM)42, infected cells were selected for cryo-focused ion beam (cryo-FIB) milling to prepare cryo-lamellae43,44. Cryo-EM of lamellae showed putative endosome-like organelles that were used as sites for tomography acquisition, and subsequent correlation allowed us to validate the presence of labeled virions in these endosomal compartments (Figure 4A–D and Figure S6). Cryo-ET revealed the ultrastructure of late endosomes (Figure 4E and 4F, Video S1) with a similar multivesicular structure as observed by ET (Figure 1V), showing protein-coated ILVs (Figure S7). In addition to ILVs, we observed a total of 43 individual viral particles in late endosomes and lysosomes of four different cells (Figure 4E, Figure S8–S10, Videos S1–S4), indicating that these viruses are unable to enter the cytoplasm, as IAV membrane fusion should occur within 10 to 15 minutes after uptake in the absence of IFITM31,45,46.

**Figure 4.**
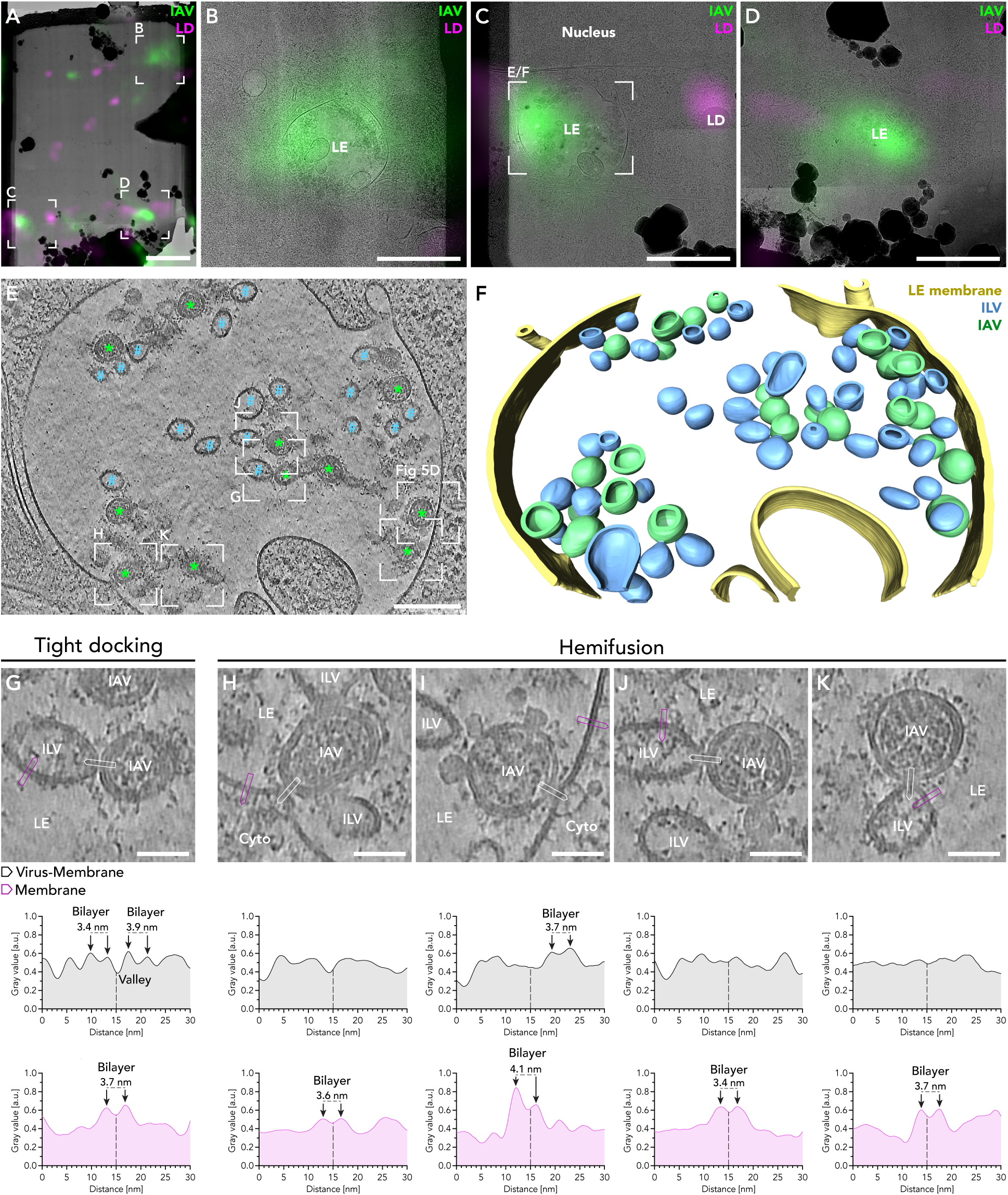
IFITM3 stabilizes hemifusion intermediates during IAV mediated fusion with the late endosomal limiting membrane and ILVs. **(A)** Cryo-lamella of an A549-IFITM3 cell infected with fluorescently labeled influenza virus A/WSN/1933-nDiO (MOI=200, 1 hpi). The fluorescent signal of IAV is shown in green. The LD signal is shown in magenta. **(B–D)** Magnified areas of the correlated cryo-lamella shown in (A). Late endosomes (LE) with a typical multivesicular morphology were found positive for fluorescent IAV signal. One correlated LD is shown in (C). (E) Tomographic slice (5.34 nm thickness) of a late endosome correlated to the IAV fluorescence signal (C). The late endosomal lumen incorporates ILVs (blue hashes) and IAV particles (green asterisks). Reconstructed tomograms were SIRT-like filtered and denoised using cryoCARE. See Video S1 for three-dimensional rendering. **(F)** The three- dimensional volume segmentation of the tomogram is shown in E. The late endosomal membrane is shown in yellow, IAV particles in green, and ILVs in blue. **(G–K)** Magnified areas of reconstructed tomogram in E, showing interaction sites between IAV particles and cellular membranes. For each example, a line profile with thickness of 5.34 nm of the virus-membrane interaction (middle row, black) and a membrane (bottom row, magenta) are shown with positions indicated in the tomograms (white square for virus membrane interaction; magenta square for membrane). The line profile of a tight docking event (G) Line profiles of hemifusion sites between viral and endosomal membranes (H–K) For all shown examples, a line profile of the phospholipid bilayer of an endosomal membrane is shown (magenta plots), with each monolayer discernable. The width of each bilayer was determined by measuring the distance between the two maxima and reported in the figure. See Figure S11 and Figure S12 for a gallery of all contact sites in the tomogram (E) and corresponding line profiles. Scale bars: (A) 3 µm, (B–D) 1 µm, (E, F) 200 nm, (G–K) 50 nm.

Viral particles showed a characteristic diameter of 87.8 nm (SD = 7.8 nm, n = 18). Intriguingly, IAV particles were regularly bound to small crystal-like structures (Figure 4I and 4L, arrowheads), structurally similar to low-density lipoprotein particles containing a cholesterol ester core47 and to structures found in late endosomes and lysosomes48. All viral particles featured a disorganized HA phenotype comparable to *in vitro* cryo-ET studies of IAV particles at pH 4.949. The M1 matrix layer was partially disassembled in 83% of the observed viral particles and fully disassembled in 9%, indicating that the observed viral particles in late endosomes were exposed to a low pH environment. The remaining 8% of viral particles showed no sign of disassembly. Importantly, IAVs were frequently observed in close contact with ILVs and the endosomal limiting membrane. These contact sites displayed deformations of the limiting endosomal membrane towards the viral membrane forming a membrane merger, a structure consistent with hemifusion intermediates between viral and endosomal membranes. We observed 21 contact sites between IAV particles and ILVs and 6 contact sites between IAV particles and the limiting late endosomal membrane (Table S2, Figures S8–S12). We classified 15% of all contact sites as tight docking and 85% as hemifusion (n=27) by linear density profiles (Figures 4G–K, S8–S12). The linear density profiles of the tight docking events showed signal peaks of two juxtaposed phospholipid bilayers separated by a distinct drop in signal, indicating that the two membranes are not in hemifusion. In contrast, linear density profiles of all events classified as hemifusion lacked juxtaposed phospholipid bilayers. The absence of a low-density valley between the viral and endosomal membrane argues for possible lipid exchange between the viral and endosomal membrane, which is in accord with live-cell fluorescence microscopy studies22. Importantly, these structures resembled liposome membrane deformations, including so-called pinching and hemifusion states formed *in vitro* between influenza virions and liposomes36,50–52. However, extended tight docking reported between liposomes and IAV was not observed. It is plausible that some differences between *in situ* and *in vitro* membrane fusion intermediates arise from the proteins present inside the endosomal membranes. Interestingly, viral membrane curvature at the hemifusion sites appeared largely intact, likely reflecting the rigidity and pressure difference between the virus and cell membrane. The M1 layer was still partially or fully intact in most cases (91%), and it might still provide additional rigidity to the virus envelope that favors the initial bent shape of the virus, as commonly seen in virus-liposome fusion *in vitro*50,51,53. No fused IAV or vRNPs were observed in the cytosol, illustrating that IFITM3 can effectively block the membrane fusion of most incoming viruses. Most contact sites (78%, n=27) were observed between viral particles and ILVs (Figure 4I and 4J, Figure S11, Table S2). In contrast, only a smaller fraction (14%, n=6) was found in contact with the limiting late endosomal membrane (Figure 4K and 4L, Figure S12), suggesting that ILVs can frequently serve as a fusion target for IAV. These data show that IFITM3 not only blocks viral entry via fusion with the limiting late endosomal membrane but can also block indirect entry pathways through fusion with ILVs and subsequent back-fusion of ILVs with the limiting late endosomal membrane. Thus IFITM3 can likely impede the endosomal escape of different viruses, which are known to hijack the back- fusion pathway^54^ by blocking membrane fusion with ILVs. This is in accord with a recent study showing that IFITM3 prevents ILV back-fusion55.

### Stabilized hemifusion diaphragms show symmetric geometry and a high energy barrier for fusion pore formation

The hemifusion sites visualized by *in situ* cryo-ET showed an axial symmetry around the central axis, and their geometry could be quantified from our tomograms. Measurements of the hemifusion sites (n=17) (Figure 5A–C) revealed a narrow average outer angle of 63° (SD=12°) and wide inner viral and cytoplasmic angles of 157° (SD=11°) and 140° (SD=13°), respectively. As no significant differences were observed between opposing contact angles, the hemifusion sites were either formed by symmetric expansions or converged into a favorable symmetric geometry56. The average hemifusion diaphragm diameter measured from the phospholipid head groups was 16.5 nm (SD=5.3 nm, n=17), consistent with stable hemifusion diaphragms observed *in vitro*50. The observed hemifusion site geometry allowed us to assess, based on the geometry of the hemifusion diaphragm, the stress in the hemifusion diaphragm and the fusion-pore formation energy as 113 kBT (SD=44 kBT, n=16). This energy barrier is higher than the energy barrier estimated by our simulations (25 kBT with no IFITM3 and 33 kBT with 0.32% IFITM3, Figure 3F). However, the energy barrier generated by our simulations strongly depends on the lipid composition in the late endosome and the monolayer tilt and bending rigidities, so an exact comparison between theory and experiment is challenging. For example, the theory predicts an energy barrier of more than 100 kBT by considering higher cholesterol concentrations (40% instead of 30%, Figure S5E) or a more rigid membrane, as expected for higher cholesterol concentrations (Figure S5F). Therefore, based on the cryo-ET results, it is likely that the cholesterol fraction in the late endosome in our model is underestimated.

**Figure 5.**
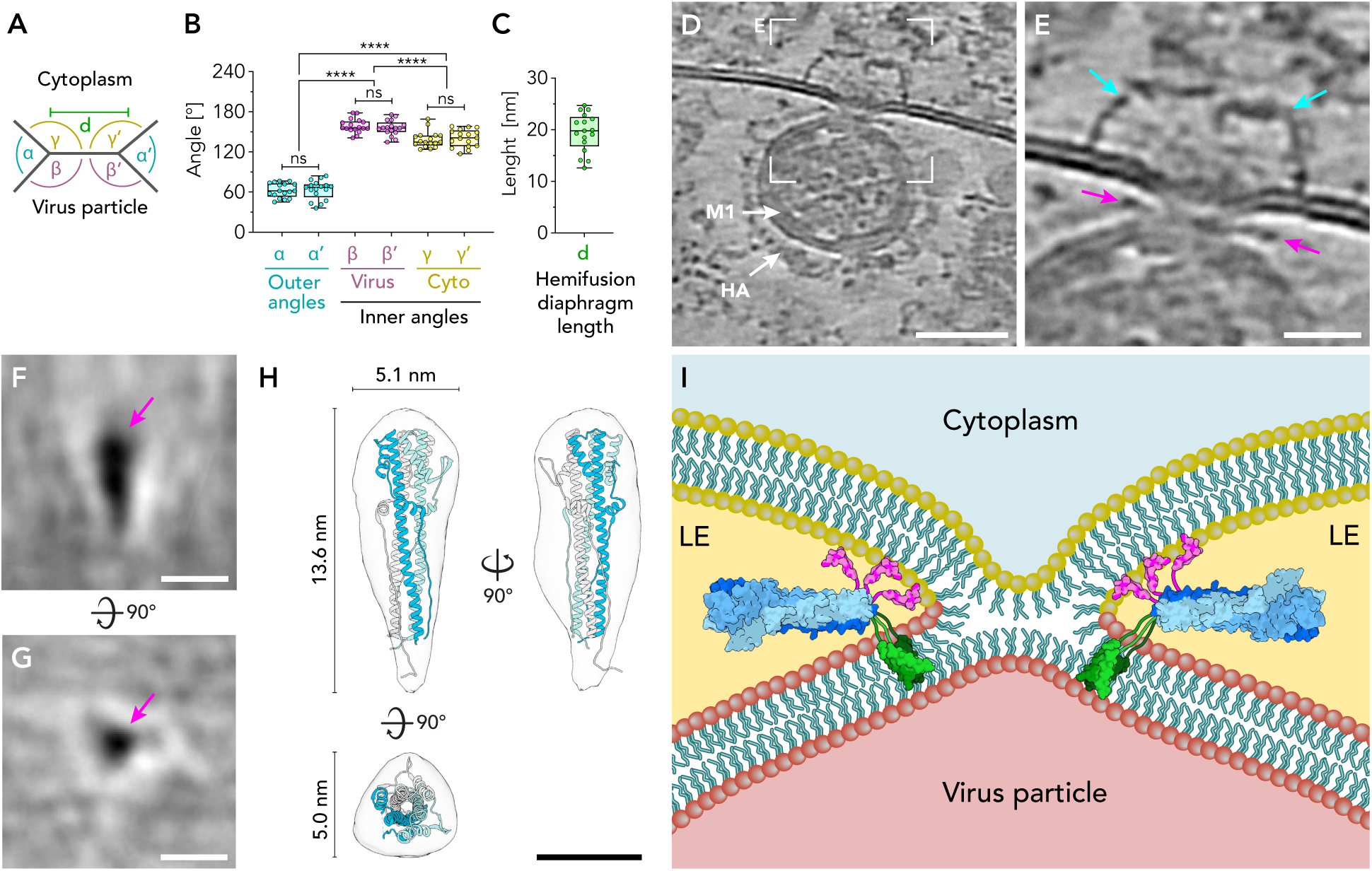
Post-fusion HA glycoproteins are localized at the hemifusion diaphragm. **(A–C)** The geometry of hemifusion diaphragms was analyzed by measurements of angles (B) and diaphragm size (C) for individual hemifusion sites (n = 17). The following angles are defined as indicated in the schematic (A): Outer angles (α and α’), inner angles facing the viral lumen (β and β’), and inner angles facing the cytoplasm (γ and γ’). Paired t-test was performed to analyze the significance of the difference between pairs of angles and between groups of angles. Data is shown as Box and Whiskers plots indicating the median (center of box), 25% and 75% quartiles (bounds of box), minimum and maximum values (bars), and all data points. **(D)** Magnified area of reconstructed tomogram (Figure 4E) showing a stabilized hemifusion site between IAV particle and late endosomal membrane. The image is rotated 90° compared to Figure 4E. **I** Magnified area of the hemifusion diaphragm between the late endosomal membrane and virus particle (D). Electron dense structures are localized orthogonal to the hemifusion diaphragm (magenta arrows), representing post-fusion HA glycoproteins at the diaphragm region. Close to the hemifusion site, two OMDPs were observed (cyan arrow). **(F–H)** Subtomogram average of post-fusion HA localized at the hemifusion diaphragm. Central slices of the subtomogram average for side-view (F) and top-view (G) are shown. The electron-dense post-fusion HA is localized in the center (magenta arrow). The post-fusion HA2 structure (PDB: 1QU1)^71^ was fitted to the three-dimensional isosurface of the subtomogram average (H). See Figure S13 for all iterative steps of the subtomogram average analysis. **(I)** Schematic model of a stabilized hemifusion diaphragm visualizing the membrane configuration and localization of post-fusion HA. Late endosomal phospholipids are shown in yellow and viral phospholipids in red. Lumina of the virus particle, cytoplasm, and late endosome (LE) are indicated. The trimeric post-fusion HA glycoprotein model without fusion peptide (PDB: 1QU1)^71^ is shown in blue, with each monomer shown in a different shade of blue, fusion peptides are shown in magenta (PDB: 1QU1)^71^. The transmembrane domain of HA is shown in green (PDB: 6HJR)^72^. Statistical significance: * for p<0.05, ** for p<0.01, *** for p<0.001 **** for p<0.0001. Scale bars: (D) 50 nm, I 20 nm, (F and G) 10 nm, (H) 5 nm, (I) 10 nm.

### Post-fusion HA is confined in stabilized hemifusion diaphragms

Intriguingly, regular cylindrical densities orthogonal to the hemifusion diaphragms were frequently observed (Figure 5D and 5E, red arrow) at both ILV membranes (Figure S11, red arrows) and late endosomal membranes (Figure S12, red arrows). Subvolume extraction and iterative averaging of potential HA glycoproteins localized at hemifusion diaphragms (n=30) revealed a rod-shaped structure featuring a stalk and head (Figure 5F and 5G, Figure S13, Video S5) with a dimension of 13.6×5.1×5.0 nm, overall consistent with the HA post-fusion conformation4. The crystal structure of the post-fusion HA28 was fitted to the electron density of the subtomogram average (Figure 5E) with high accuracy (97.1% atoms inside the isosurface). Hence, the observed density at the hemifusion diaphragm likely represents the viral HA glycoprotein in its post-fusion state. No densities corresponding to HA1 subunits were evident, presumably due to HA1 dissociation at reducing conditions of endosomes57. In addition to the HA2 density, the average revealed both the endosomal and the viral membranes (Figure S13A, green dotted lines), confirming the high symmetric geometry of the hemifusion sites. On two occasions, the outer membrane dome protein (OMDP), a protein complex previously characterized by us^48^, was observed on the cytosolic site of the hemifusion site (Figure 5D and 5E, cyan arrows). However, since our data show that IFITM3 blocks membrane fusion in the absence of OMDPs, it is unlikely that OMDPs play an important role in the antiviral function of IFITM3.

Hence, IFITM3 does not exclude viral fusion machinery from the fusion sites, and it does not interfere with the conformational transitions of the viral fusion protein, which is consistent with previous data showing that IFITM3 does not alter endosomal pH16. This argues for the role of the IFITM3-lipid sorting activity in stabilizing the hemifusion state and shows that IFITM3 can inhibit a broad variety of viral fusion proteins.

### Influenza A virus membrane fusion in non-inhibitory conditions

We finally aimed to compare the observed stabilized hemifusion sites in A549-IFITM3 cells (Figure 4 and Figure 5) to non-inhibited influenza A virus membrane fusion in wild-type A549 cells. To this aim, we infected A549 cells that do not express IFITM3 (Figure 1A, S1C) with the same infection conditions as the A549-IFITM3 cells (MOI=200, 1 hpi). In a total of 9 tomograms of late endosomal organelles from 2 independent experiments, we could not observe any virus particles, indicating that in A549-IFITM3 cells, the observed hemifusion sites at 1 hpi are indeed stabilized by IFITM3.

To structurally characterize viral entry in A549 cells, we determined the timing of IAV membrane fusion in A549 cells by an infection time course (Figure 6A and B) where NH4Cl was added at different times post-infection. Since NH4Cl rapidly diffuses through membranes and elevates endosomal pH^58^, this approach allows estimating the duration of IAV endocytic uptake till the onset of viral fusion59. The number of infected cells over time followed a sigmoidal curve with a half-time of 45 minutes post-infection (mpi), at which IAV endosomal membrane fusion is completed in 50% of the infected cells.

Based on these data, we structurally characterized IAV membrane fusion in late endosomes by *in situ* cryo-ET for time points between 18 and 31 mpi at a high MOI of 3×104 PFU/cell. In a total of 42 *in situ* cryo-electron tomograms, we captured 473 individual viral particles at different stages of viral entry. As an initial stage, we observed viral particles in early endosomal organelles featuring fully assembled M1 layers and HA spikes in prefusion conformation (40% of virions, n=187), which indicates a neutral pH (Figure 6C and D). In addition, we captured virions harboring unstructured HA and partially disassembled M1 layer, indicating that virions were exposed to low pH in late endosomal compartments (51% of the virions, n=240) (Figure 6E and F). Similar structural changes of HA glycoproteins and M1 layer were previously reported by *in vitro* cryo-ET studies of IAV particles subjected to low pH36,60,61. We found 7% of the virions (n=31) in a hemifusion state (Figure 6G and H, Figure S15). In a few cases (n=5), we found the hemifusion diaphragm to be extended to a length of more than 200 nm (Figure 6I and J), although due to the small number of observations, it remains an open question if hemifusion expansion is a characteristic stage during IAV entry in uninhibited conditions. Finally, we were able to observe post-fusion states for 3% of the viral particles (n=15) at the limiting late endosomal membrane as well as ILVs (Figure 6I and J), with vRNPs already present at the cytoplasm or ILV lumen, whereas the unstructured HA localized at the inner leaflet of the endosomal membrane. Interestingly, we regularly observed patches of assembled M1 layer on the cytoplasmic side of the late endosomal membrane at post-fusion sites, indicating that a fully disassembled M1 layer might not be necessary for IAV membrane fusion. Viral particles in endosomes of A549-IFITM3 cells (Figures 4, 5, S8–S12) showed a similar morphology to the pre-fusion viral particles with unstructured HA in A549 cells (Figure 6E and F), although with the difference of accumulated hemifusion sites both at the limiting endosomal membrane and ILVs. This is particularly striking, as the stabilized hemifusion sites in A549-IFITM3 cells were observed at a much later time point of 1 hpi, where the majority of virus particles are expected to have already fused in non-inhibitory conditions of A549 cells (Figure 6A).

**Figure 6.**
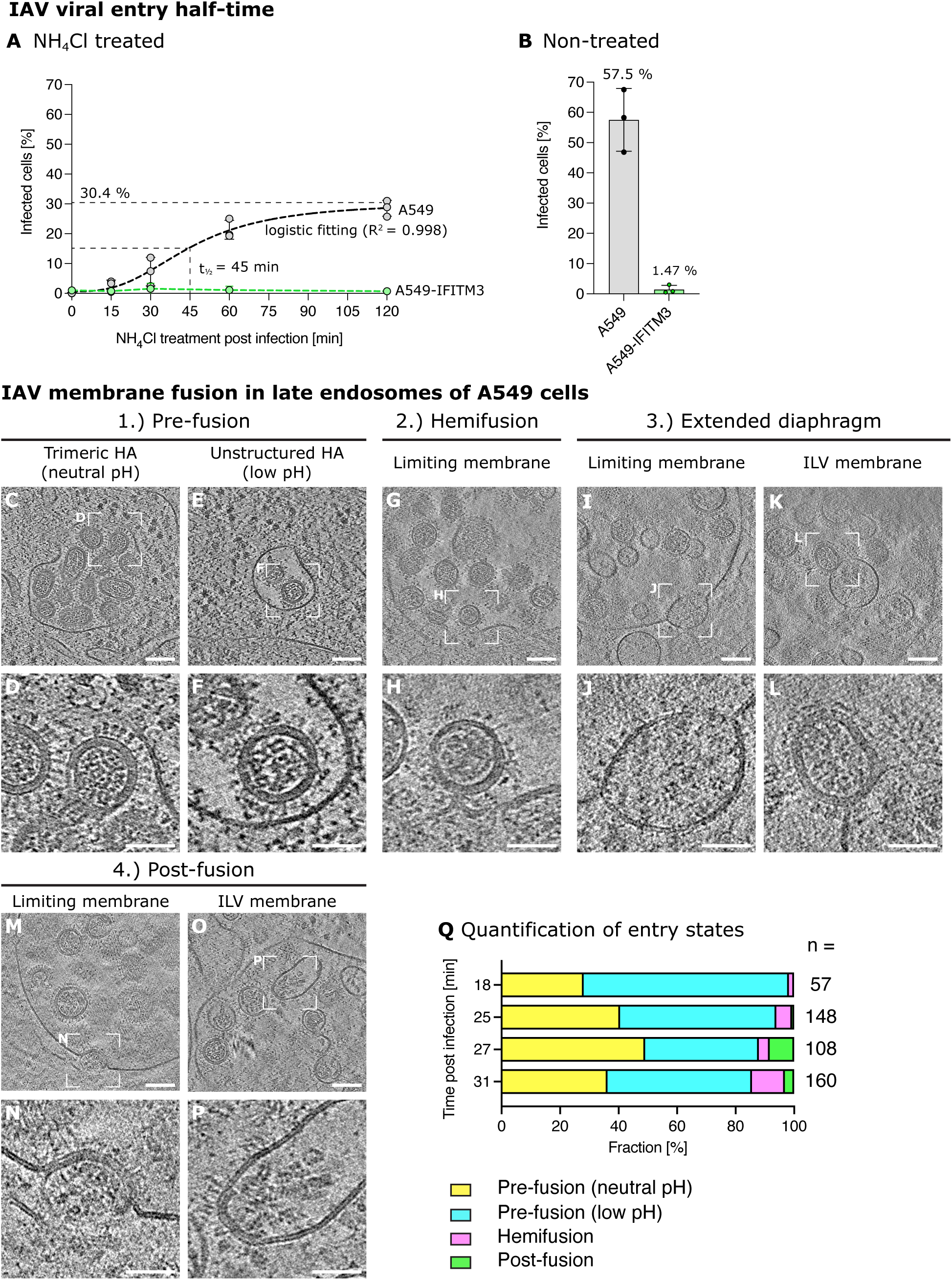
Temporal and structural analysis of IAV cell entry and membrane fusion in A549 cells. **(A and B)** IAV penetration assay by NH4Cl add-in time course. (A) A549 cells (black dotted line) and A549-IFITM3 cells (green dotted line) were infected with A/WSN/1933-PA-mScarlet (MOI = 3). Cells were treated with 50 mM NH4Cl at different time points post-infection (0 mpi, 15 mpi, 30 mpi, 60 mpi, and 120 mpi) and fixed at 12–14 hpi. The percentage of infected cells was determined by fluorescence microscopy (see Figure S14 for a detailed analysis of the individual time points) in two independent experiments. For A549 cells, a four-parameter logistic (4PL) curve was fitted to the data (R^2^ = 0.998), and the half-time for viral penetration was determined to be *t_1/2_*/ = 45 min. Due to the low infection rates in A549-IFITM3 cells, curve fitting and determination of the half-time were not possible. (B) Control infections in the absence of NH4Cl treatment. **(C–P)** Slices of cryo-electron tomograms capturing IAV particles in endosomal organelles at different stages of viral entry. A549 cells were infected with A/WSN/1933-PA-mScarlet (MOI=3×10^4^ PFU/ml) and plunge-frozen between 18 and 31 mpi. See also Figure S15 for a gallery of hemifusion sites. **(Q)** Quantification of entry stages found at different times post-infection. Scale bars: (C, E, G, I, K, M, O) 100 nm, (D, F, H, J, L, N, P) 50 nm.

## DISCUSSION

IFITM3 has evolved as a broad and unspecific host countermeasure against a wide range of viruses that enter the cell via the late endosomal pathway, like IAV, West Nile virus, Chikungunya virus, respiratory syncytial virus^12^, Sindbis and Semliki Forest virus^16^, tick-borne encephalitis virus^62^ as well as filoviruses63. Therefore, it is likely to function through general principles of viral-host membrane fusion inhibition. However, despite its substantial medical importance, the underlying principles were not understood. We showed that the number of ILVs in the late endosomal lumen is not modulated by IFITM3, which contradicts the ‘fusion decoy’ model22. Atomistic MD simulations revealed that cholesterol specifically interacts with the palmitoylated Cys^72^, which was recently identified as essential for the antiviral properties of IFITM321. Furthermore, atomistic MD simulations in lipidic systems mimicking an endosomal membrane composition and binary POPC–cholesterol mixtures show that IFITM3, despite the interaction of cholesterol with palmitoylated Cys^72^, overall repels cholesterol from its vicinity. Cholesterol is being replaced by phospholipids, including LBPA. Interestingly a recent study suggests that LBPA promotes IAV membrane fusion^64^, which further indicates that IFITM3 drives local lipid redistribution to disfavor IAV membrane fusion. Since IFITM3 is excluded from the hemifusion diaphragm due to its transmembrane domain and the structure of the hemifusion diaphragm rim, our MD simulations data indicates that IFITM3 can locally modulate the bilayer lipid composition and thereby withdraw fusion-pore driving lipids from the hemifusion diaphragm. Furthermore, the mechanism by which IFITM3 depletes cholesterol from its vicinity does not depend on a particular amino acid region but is instead due to its overall conformational structure. Since IFITM proteins share a similar three-dimensional structure, it is plausible that the mechanism by which IFITM3 depletes cholesterol could be shared within this family of proteins.

Using membrane model systems, previous studies estimated that while hemifusion takes place within 5–30 seconds, fusion pore formation can occur in the order of many seconds to minutes (20–200 sec)51,64–68. Here conducted continuum membrane modeling suggests that IFITM3-induced lipid sorting increases the energy barrier of fusion pore formation, leading up to 3 orders of magnitude increase in hemifusion dwell time (Figure 3G). Therefore, viruses are trapped in the late endosome and subjected to degradation by proteases and lipases in the endosomal-lysosomal system. Virus degradation is supported by the observation of rod-like structures with a diameter of 13.7 nm consistent with vRNPs presumably released upon viral degradation in late endosomes in the A549-IFITM3 cell line (Figure S8F and S8G). It remains to be seen whether the previously reported IFITM3 clustering in the membrane, which depends on IFITM3 palmitoylation18,19, enhances a lipid sorting effect and thereby also the dwell time.

We directly visualized stabilized hemifusion during viral-induced membrane fusion in late endosomes of IFITM3 overexpressing A549 cells using correlative *in situ* cryo-ET, which revealed IAV arrested at the hemifusion state, hence confirming the previously proposed ‘hemifusion stabilization’ hypothesis^22^ at molecular resolution. In accordance with these data, we could show different IAV entry and fusion events, including post-fusion endosomal escape in A549 wild-type cells in uninhibited conditions. Furthermore, subtomogram averaging revealed post-fusion HA localized close to the hemifusion sites, providing direct evidence that IFITM3 does not affect the fusogenic activity of the viral fusion protein. Since the backfolding of the extended HA intermediate to the coiled-coil post-fusion state is not inhibited, the energy released during this irreversible process was insufficient to overcome the energy barrier for fusion pore formation, supporting the IFITM3-induced lipid-sorting as the principal inhibitory mechanism of IFITM3.

### Limitations of the study

This study provides *in situ* cryo-ET datasets on viral membrane fusion directly inside late endosomes. This implies that membrane configurations might not be identified as clearly as in earlier *in vitro* studies using artificial membranes that often lack proteins. Nevertheless, our data quality is sufficiently good to differentiate tight docking events from hemifusion events unambiguously. Intermediates that occur during the transition between tight docking and hemifusion, however, can currently not be distinguished. To enhance the quality of our sampling, we restricted our full atomistic MD simulations to a single IFITM3 monomer embedded in a lipid bilayer with 300 lipids. By imposing this limitation, we ensured statistically consistent lipid distribution across three independent simulations in all systems. However, investigating the impact of IFITM3 clustering on lipid sorting necessitates a larger membrane patch and significantly longer simulation times to achieve convergence. In the future, such simulations could be performed with coarse-grained MD simulations, which have been shown before to be able to describe the function of transmembrane proteins in specific lipid mixtures69. We limited our continuum modeling to the standard pathway of fusion. Although other off-pathway fusion events, such as so-called leaky-fusion, were reported for protein-free liposomal systems at low cholesterol concentrations (below 15 mol%)50,70. However, we did not observe any indications of such rupture events in our cell experiments. Furthermore, our continuum modeling only allows us to estimate the fold change resulting from IFITM3 presence, but the absolute time needed to complete the fusion reaction cannot be determined from the model.

## ACKNOWLEDGMENTS

We want to thank Frank Orth for the cover art. We acknowledge access to the infrastructure and support provided by the Infectious Diseases Imaging Platform (IDIP) at the Center for Integrative Infectious Disease Research Heidelberg (Germany), the Cryo-EM Network at the Heidelberg University (HD-cryoNet) (Germany), the Electron Microscopy Core Facility at EMBL Heidelberg (Germany), the computational resources at CSC-IT Centre for Science Ltd. (Espoo, Finland) and IT support at BioQuant Heidelberg (Germany). We want to thank Dr. Marco Binder for access to the cDNA library and plasmids for lentiviral production, Prof. Dr. Ervin Fodor for providing plasmids for A/WSN/1933 reverse genetics system, and Prof. Dr. Andrew Mehle for providing plasmids for the fluorescently tagged PA. We want to thank Jana Koch and Dr. Pierre-Yves Lozach for providing the plasmids for Rab7 and LAMP1 and Prof. Stefan Pöhlmann for providing the plasmids for IFITM1 and IFITM2. We are grateful to Hwayoung Lee, Prof. Wonpil Im, and Prof. Howard Hang for providing us with the Cys^72^ palmitoylated IFITM3 initial structure for performing atomistic MD simulations. This work was supported by a research grant from the Chica and Heinz Schaller Foundation (Schaller Research Group Leader Program) to P.C., N.R., R.B, C.L., J.M. and A.K. In addition, this work was funded by the Deutsche Forschungsgemeinschaft (DFG, German Research Foundation) Sonderforschungsbereich (SFB, Collaborative Research Centre) 1129 to Y.S., B.B., U.S.S., P.C., S.P. and S.K; SFB TRR^83^ to F.L. and W.N.; DFG Ni 423/10-1 to D.B. and W.N.; TRR186 to B.B.; DFG-FNR EMSIVILIN to P.C. and M.W.-M.; G.G. acknowledges support by the Minerva Stiftung. The authors gratefully acknowledge the data storage service SDS@hd supported by the Ministry of Science, Research, and the Arts Baden-Württemberg (MWK) and the DFG through grant INST 35/1314-1 FUGG and INST 35/1503-1 FUGG.

## AUTHOR CONTRIBUTIONS

**Steffen Klein:** Conceptualization; Investigation and formal analysis (creating stable cell line and reporter virus, confocal microscopy, BlaM assay, immunoblotting, infection assay, NH4Cl add-in assay, colocalization assay, ET, cryo-CLEM, *in situ* cryo-ET, STA); Visualization; Writing – Original Draft. **Gonen Golani:** Investigation and formal analysis (membrane modeling and theory, cholesterol affinity and repulsion model, energy barrier calculation from cryo-ET data); Writing – Original Draft. **Fabio Lolicato**: Investigation and formal analysis (MD simulations design, computational resources, data curation, methodology); Writing – Original Draft. **Carmen Lahr:** Investigation (creation of reporter virus, infection assay, NH4Cl add-in assay, viral entry assay, western blot analysis). **Daniel Beyer:** Investigation and formal analysis (systems preparation and equilibration for MD simulations). **Alexia Herrmann:** Investigation and formal analysis (cholesterol labeling and IP). **Moritz Wachsmuth-Melm:** Formal analysis (cryo-ET). **Nina Reddmann:** Investigation (creation of stable cell line and reporter virus). **Romy Brecht:** Investigation (creation of stable cell line, BlaM assay). **Mehdi Hosseinzadeh:** Formal analysis (ET). **Androniki Kolovou:** Investigation (HPF/FS sample preparation). **Jana Makroczyova:** Investigation (immunoblotting, immunofluorescence). **Sarah Peterl**: Investigation (Cryo-EM sample preparation). **Martin Schorb:** Investigation (ET data acquisition). **Yannick Schwab:** Writing – Review & Editing. **Britta Brügger:** Writing – Review & Editing. **Walter Nickel:** Writing – Review & Editing. **Ulrich Schwarz:** Writing – Review & Editing. **Petr Chlanda:** Conceptualization; Investigation and formal analysis (*in situ* cryo-ET); Funding acquisition; Supervision; Writing – Original Draft; Writing – Review & Editing.

## METHODS

### Experimental model and subject details

#### Human embryonic kidney 293 cells expressing SV40 large T antigen (HEK293T)

HEK293T cells were obtained from the American Type Culture Collection (ATCC). Cells were cultured at 37 °C in 5% CO_2_ in Dulbecco’s modified Eagle medium (DMEM) supplemented with GlutaMAX™-I, 10% fetal bovine serum (FBS), and 1% penicillin/streptomycin (P/S).

#### HEK293T-Master Cell Bank cells (HEK293T-MCB)

HEK293T-MCB cells were kindly provided by Dr. Marco Binder (DKFZ, Heidelberg, Germany). Cells were cultured at 37 °C in 5% CO_2_ in DMEM supplemented with GlutaMAX™-I, 10% FBS, and 1% P/S.

#### Madin-Darby canine kidney cells (MDCK)

MDCK cells were kindly provided by Prof. João Amorim (Instituto Gulbenkian de Ciência, Portugal). Cells were cultured at 37 °C in 5% CO_2_ in DMEM-F12 supplemented with 10% FBS and 1% P/S.

#### A549 cells

A549 is a cell line derived from carcinomatous lung tissue from a Caucasian male at age 5875 and is regularly used as a model cell line for IAV infection studies. Cells were obtained from ATCC and cultured at 37 °C in 5% CO_2_ in DMEM-F12 supplemented with 10% FBS and 1% P/S.

#### A549 cells expressing IFITM3 (A549-IFITM3)

To study the antiviral effect of IFITM3, we established an A549 cell line stably overexpressing IFITM3 using lentiviral transduction, as described in the following section.

##### Cloning

The IFITM3 gene sequence, including a stop codon, was transferred from a cDNA entry vector (pENTR221-clone3728-IFITM3) to the destination vector pWPI-IRES-Puro. 150 ng of entry and destination vector were added with 1 µl Gateway™ LR Clonase Enzyme Mix (ThermoFisher Scientific, Invitrogen) in a total volume of 5 µl TE buffer (10 mM Tris, 0.1 mM EDTA, pH 8). The reaction was run for 2 hours (h) at 25 °C on a thermocycler.

##### Transformation

50 µl *E. coli* Stellar™ competent cells (Takara) were combined with 5 µl reaction mix from the cloning step and incubated for 30 min on ice and subsequently heat shocked for 45 sec at 42 °C on a heating block. 450 µl S.O.C. medium (ThermoFisher Scientific, Invitrogen) was added, cells were transferred to a 14 ml round bottom tube and incubated for 1 h at 37 °C on a shaker with 180 rpm. Cells were plated on 1.5% LB agar plates with 50 µg/ml ampicillin and incubated overnight at 37 °C. Individual colonies were picked, added to 50 ml LB media containing 50 µg/ml ampicillin, and incubated for 16 h at 37 °C on a shaker with 180 rpm. Plasmids were purified using the QIAprep Spin Miniprep Kit (QIAGEN), and individual clones were sequenced at Microsynth using the IRES-rev primer (TATAGACAAACGCACACCG). The correct insert was validated, and a second 200 ml overnight culture was purified using the Plasmid Maxi Kit (QIAGEN). The new plasmid is named pWPI- IFITM3.

##### Lentiviral production

All work with infectious lentiviruses was performed under biosafety level 2 (BSL-2) laboratory conditions. Lentivirus was produced using a second-generation packaging system utilizing the envelope plasmid pCMV-VSV-G, the packaging plasmid psPAX2, and the previously cloned transfer plasmid pWPI-IFITM3. For enhanced protein expression, we further used the pAdVantage™ plasmid (Promega). HEK293T-MCB cells were seeded in a 6-well plate with 4 × 105 cells per well in a 2.5 ml cell culture medium (DMEM- GlutaMAX™-I, 10% FBS, 1% P/S) and incubated overnight at 37 °C with 5% CO2. The transfection mix was prepared as follows for one well: 0.375 µg pCMV-VSV-G plasmid, 0.75 µg psPAX2 plasmid, 1.25 µg pWPI-IFITM3 plasmid, 0.125 µg pAdVantage™ plasmid were added in 250 µl Opti-MEM medium (ThermoFisher Scientific, Gibco). 7.5 µl transfection reagent TransIT-2020 (Mirus Bio) was added and incubated for 30 min at RT. The transfection mix was added to one well of the 6-well plate. 6 h post-transfection, the cell culture medium was replaced with 1.5 ml fresh medium. 48 h post-transfection, lentivirus containing supernatant was harvested and filtered through a 0.45 µm CME filter (Roth). A Lenti-X™ GoStix (Takara) antigen test was used to verify successful lentivirus production.

##### Transduction

A549 cells were seeded in a 12-well plate with 2 × 104 cells per well in 1 ml cell culture medium (DMEM-F12, 10% FBS, 1% P/S) and incubated overnight at 37 °C with 5% CO2. 350 µl filtered lentivirus supernatant was supplemented with polybrene infection reagent (Merck, Sigma-Aldrich) in a final concentration of 5 µg/ml. The cell culture medium of A549 cells was removed, and 350 µl lentivirus supernatant was added to the cells. 2 days post-transduction, the antibiotic selection was started with 0.8 µg/ml puromycin. After 2 days of selection, the puromycin concentration was increased to 1.2 µg/ml. The selection was continued for 2 more days until all non-transduced control cells detached from the dish. Selected cells were trypsinized and transferred to a 10 cm cell culture dish and were further grown in a selection medium with 1.5 µg/ml puromycin. Individual cell colonies could be observed after 2 more days. Individual colonies were separated using cloning cylinders (Sigma-Aldrich) and further expanded to establish the monoclonal cell line A549-IFITM3.

#### Influenza A virus (strain A/WSN/1933)

##### Reverse genetics

All work with infectious IAV was performed under BSL-2 laboratory conditions. The IAV strain A/WSN/1933 (H1N1) was rescued from a reverse genetics system established by Hoffmann et al.76. 4 × 106 HEK293T cells were seeded in a 10 cm cell culture dish and grown overnight in 10 ml complete growth media (DMEM-GlutaMAX™-I, 10% FBS, 1% P/S). The transfection mix was prepared as follows: 2.5 µg of each of the 8 plasmids of the reverse genetics system (pHW2000-PB1-WSN, pHW2000-PB2-WSN, pHW2000-PA-WSN, pHW2000-NP-WSN, pHW2000-NA-WSN, pHW2000-M-WSN, pHW2000-NS-WSN, pHW2000-HA-WSN) were added to a total volume of 2 ml Opti-MEM medium. 60 µl transfection reagent TransIT-293 (Mirus Bio) was added and incubated for 30 min at RT and added to the 10 cm dish with HEK293T cells. After 24 h, the medium was exchanged with FBS-free infection medium (DMEM-GlutaMAX™-I, 1% P/S, 0.3% BSA, 2 µg/ml TPCK-trypsin), and 4 × 106 MDCK cells were seeded in the same cell culture dish to be co-cultured with the transfected HEK293T cells. After 24 h, virus-containing supernatant (P0) was harvested and centrifuged for 10 min with 1,000 × g to remove cell debris. The supernatant was aliquoted, shock frozen in LN2, and stored at −80 °C until further use.

##### Virus propagation and purification

6 × 106 MDCK cells were seeded in 30 ml complete growth media (DMEM-GlutaMAX™-I, 10% FBS, 1% P/S) in T175 cell culture flasks and grown overnight. The virus-containing supernatant (P0), recovered from the reverse genetics system, was diluted 1:10 in FBS-free infection media (DMEM-GlutaMAX™-I, 1% P/S, 0.3% BSA, 2 µg/ml TPCK-trypsin). MDCK cells were washed 3 times with PBS. 10 ml of the diluted virus supernatant was added to the cells and incubated for 1 h at 37 °C with 5% CO2. Cells were washed 1 time with PBS, and 40 ml FBS-free infection medium was added. Cells were grown for 3 days until a cytopathic effect was observed. The virus-containing supernatant (P1) was centrifuged 2 times for 15 min at 2,200 × g to remove cell debris. Cleared supernatant was subsequently sucrose purified: 33 ml of virus-containing supernatant was overlaid on 5 ml of 30% sucrose in HEN buffer (10 mM HEPES, 1 mM EDTA, 100 mM NaCl, pH 7.4) in a thin-walled centrifugation tube and centrifuged for 90 min at 83,018 × g, 4 °C using the swing-out rotor SW32 on an Optima L-90K (Beckman Coulter) ultracentrifuge. The supernatant was discarded, and the pellet was resuspended in 1 ml HEN buffer. The sample was centrifuged again for 30 min at 15,728 × g, 4 °C using the fixed-angle rotor TLA-120.2 on an Optima TLX (Beckman Coulter) ultracentrifuge. The supernatant was discarded, and the pellet was dissolved in 200 µl HEN buffer. The purified virus was aliquoted, shock frozen in LN2, and stored at −80 °C until further use.

##### Plaque assay

1 × 106 MDCK cells per well were seeded in a 6-well plate in 2 ml complete growth medium (DMEM-F12, 10% FBS, 1% P/S) and grown overnight. Dilution series of purified virus stock was prepared in an FBS-free infection medium (DMEM-GlutaMAX™-I, 1% P/S, 0.3% BSA, 2 µg/ml TPCK-trypsin) from 10-3 to 10-9. Cells were washed 3 times with PBS, and 800 µl of each virus dilution was added per well. Cells were incubated for 1 h at 37 °C and 5% CO2, virus dilution was removed, cells were washed 2 times with infection medium and overlayed with 3 ml plaque medium containing Avicel (DMEM-GlutaMAX™-I, 0.5% P/S, 0.15% BSA, 1 µg/ml TPCK-trypsin, 1.2 % Avicel RC-581) per well. After 2 days, the plaque medium was aspirated, and cells were washed 2 times with PBS and fixed with 1% glutaraldehyde (GA) in PBS for 30 min at RT. Cells were washed with PBS and stained with 1% crystal violet solution (Sigma-Aldrich) for 10 min at RT. Cells were thoroughly washed with water, and plaques were counted. For each dilution, virus titer, *T_virus_*, was determined by with the number of plaques, *n_plaque_*, the dilution factor, *d*, and the volume of virus dilution, *V*. The average titer of all wells containing plaques was determined and used as final virus titer.

#### Fluorescently labeled influenza A virus (A/WSN/1933-nDiO)

For *in situ* cryo-CLEM, we fluorescently labeled A/WSN/1933 IAV using the fluorescent and lipophilic membrane dye neuro-DiO (nDiO). The virus was recovered from a reverse genetics system and purified as described above. 5 µl of ready to use nDiO solution ‘Cellbrite™ Green’ (Biotium) was added to 200 µl purified virus and incubated for 1 h on a rotation wheel at RT. The virus was subsequently purified again by sucrose purification: 200 µl purified and fluorescently labeled virus was overlaid on 500 µl of 10% sucrose in HEN buffer in a thick-walled centrifugation tube and centrifuged for 90 min at 107,000 × g, 4 °C using the swing-out rotor TLS-55 on an Optima Max (Beckman Coulter) ultracentrifuge. The supernatant was discarded, and the pellet was resuspended in 1 ml HEN buffer. The sample was centrifuged again for 30 min at 15.728 × g, 4 °C using the fixed-angle rotor TLA-120.2 on an Optima TLX (Beckman Coulter) ultracentrifuge. The supernatant was discarded, and the pellet was dissolved in 200 µl HEN buffer. The fluorescently labeled virus was aliquoted, shock frozen in LN2, and stored at −80 °C until further use. Viral titer was determined again using the plaque assay as described above.

#### Fluorescent influenza a reporter virus (A/WSN/1933-PA-mScarlet)

For the infection assay, we established a reporter virus A/WSN/1933 expressing a fluorescently tagged polymerase acidic protein (PA) based on Tran et al.77. For increased integration stability of the fluorescent tag in the viral genome, we performed codon optimization of the gene sequence coding for the fluorescent protein mScarlet by substitution of CpG and TpA sites, which are known to be recognized by the Zinc finger antiviral protein restriction factor78.

##### mScarlet codon optimization

The gene sequence coding for mScarlet^79^ was codon-optimized for mammalian gene expression using the GenSmart™ codon optimization algorithm (version 1.0, GenScripts). Subsequently, all CpG and TpA sites in the codon-optimized sequence were manually exchanged without modifying the aminoacidic sequence. This optimized sequence was synthesized and inserted in a pcDNA3.4 plasmid by the GeneArt gene synthesis service (ThermoFisher Scientific). The newly synthesized plasmid is named pcDNA3.4-mScarlet- codon-optimized.

##### Cloning

Using In-Fusion cloning (Takara), we exchanged the mScarlet of the reverse genetics’ plasmid pHW2000-PA-mScarlet (kindly provided by Prof. Dr. Andrew Mehle) with the codon-optimized mScarlet, generated as described above. Overhang PCR was prepared as follows: 1 ng pcDNA3.4-mScarlet-codon-optimized plasmid, 0.2 µM forward overhang primer (CCCACGCCCTGCGCGCGGCAGCAATGGTGTCCAAGGGTGAAGC), 0.2 µM reverse overhang primer (AAGCAGTTTTCTAGATCACTTGTACAGCTCATCCATTCCAC), 12.5 µl

CloneAmp™ HiFi PCR Premix (Takara) and 1 µl DMSO were mixed in a total volume of 25 µl H2O. PCR reaction was run for 35 cycles with denaturation at 98 °C for 10 sec, annealing at 64 °C for 15 sec, and elongation at 72 °C for 10 sec. 8 µl Cloning Enhancer (Takara) was added to 20 µl of the PCR product and incubated for 15 min at 37 °C on a thermocycler, followed by an inactivation step at 80 °C for 15 min. The pHW2000-PA-mScarlet plasmid was linearized by restriction digestion: 1 µg pHW2000-PA-mScarlet plasmid, 1 u XbaI (New England Biolabs), 1.33 u BssHII (New England Biolabs) were added in a total of 50 µl 1× CutSmart buffer (New England Biolabs). The reaction was run for 15 min at 37 °C on a thermocycler, followed by an inactivation step at 65 °C for 20 min. Restriction digestion products were separated on a 0.7% agarose gel, and linearized plasmid was purified from the gel using the NucleoSpin Gel and PCR Clean-up kit (Macherey-Nagel). In-Fusion reaction was prepared as follows: 25 ng PCR fragment with overhang, 50 ng linearized and purified vector, 1 µl 5× In-Fusion Enzyme Mix (Takara) were mixed in a total of 5 µl H2O. The reaction was run for 15 min at 50 °C on a thermocycler.

##### Transformation

50 µl *E. coli* Stellar™ competent cells (Takara) were combined with 5 µl In- Fusion reaction product and incubated for 30 min on ice and subsequently heat shocked for 45 sec at 42 °C on a heating block. 450 µl S.O.C. medium (ThermoFisher Scientific, Invitrogen) was added, cells were transferred to a 14 ml round bottom tube and incubated for 1 h at 37 °C on a shaker with 180 rpm. Cells were plated on 1.5% LB agar plates with 50 µg/ml ampicillin and incubated overnight at 37 °C. Individual colonies were picked, added to 50 ml LB media containing 50 µg/ml ampicillin, and incubated for 16 h at 37 °C on a shaker with 180 rpm. Plasmids were purified using the QIAprep Spin Miniprep Kit (QIAGEN), and individual clones were sequenced at Microsynth using a forward primer (GGCAAACAACAGATGGCTGGCAAC) and reverse primer (GTATGCATCTCCACAACTAGAAGG).

The correct sequence was validated, and a second 200 ml overnight culture was purified using the Plasmid Maxi Kit (QIAGEN). The new plasmid is named pHW2000-PA-WSN-mScarlet- codon-optimized.

##### Reverse genetics

A fluorescent reporter virus was recovered and purified as described above for the A/WSN/1933 virus using the reverse genetics system established by Hoffmann et al.76, but here the plasmid pHW2000-PA-WSN was exchanged with pHW2000-PA-WSN-mScarlet- codon-optimized.

### Methods details

#### Immunoblot analysis

The expression levels of IFITM3 in A549 cells, with and without IFNβ-1b treatment, and of the established stable cell line A549-IFITM3 (as described above) were analyzed by immunoblot analysis as described below.

##### Protein extraction

1 × 106 A549 or A549-IFITM3 cells were seeded in 10 cm cell culture dishes in complete growth medium (DMEM-F12, 10% FBS, 1% P/S). A549 cells were treated with different concentrations of IFNβ-1b (0, 100, 500, 1,000, 2,000 units/ml). After 48 h, cells were washed 2 times with ice-cold PBS and lysed with 650 µl RIPA buffer (50 mM Tris-HCL, 150 mM NaCl, 1% v/v Triton X-100, 0.5% v/v sodium deoxycholate, 0.1% v/v SDS, 1× protease inhibitor cocktail (ThermoFisher Scientific, Roche) in H2O) on ice for 20 min. The cell lysate was centrifuged with 12,000 × g for 20 min at 4 °C to remove cell debris. The protein concentration of the supernatant was quantified using the Pierce™ BCA Protein Assay Kit (ThermoFisher Scientific).

##### Gel electrophoresis

All samples were diluted in RIPA buffer to a final protein concentration of 0.7 µg/µl. 66.7 µl 4× Laemmli sample buffer (Bio-Rad), supplemented with 0.2 M dithiothreitol, was added to 200 µl of diluted sample and incubated for 8 min at 90 °C on a heating block. 40 µl of each sample was loaded to a 4 – 15% precast polyacrylamide gel (Bio-Rad). As a size marker, 10 µl of a broad range color pre-stained protein standard (New England BioLabs) was added to the outer lanes. Samples were separated by gel electrophoresis in 1× TGS running buffer (Bio-Rad) for 60 min with 120 V using a Mini-PROTEAN electrophoresis chamber (Bio-Rad).

##### Western blotting

Separated proteins were transferred from the polyacrylamide gel to a 0.2 µm PVDF membrane (Bio-Rad) for 7 min with a constant 2.5 A using a Trans-Blot Turbo transfer system (Bio-Rad).

##### Immunolabeling

PVDF membrane was cut at 26 kDa to analyze the expression of IFITM3 and actin on the same membrane. Membranes were washed 3 times for 5 min in TBS supplemented with 0.1% v/v Tween-20 (TBS-T) and subsequently blocked in 5% BSA in TBS- T for 1 h at RT on a rocker. The lower part of the membrane (0 – 26 kDa) was incubated in primary antibody against IFITM3 (Proteintech) with a dilution of 1:2,000 in TBS-T for 1 h at RT on a rocker. The upper part of the membrane (26 – 250 kDa) was incubated in primary antibody against actin (Sigma-Aldrich) with a dilution of 1:4,000 in TBS-T for 1 h at RT on a rocker. Membranes were washed 3 times for 5 min in TBS-T and incubated with the secondary antibodies: anti-rabbit-HRP (Santa-Cruz) with a dilution of 1:1,000 in TBS-T for the lower part of the membrane (0 – 26 kDa), immunolabelled against IFITM3 or anti-mouse-HRP (Santa-Cruz) with a dilution of 1:1,000 in TBS-T for the upper part of the membrane (26 – 250 kDa), immunolabelled against actin, respectively. Membranes were washed 3 times for 5 min with TBS and were subsequently incubated with Clarity Western ECL substrate working solution (Bio-Rad) for 5 min. Chemiluminescent signal was acquired using the Azure 400 imaging system (Azure Biosystems).

##### Quantification of protein expression

Individual bands of IFITM3 and actin of each sample were quantified using the Gel Analyzer plugin of FIJI80. IFITM3 signal was normalized to the corresponding actin, and relative IFITM3 expression levels were calculated.

#### Confocal fluorescent light microscopy

Confocal fluorescent light microscopy analysis was performed to study the impact of IFNβ-1b treatment or IFITM3 overexpression on the number and volume of IFITM3 positive organelles in A549 cells.

##### Immunofluorescence

5 × 104 A549 or A549-IFITM3 cells were seeded on 12 mm microscopy coverslips NO. 1 (Marienfeld) in a 6-well plate in complete growth medium (DMEM-F12, 10% FBS, 1% P/S). The next day, A549 cells were treated with 1 × 103 units/ml IFNβ-1b (Immuno Tools). After 24 h IFNβ-1b treatment, cells were fixed with 4% PFA in PBS for 15 min and washed 3 times with PBS. PFA was quenched by incubating the cells with 20 mM glycine in PBS for 10 min, followed by 3 washing steps with PBS for 5 min each. Cells were permeabilized by incubation with 0.2% Triton X-100 in PBS for 5 min, followed by 3 washing steps with PBS for 5 min each. Cells were incubated for 1 h with blocking buffer (3% BSA in PBS supplemented with 0.1% Tween-20 (PBS-T)). After 3 short washing steps with PBS-T, cells were incubated for 1 h with primary antibody against IFITM3 (Proteintech) using an antibody dilution of 1:200 in dilution buffer (1% BSA in PBS-T) at RT on a rocker. After 3 washing steps with PBS-T for 5 min each, cells were incubated with the secondary antibody ‘goat anti-Rabbit Alexa Fluor™ 633’ (ThermoFisher Scientific, Invitrogen) in a dilution of 1:2,000 in dilution buffer for 1 h at RT on a rocker. After 3 washing steps with PBS-T for 5 min each, nuclei were fluorescently labeled by incubation with 1 µg/µl Hoechst 33342 (Sigma-Aldrich) in PBS for 5 min. Cells were washed 3 times with PBS for 5 min each, followed by a short washing step with deionized water. Coverslips were mounted on microscopy slides using 7 µl ProLong Glass Antifade mounting medium (ThermoFisher Scientific, Invitrogen). The mounting medium was cured for 24 h at RT.

##### Confocal microscopy

Confocal microscopy data was acquired using the SP8 TCS laser scanning confocal microscope (Leica) equipped with a 63×/1.4 HC PL APO CS2 oil immersion objective using a UV laser with an excitation wavelength of 405 nm (for Hoechst 33342) and a helium-neon laser with an excitation wavelength of 633 nm (for Alexa Fluor™ 633) in sequential acquisition mode with 4× line accumulation. A pixel size of 60.13 nm was used, and Z-stacks were acquired using a Z-spacing of 200 nm.

##### Deconvolution

Image stacks were deconvolved with AutoQuant X3 (Media Cybernetics) using a theoretical and adaptive point spread function (PSF) for 10 iterations with the following parameters: lens immersion refractive index 1.515; sample embedding refractive index 1.52; sample distance from coverslip of 0 nm; emission wavelength of 461 nm (for Hoechst 33342) or 647 nm (for Alexa Fluor™ 633) and appropriate settings for the used objective (NA 1.4; objective lens magnification 63×).

##### Segmentation and quantification

The IFITM3 signal of the deconvolved image stacks was segmented in 3 dimensions using the surface segmentation algorithm of Imaris (Verison 9.8.2, Oxford Instruments). The number of segmented objects per cell and the volume distribution of all objects were quantified.

#### Colocalization analysis

To evaluate the colocalization between IFITM3 and Rab7 or LAMP1 in the A549-IFITM3 cell line, we performed immunofluorescence, confocal microscopy and colocalization analysis as described below.

##### Transfection

1 × 105 A549-IFITM3 cells were seeded on 12 mm microscopy coverslips NO. 1 (Marienfeld) in a 6-well plate in complete growth medium (DMEM-F12, 10% FBS, 1% P/S). The next day, cells were transfected with either pC1-Rab7-eGFP or pN1-LAMP1-eGFP. A transfection mix was prepared by adding 2.5 µg plasmid and 7.5 µl transfection reagent TransIT-LT1 (Mirus) in 250 µl OptiMEM medium. Transfection mix was added to the cells (250 µl per well), and incubated for 30 min at RT. Cells were incubated at 37°C, 5% CO_2_ for 24 h.

##### Immunofluorescence

Cells were fixed with 4% PFA in PBS for 30 min and washed 3 times with PBS. Cells were permeabilized by incubation with 0.2% Triton X-100 in PBS for 5 min, followed by 3 washing steps with PBS for 5 min each. Cells were incubated for 1 h with blocking buffer (3% BSA in PBS supplemented with 0.1% Tween-20 (PBS-T)). After 3 short washing steps with PBS-T, cells were incubated for 1 h with primary antibody against IFITM3 (Proteintech) using an antibody dilution of 1:200 in dilution buffer (1% BSA in PBS-T) at RT on a rocker. After 3 washing steps with PBS-T for 5 min each, cells were incubated with the secondary antibody ‘goat anti-Rabbit Alexa Fluor™ 633’ (ThermoFisher Scientific, Invitrogen) in a dilution of 1:2,000 in dilution buffer for 1 h at RT on a rocker. After 3 washing steps with PBS-T for 5 min each, nuclei were fluorescently labeled by incubation with 1 µg/µl DAPI in PBS for 1 min. Cells were washed 3 times with PBS for 5 min each, followed by a short washing step with deionized water. Coverslips were mounted on microscopy slides using 7 µl ProLong Glass Antifade mounting medium (ThermoFisher Scientific, Invitrogen). The mounting medium was cured for 24 h at RT.

##### Confocal microscopy

Confocal microscopy data was acquired using the SP8 TCS laser scanning confocal microscope (Leica) equipped with a 63×/1.4 HC PL APO CS2 oil immersion objective using a UV laser with an excitation wavelength of 405 nm (for DAPI), an argon laser with an excitation wavelength of 488 nm (for eGFP) and a helium-neon laser with an excitation wavelength of 633 nm (for Alexa Fluor™ 633) in sequential acquisition mode with 4× line accumulation. A pixel size of 72.1 nm was used, and Z-stacks were acquired using a Z-spacing of 200 nm.

##### Deconvolution

Image stacks were deconvolved with AutoQuant X3 (Media Cybernetics) using a theoretical and adaptive point spread function (PSF) for 10 iterations with the following parameters: lens immersion refractive index 1.515; sample embedding refractive index 1.52; sample distance from coverslip of 0 nm; emission wavelength of 461 nm (for DAPI), 507 nm (for eGFP) or 647 nm (for Alexa Fluor™ 633) and appropriate settings for the used objective (NA 1.4; objective lens magnification 63×).

##### Colocalization analysis

Colocalization analysis between IFITM3 and Rab7, LAMP1 or DAPI was performed by calculating the Pearson’s and Manders’ correlation coefficients^81^ using the ‘coloc2’ tool in ImageJ/FIJI^80^ using a manually selected region of interest for each analyzed cell. Before correlation analysis, the background signal was substracted using a rolling ball algorithm (radius = 30 px)82 implemented in ImageJ/FIJI80.

#### Infection assay using wide-field fluorescent light microscopy

To analyze the antiviral effect of stable overexpression of IFITM3 in A549 cells, we performed an infection assay using the here established reporter virus A/WSN/1933-PA-mScarlet (see section above) as described below:

##### Viral infection

5 × 104 A549 or A549-IFITM3 cells were seeded on 12 mm microscopy coverslips NO. 1 (Marienfeld) in a 24-well plate in complete growth medium (DMEM-F12, 10% FBS, 1% P/S). The next day, the cells were infected with a fluorescent report virus A/WSN/1933-PA-mScarlet using an MOI of 3 in serum free DMEM-F12 medium for 1 h at 37°C, 5% CO2. Cells were washed 3 times with a complete growth medium and incubated at 37°C, 5% CO2. Cells were fixed at 24 hpi with 4% PFA in PBS for 30 min at RT. After fixation, cells were washed 3 times with PBS.

##### Immunolabeling

Cells were permeabilized by incubation with 0.2% Triton X-100 in PBS for 5 min, followed by 3 washing steps with PBS for 5 min each. Cells were incubated for 1 h with a blocking buffer (3% BSA in PBS supplemented with 0.1% PBS-T). After 3 short washing steps with PBS-T, cells were incubated for 1 h with primary antibody against M2 (Proteintech) using an antibody dilution of 1:50 in dilution buffer (1% BSA in PBS-T) at RT on a rocker. After 3 washing steps with PBS-T for 5 min each, cells were incubated with DAPI using a 1:1000 dilution (Sigma-Aldrich) and the secondary antibody ‘goat anti-Mouse Alexa Fluor™ 633’ (ThermoFisher Scientific, Invitrogen) in a dilution of 1,000 in dilution buffer for 1 h at RT on a rocker. Cells were washed 3 times with PBS for 5 min each, followed by a short washing step with deionized water. Coverslips were mounted on microscopy slides using 7 µl ProLong Glass Antifade mounting medium (ThermoFisher Scientific, Invitrogen). The mounting medium was cured for 24 h at RT.

##### Automated high-throughput fluorescent imaging

Wide field microscopy data was acquired using the Celldiscoverer 7 automated microscope (Zeiss) equipped with a 20×/0.95 NA PL APO COR objective and Axiocam 712 camera.

##### Quantification

Stitched composite images were binned 6 times in ImageJ/FIJI and nuclei were segmented using the StarDist plugin in ImageJ/FIJI83. The average signal intensity in the segmented area of each individual nucleus was calculated for M1 and PA-mScarlet. Maximum average signal intensity for M2 and PA-mScarlet of non-infected control cells was determined and set as a threshold to determine infected cells. Each nucleus with an average signal intensity for M2 ≤ 1,150 a.u. or PA ≤ 1,450 a.u. was counted as infected.

#### Βeta-lactamase (Blam) membrane fusion assay

To analyze the impact of IFNβ-1b treatment or IFITM3 overexpression in A549 cells on viral membrane fusion in late endosomes, we utilize a β-lactamase (Blam) based membrane fusion assay using influenza A virus-like particles (VLPs) expressing M1 with covalently bound Blam. Cells were laden with a fluorescence resonance energy transfer (FRET) substrate CCF4-AM (ThermoFisher Scientific, Invitrogen™), which can be proteolytically cleaved by Blam, resulting in an emission shift from 530 nm to 460 nm^84^, enabling the detection of cytoplasmic entry of M1-BlaM VLPs.

##### M1-Blam VLP production

2.7 × 106 HEK293T cells were seeded in 10 ml complete growth medium (DMEM-GlutaMAX™-I, 10% FBS, 1% P/S) in a 10 cm cell culture dish. Cells were grown overnight at 37 °C and 5% CO2. A transfection mix was prepared as follows: 1.48 µg pCAGGS-A/Hong Kong/1968-HA plasmid, 1.40 µg pCAGGS-A/Singapore/1957-NA, 7.12 µg pCAGGS-M1-Blam plasmid were added to a total volume of 1 ml Opti-MEM medium. 30 µl transfection reagent PEI (1 µg/µl, Polysciences) was added and incubated for 30 min at RT and added to the 10 cm dish with HEK293T cells. 6 h post transfection, cells were washed with PBS, and 10 ml OptiMEM medium supplemented with 1% P/S was added. After 48 h incubation at 37 °C and 5% CO2, VLP-containing supernatant was harvested and centrifuged at 1,000 × g for 10 min to remove cell debris. 2.5 µl TPCK-Trypsin (10 µg/µl, Sigma-Aldrich) was added to the supernatant and incubated for 30 min at 37 °C to proteolytically cleave HA. TPCK-Trypsin was subsequently deactivated by adding 7 µl trypsin inhibitor (5 µg/µl in PBS, Sigma-Aldrich) and incubated for 10 min at 37 °C. VLP-containing supernatant was aliquoted, shock frozen in LN2, and stored at −80 °C until further use.

##### Blam membrane fusion assay

For better cell attachment, a black-walled clear-bottom 96- well plate (Corning) was coated with fibronectin: 25 µl fibronectin solution (57.2 µg/ml fibronectin in PBS) was added to each well and incubated for 6 h at RT. The remaining solution was removed, and wells were washed 1 time with PBS. In each well, 1.2 × 104 A549 or A549- IFITM3 cells were seeded in 100 µl complete growth medium (DMEM-F12, 10% FBS, 1% P/S). The next day, A549 cells were treated with different concentrations of IFNβ-1b (0, 100, 500, 1,000, 2,000 units/ml). After 48 h, cells were infected with 190 µl M1-Blam IAV-VLPs. Infection efficiency was enhanced by spinoculation with 250 × g for 1 h at RT. Cells were washed with PBS, and 90 µl Opti-MEM supplemented with 1% P/S and 20 mM HEPES was added to each well. Cells were incubated for 3 h at 37 °C and 5% CO2. 20 µl of 6 µM CCF4- AM (ThermoFisher Scientific) in Blam loading solutions (ThermoFisher Scientific) was added to each well, and cells were incubated overnight at 8 °C on a cooling block.

##### Quantification with plate reader

Fluorescent signal was measured using the Infinite 200 plate reader (Tecan) with an excitation wavelength of 410 nm (9 nm bandwidth) and emission wavelengths of 450 nm or 520 nm (20 nm bandwidth) using a manual gain of 160 and 3 × 3 reads per well with an integration time of 20 µs and 50 reads per position. The background emission signal of the surrounding medium only was subtracted from each measurement, and the ratio between the 450 nm and 520 nm emission signal was calculated. 450 nm / 520 nm ratios of replicates for each sample were averaged. Averaged ratios were normalized to the ratio of A549 cells and reported as relative membrane fusion inhibition between 0 and 1.

##### Fluorescent microscopy

After plate reader acquisition, representative images of each well were acquired using the Nikon Eclipse Ts2 fluorescent microscope equipped with a DS-Fi3 camera, 20× / 0.4 NA lens, and a custom filter cube optimized for Blam assay image acquisition.

#### Viral entry assay by NH4Cl add-in time course

To evaluate the time for influenza A virus entry through the endosomal pathway, we performed a viral infection assay with NH4Cl treatment at different time points post-infection. As NH4Cl neutralizes the endosomal system, and thus inhibits viral membrane fusion, the half-time of viral entry can be determined as described below.

##### Viral infection with NH4Cl add-in time course

5 × 10^4^ A549 or A549-IFITM3 cells were seeded on 12 mm microscopy coverslips NO. 1 (Marienfeld) in a 24-well plate in complete growth medium (DMEM-F12, 10% FBS, 1% P/S). The next day, the cells were infected with a fluorescent reporter virus A/WSN/1933-PA-mScarlet using an MOI of 3 in serum free DMEM- F12 medium for 1 h on ice, to ensure a synchronized infection. Cells were washed 3 times with cold PBS, and cells were incubated at 37°C, 5% CO2. Cells were treated with 50 mM NH4Cl at 0 mpi, 15 mpi, 30 mpi, 60 mpi, and 120 mpi. Cells were fixed at 12 – 14 hpi with 4% PFA in PBS for 30 min at RT. After fixation, cells were washed 3 times with PBS.

##### Immunolabeling

Cells were permeabilized by incubation with 0.2% Triton X-100 in PBS for 5 min, followed by 3 washing steps with PBS for 5 min each. Cells were incubated for 1 h with a blocking buffer (3% BSA in PBS supplemented with 0.1% Tween-20 (PBS-T)). After 3 short washing steps with PBS-T, cells were incubated for 1 h with primary antibody against M2 (Proteintech) using an antibody dilution of 1:50 in a dilution buffer (1% BSA in PBS-T) at RT. After 3 washing steps with PBS-T for 5 min each, cells were incubated with DAPI using a 1:1,000 dilution (Sigma-Aldrich) and the secondary antibody ‘goat anti-Mouse Alexa Fluor™ 488’ (ThermoFisher Scientific, Invitrogen) in a dilution of 1:1,000 in the dilution buffer for 1 h at RT on a rocker. Cells were washed 3 times with PBS for 5 min each.

##### Automated high-throughput fluorescent imaging

Widefield microscopy data was acquired using the Celldiscoverer 7 automated microscope (Zeiss) equipped with a 5×/0.35 NA PL APO objective and Axiocam 712 camera.

##### Quantification

Stitched composite images were binned 2 times in ImageJ/FIJI, and nuclei were segmented using the StarDist plugin in ImageJ/FIJI83. The average signal intensity in the segmented area of each individual nucleus was calculated for M2 and PA-mScarlet. Maximum average signal intensity for M2 and PA-mScarlet of non-infected control cells was determined and set as a threshold to determine infected cells. Each nucleus with an average signal intensity for M2 ≤ 750 a.u. or PA-mScarlet ≤ 600 a.u. was counted as infected.

##### Determination of viral entry half-time

The calculated infection rates of three independent experiments were plotted against the NH4Cl add-in times and a four parameter logistic (4PL) curve was fitted. The inflection point (IC50) was determined and reported as viral penetration at half-time *t^1/2^*.

#### Transmission electron tomography (ET)

To analyze the late endosomal morphology and quantify the number of ILVs, we performed ET on high-pressure frozen and freeze-substituted (HPF/FS) cells embedded in HM20 Lowicryl. To assess the localization of IFITM3, on-section immunolabelling with anti-IFITM3 and 10 nm Protein A Gold was performed.

##### High-pressure freezing (HPF)

2.8 × 105 A549 or A549-IFITM3 cells were seeded on carbon-coated 3 mm sapphire discs (Leica) in a 6-well plate in complete growth medium (DMEM-F12, 10% FBS, 1% P/S). A549 cells were treated with 2 × 103 units/ml IFNβ-1b (Immuno Tools). After 24 h IFNβ-1b treatment, cells were fixed by HPF: Carrier type A and B (Leica) were coated with 1-hexadecene. Excess 1-hexadecene was removed by blotting with a filter paper (Whatman, No. 1). The 100 µm deep inner groove of carrier A was filled with a complete growth medium. The sapphire disc with cells was blotted on filter paper (Whatman, No. 1) and placed with the cells facing down on carrier A. Carrier B was placed with the flat site on top to close the sandwich. The sample was subsequently high-pressure frozen using an EM ICE HFP system (Leica).

##### Automated freeze substitution (AFS) and resin embedding

Lowicryl solution was prepared by adding 34.04 g monomer E, 5.96 g crosslinker D, and 200 mg initiator C (Lowicryl HM20 resin kit, Polysciences Inc.). Freeze-substitution solution was prepared by dissolving 0.1 % uranyl acetate (UA) in acetone. All solutions and high-pressure frozen sapphire discs were loaded to the automated freeze substitution system AFS2 (Leica). A freeze substitution program was performed according to the following table:

**Automated freeze substitution program:**

| Step | T start | T end | Slope | Time | Reagent | Transfer | Agitation | UV |
| --- | --- | --- | --- | --- | --- | --- | --- | --- |
| 1 | -90°C | -90°C |  | 48 h | 0.1% UA | stay | off | off |
| 2 | -90°C | -45°C | 5 °C/h | 9 h | 0.1% UA | stay | off | off |
| 3 | -45°C | -45°C |  | 5 h | 0.1% UA | stay | off | off |
| 4 | -45°C | -45°C |  | 10 min | Acetone | exch/fill | off | off |
| 5 | -45°C | -45°C |  | 10 min | Acetone | exch/fill | off | off |
| 6 | -45°C | -45°C |  | 10 min | Acetone | exch/fill | off | off |
| 7 | -45°C | -45°C |  | 4 h | 10% Lowicryl | mix | on | off |
| 8 | -45°C | -45°C |  | 4 h | 25% Lowicryl | mix | on | off |
| 9 | -45°C | -35°C | 2.5 °C/h | 4 h | 50% Lowicryl | mix | on | off |
| 10 | -35°C | -35°C | 2.5 °C/h | 4 h | 75% Lowicryl | mix | on | off |
| 11 | -35°C | -35°C |  | 10 h | 100% Lowicryl | exch/fill | off | off |
| 12 | -35°C | -35°C |  | 10 h | 100% Lowicryl | exch/fill | off | off |
| 13 | -35°C | -35°C |  | 10 h | 100% Lowicryl | exch/fill | off | off |
| 14 | -35°C | -35°C |  | 48 h | 100% Lowicryl | stay | off | on |
| 15 | -35°C | 20°C | 5 | 9 h | 100% Lowicryl | stay | off | on |
| 16 | 20°C | 20°C |  | 48 h | 100% Lowicryl | stay | off | on |
| 17 | 20°C | 20°C |  | 72 h | 100% Lowicryl | stay | off | off |

##### Ultramicrotomy

Lowicryl embedded samples were trimmed and cut with a diamond knife (DiATOME) using a UC7 ultramicrotome (Leica) to obtain sections with 250 nm nominal thickness. Sections were placed on 2 × 1 mm copper slot grids (Gilder) with 1% formvar film, sputter-coated with 2 nm carbon using ACE600 sputter coater (Leica).

##### Immunolabeling

Sections were blocked for 30 min in blocking solution (PBS supplemented with 0.8% BSA and 0.1% FSG) and incubated for 1 h with the primary antibody against IFITM3 (Proteintech), using an antibody dilution of 1:5 in blocking buffer. Sections were washed 5 times for 5 min each with PBS. Samples were incubated for 1 h with 10 nm Protein-A Gold (PAG) (Aurion) diluted 1:50 in a blocking buffer. Sections were washed 5 times for 5 min each with PBS and subsequently 5 times for 2 min in H2O.

##### Tilt series acquisition

Tilt series were acquired on a 300 keV transmission electron microscope (TEM) Tecnai F30 (FEI), equipped with a Gatan OneView 4K camera, from +60° to −60° tilt with 1° steps using a pixel spacing of 1.03 nm using the SerialEM software85.

##### Segmentation and quantification

Tomograms were reconstructed in a batch with patch tracking and R-weighted back projection using the IMOD software package86. To remove bias in data analysis, a de-identified dataset was created. All late endosomes in the reconstructed tomograms were segmented in three dimensions using the IMOD software package86. The endosome was manually segmented every 10 computational slices using the sculpt drawing tool. Segmented sections were interpolated to generate a volume. Volume segmentation was meshed and the total volume inside the mesh was calculated. The number of ILVs was manually counted for each late endosome. The ILV density was calculated for each late endosome by dividing the number of ILVs by the volume of the late endosome.

##### Cholesterol-analog cross-linking and immunoprecipitation

A549-IFITM3 cells were seeded in 10 cm cell culture dishes at a density of 4 × 106 cells per dish in a complete growth medium (DMEM-F12, 10% FBS, 1% P/S). After 24 h, cells were labeled with a photo-reactive and clickable cholesterol analog (pacChol) (Sigma-Aldrich) with a final concentration of 6 µM for 30 min at 37 °C and 5% CO2. As a control, 0.096 v/v% DMSO was used. After 2 washing steps with phosphate-buffered saline (PBS) supplemented with 1mM MgCl2 and 0.1 mM CaCl2, UV crosslinking was performed using the BIO-SUN device (Vilber Lourmant) with 5 J/cm^2^ for 20 min. Cells were lysed using 500 µl lysis buffer (PBS with 1% Triton X-100, 0.1% SDS, 2× protease inhibitor cocktail, 0.125 units/µl benzonase) per 10 cm dish for 1 h at 4 °C on a rotary wheel. The lysate was centrifuged for 15 min at 4 °C with 16,000 × g. 39 µl Biotin click reaction working solution (1,000 µM CuSO4, 100 µM Biotin-azide, 1,000 µM TCEP, 100 µM TBTA) was added to 500 µl of lysate supernatant and incubated for 3 h at 37 °C on a shaker with 1,000 rpm. Click-reaction was stopped by adding 4 ml of ice-cold methanol. The sample was centrifuged for 20 min at 4 °C with 3,000 × g. The supernatant was discarded, and the pellet was washed 2 times with ice-cold methanol and subsequently air-dried. Pellet was resuspended in 15 µl of 4% SDS in PBS and dissolved by sonication. The sample was diluted 1:40 in PBS with 1% Triton X-100 for further analysis. 20 µl high capacity NeutrAvidin (Sigma-Aldrich) agarose beads were added to 600 µl sample and incubated overnight at 4 °C. The sample was centrifuged for 3 min at 4 °C with 3,000 × g. Beads were washed 3 times with PBS supplemented with 1% Triton X-100. The sample was eluted with 20 µl Laemmli buffer and further analyzed by immunoblotting using anti- IFITM3 (1:2,000) and anti-Caveolin1 (1:1,000) antibodies on a pre-cast 10 – 20% Tricine gel (ThermoFisher Scientific). The ‘Precision Plus’ (Bio-Rad) prestained protein size marker was used.

##### Molecular dynamics simulations

The initial IFITM3 structure (residues 59 – 133) and its orientation in the membrane was kindly provided by Emma H. Garst21. IFITM3 was inserted into model membranes using the CHARMM-GUI Membrane Builder web interface87,88. First, the palmitoyl group was added at Cys^72^ during the membrane builder system building process. Next, several potassium or chloride atoms were added to neutralize the systems. Finally, KCl salt at a concentration of 150 mM was added to mimic the intracellular environment. Protein, lipids, and salt ions were described using the CHARMM36m force field^89–91^. For water, we used the TIP3 mode^92^.

Equilibration of the systems was conducted with CHARMM-GUI standard protocol. Briefly, all the systems were subjected to energy minimization using the steepest descent algorithm. After minimization, we ran six steps of equilibration runs where we gradually reduced the force constant applied to restrain the positions of the protein and the lipids. For the production run, we employed the Parrinello-Rahman barostat^93^ with a semi-isotropic pressure coupling scheme and a time constant set to 5.0 ps to maintain the pressure constant. The pressure was set to 1.0 bar and the isothermal compressibility to 4.5 × 10–5 bar–1. The temperature was maintained at 310 K using the Nose-Hoover thermostat94,95 with a time constant of 1.0 ps. Electrostatic interactions were handled using the PME method96,97. The cut-off length of 1.2 nm was used for electrostatic (real space component) and van der Waals interactions. Hydrogen bonds were constrained using the LINCS algorithm98. Finally, periodic boundary conditions were applied in all directions. The simulations were carried out using an integration time step of 2 fs with coordinates saved every 100 ps. Due to the presence of 8 different lipid types, the late endosomal-like membrane systems were simulated for 6 microseconds, whereas the binary and tertiary lipid mixtures were simulated for 1 microsecond. All simulations have been carried out usingthe GROMACS-2021 software99. The images were rendered using VMD100.

##### Late endosomal membrane composition

The evaluation of the lipid composition in the late endosome is highly inaccurate and should be considered a rough estimation since the lipid composition changes as the endosome matures. Moreover, the ILVs and the limiting membrane have different lipid compositions101. The cholesterol/phospholipids ratio decreases from 1/1 at the early endosome to 1/2 in the late30. The major phospholipids head groups in the late endosome are^29^ phosphatidylcholine (PC) 48.2%, phosphatidylethanolamine (PE) 19.8%, lysobisphosphatidic acid (LBPA) 15.7%, Sphingomyelin (SPH) 9.2%, phosphatidylinositol (PI) 4.1% and phosphatidylserine (PS): 2.4%. The level of acyl chain saturation is unknown in the late endosome, so we estimate it based on the known values in the plasma membrane102. The majority of LBPA tails, which is not present in the plasma membrane, are 18:1 (∼ 92%)29. Finally, the percentage of lysolipids (excluding LBPA) is about 4 – 5% in all the cell organelles examined103. The used late endosomal-like lipid composition used for MD simulation is summarized in the following table:

**Complex lipid composition mimicking the late endosomal membrane**

**Complex lipid composition mimicking the late endosomal membrane:**
| Abbreviation | Full name | Mole percent | Intrinsic curvature | References for intrinsic curvature |
| --- | --- | --- | --- | --- |
| POPC | 1-palmitoyl-2-oleoyl-sn-glycero-3-phosphocholine | 35.1 | -0.02 | Kollmitzer et al.<sup>104</sup> |
| POPE | 1-palmitoyl-2-oleoyl-sn-glycero-3-phosphoethanolamine | 7.3 | -0.32 | Kollmitzer et al.<sup>104</sup> |
| DOPE | 1,2-dioleoyl-sn-glycero-3-phosphoethanolamine | 7.3 | -0.35 | Fuller and Rand<sup>105</sup> |
| PSM | Palmitoylsphingomyelin | 5.3 | -0.13 | Kollmitzer et al.<sup>104</sup> |
| LBPA | Lysobisphosphatidic acid | 9.9 | Negative, exact value unknown | Kobayashi et al.<sup>29</sup> |
| CHOL | Cholesterol | 29.8 | -0.5 | Kollmitzer et al.<sup>104</sup> |
| LPC16 | Lysophosphatidylcholine | 4.0 | 0.15 | Fuller and Rand<sup>105</sup> |
| LPE16 | Lysophosphatidylethanolamine | 1.3 | 0.03 | Fuller and Rand<sup>105</sup> |

##### Enrichment/Depletion analysis

For enrichment/depletion analysis, we calculate the number of lipids (for each lipid type) within a radial cut-off distance of 0.4. In addition, the molar fractions of lipids surrounding IFITM3 have been calculated with respect to the lipid bulk concentrations (lipids above the radial cut-off distances). The analysis was carried out employing the ‘gmx select’ and in-house python scripts. The first 20 ns have been considered as equilibration time and then excluded from the analysis. The standard deviation is reported in the error bar with n = 3.

##### Normalized contact occupancy

Over time, the number of contacts has been calculated using the ‘gmx mindist’ GROMACS analysis tool. A contact is defined if the distance between any atoms of IFITM3 and POPC (or cholesterol) is less than 0.4 nm. Data are normalized based on the total number of lipids of each lipid type. The first 200 ns have been considered as equilibration time and then excluded from the analysis. The error bar is shown as the standard deviation.

#### Relative affinity and repulsion of lipid to IFITM3

We calculate the relative affinity and repulsion of different lipid species to IFITM3 by considering the two competing forces governing the lipid distribution – the direct interaction between lipid to IFITM3 and the demixing entropy that resists it. To do so, we use the following assumptions: Each monolayer is composed of *M* number of different lipid species, and each lipid species has *n*_j_ lipids in each monolayer. No lipid flip-flop was observed during the MD simulations; therefore, we prohibit it in the following derivation. We define two regions in each monolayer – the immediate vicinity of the IFITM3 (Figure 2B, green) and the bulk (Figure 2B, orange), given by the superscripts *b* (bound) and *f* (free), respectively. Lipids can freely diffuse between the regions, but since the lipids are closely packed and have a high stretching- compression modulus, we assume their density and number in each region are fixed, *N_b_*^3^ and *N_b_*^4^, respectively. Therefore, the free energy can only be relaxed by lipid exchange between the bound and free regions, Δ*n_j_*, which must satisfy the following requirement:

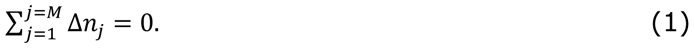

The total free energy that is associated with the lipid’s interaction energy with IFITM3 and mixing entropy is given by

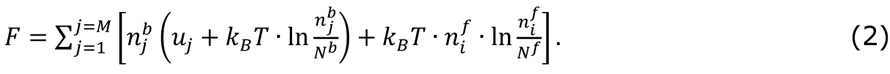

To find the equilibrium distribution, we minimize Equation 2 with respect to Δ*n_j_* while maintaining the constraint of a fixed number of lipids and Equation 1, which can be done numerically. However, MD simulations found that most lipid exchange is due to cholesterol repulsion from IFITM3. Therefore, we consider cholesterol exchange with all of the other lipids combined. With that, Equation 2 is minimized by allowing only the exchange of Δ*n* cholesterols with any other lipid. The relative repulsion of cholesterol is given by

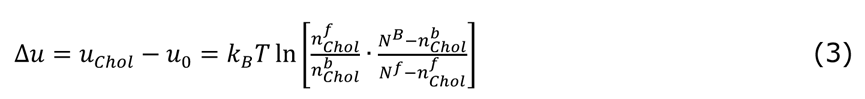

with *n^b,f^_chol_* and *N^b,f^* the number of cholesterol and the total number of lipids in the free or bound areas (donated by the super-script *b* or *f*), respectively. *u_o_* is the averaged interaction energy of IFITM3 with all other lipids except cholesterol and *u_chol_* is the interaction energy with cholesterol. This procedure is performed separately on the luminal and cytosolic facing and monolayers to find the relative repulsion of each, Δ*u_l_* and Δ*u_c_*, respectively. We also performed this procedure on the two monolayers combined to find the relative repulsion of cholesterol to IFITM3 in the membrane.

#### Modeling of membrane fusion and lipid sorting

##### Membrane elasticity theory

We use a continuum approach based on lipid tilt and splay theory to model the lipid deformations in the hemifusion diaphragm and its vicinity. Previous works present the theory and related computations in detail32–34,106. In short, the hemifusion diaphragm is modeled as a membrane surrounded by the virus and endosome membranes, and the diaphragm rim represents the merger of the three (Figure 6A). Each membrane comprises two lipid monolayers that contact along a joint mid-plane. The deformation of the lipids in each monolayer is described by their tilt, splay, and saddle-splay39,40. All are defined with respect to the monolayer dividing plane. Here we take the distance from the membrane mid-plane to the monolayer dividing plane to be δ = 1.5 nm^107^ and explicitly assume that they are parallel.

Lipid tilt deformation is generated due to the shearing of the hydrocarbon tails in the vicinity of the diaphragm rim. It is defined as 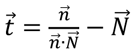 with *n* the lipid director and *N* the normal to monolayer dividing plane^39^, as presented in Figure 3C. Splay deformations originate from the curvature deformations and the lateral change of the lipid tilt39,108. Mathematically, the lipid splay tensor is defined as the covariant derivative of the lipid director,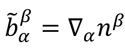, where the sub- and superscripts denote, respectively, the co- and contravariant components in the local coordinate basis of the monolayer dividing plane and39. Lipid splay is the trace of the splay tensor, defined as 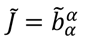, where the summation is performed over repeating indices. In the absence of tilt, lipid splay has the meaning of the sum principal curvatures, *c_1_* and *c_2_*, referred to as total curvature, *J = c_1_ = c_2_*. Similarly, lipid saddle-splay is a generalization of Gaussian curvature, *K* = *c_1_*.*c_2_*, in the presence of both bending and tilt deformations. In co-contra variant form, it is the determinant of the splay tensor^39^, 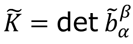. The elastic energy density is given by

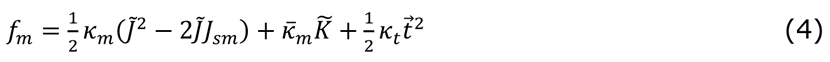

with *K*_E_, *K*_F_ and *K*_E_ the monolayer’s bending, tilt, and saddle-splay rigidities, respectively. Their values are estimated from the literature to be 14.4 kBT^109^, −7.2 kBT^110, 111^, and 40 mN/m112,113, respectively. The spontaneous monolayer curvature, *J_SM_*, is determined by the monolayer constituting lipids and proteins intrinsic curvatures, ζ_2_, and their relative mol fractions, Φ_2_. With *j* being index representing the different components. The overall spontaneous monolayer curvature is the sum of its components^104,114,115^

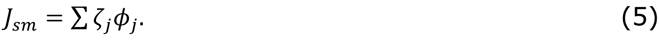

However, since IFITM3 is strongly repelling cholesterol, we consider the membrane to be composed of just two lipid subgroups – cholesterol with intrinsic curvature of ζ_>:;+_ = −0.5 nm^=/^ and all of the other lipid species grouped have the mean intrinsic curvature of ζ_<_ = −0.1 nm^=/^, we term the second group ‘background’ lipids.

The overall elastic energy is found by integrating the energy density in Equation 4 over the area of the monolayers

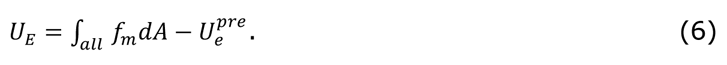

The elastic energy is with respect to the pre-fusion configuration of two flat non-interacting membranes, *U^pre^_e_*. The equilibrium shape of the fusion site is found by minimizing Equation 6, as explained in the following.

*IFITM3-induced lipid sorting*: In addition to lipids, the endosomal membrane contains uniformly distributed IFITM3 proteins that repel cholesterol and attract other lipids. The virus and diaphragm membranes do not contain IFITM3 because the diaphragm rim acts as a semi-permeable barrier that allows only lipids to pass. The formation of the IFITM3-free region allows the system to relax some of the relative repulsion energy between cholesterol and IFITM3 at the cost of entropic demixing energy.

To calculate the magnitude of exchange, we consider two membrane domains with a total of *N_b_* lipids. One domain represents all the lipids that are closely bound to IFITM3 with *N_b_*^3^ = *N_b_*_IJIK6L_ · *n*_"&F.#_ lipids, *N_b_*_IJIK6L_ is the total number of IFITM3 proteins in the endosome and *n*_"&F.#_ the number of lipids interacting with each, as found in MD simulations (*n*_IJIK6L_ = 52 was used in all simulations as the value found in the ‘late endosome composition). The second domain contains the remaining lipids *N_b_*^4^ = *N_b_* − *N_b_*^3^ = *N_b_* − *N_b_*_IJIK6L_ · *n*_"&F.#_. As explained above, we consider two subgroups of lipids: *N_b_*_>:;+_ cholesterols lipids and *N_b_*_<_ = *N_b_* − *N_b_*_>:;+_ background lipids. The relative repulsion of the two to IFITM3 is Δ*u* = *u*_9:;+_ − *u*_<_. The number of cholesterol lipids is *n*^4^ and the number of background lipids is *n*^4^ = *N_b_*^4^ − *n*^4^ = *N_b_* − *N_b_*_IJIK6L_ · *n*_"&F.#_ − *n*^4^ in the free domain. The lipid numbers in the bound domain are *n*^3^ = *N_b_*_>:;+_ − *n*^4^ and *n*^3^ = *N_b_*^3^ − >:;+ >:;+ < *n*^3^ = *N_b_*_IJIK6L_ · *n*_"&F.#_ − *N_b_*_>:;+_ + *n*^4^ . To find the equilibrium distribution, one can use a similar >:;+ >:;+ procedure leading to Equation 3 to have

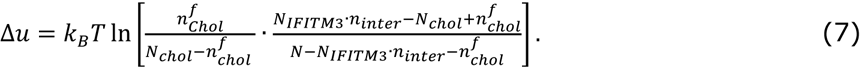

We define the mole fractions: Φ_IJIK6L_ = *N_b_*_IJIK6L_⁄*N_b_* , Φ_>:;+_ = *N_b_*_>:;+_⁄*N_b_*, and Φ^4^ = *n*^4^ and rewrite Equation 7 as

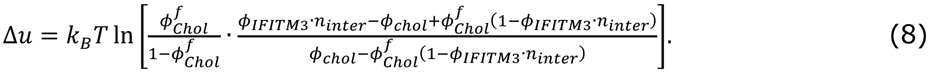

The total cholesterol mole fraction is estimated based on the known literature to be Φ_>:;+_ = 0.3 in the late endosome^116^, Δ*u* is found based on the MD simulation as described above and Φ_IJIK6L_ is the free parameter of the model. We find Φ^4^ by rearranging Equation 8 and solving the quadratic equation

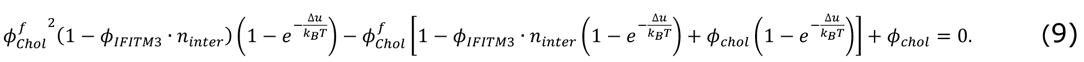

The magnitude of lipid exchange is defined as the change of cholesterol level from the baseline levels with no IFITM3 in the endosome 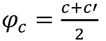.

Next, we derive the spontaneous curvature in the endosome, virus, and diaphragm monolayers. Since the relative affinities of lipids to IFITM3 are similar on the luminal and cytosolic sides, we consider the magnitude of the lipid exchange to be the same in both leaflets. In addition, because the typical time in hemifusion is in the order of seconds while the flip-flop time of cholesterol is measured in milliseconds, we also assume the system to be in equilibrium with respect to lipid flip-flop.

The virus and hemifusion diaphragm have the lipid composition of the ‘free’ domain, and their monolayer spontaneous-curvature is (Equation 5)

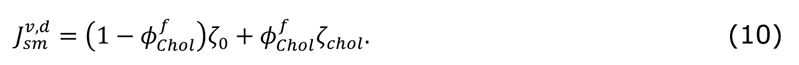

The endosomal membrane contains patches of both free and bound domains since IFITM3 proteins are dispersed in it. However, since the IFITM3 freely diffuse and are uniformly distributed, the lipids are also uniformly distributed on time-average. So, although IFITM3 triggers a local change in lipid composition in its immediate vicinity (< 1 nm), we treat the endosomal membrane as a smooth continuum and average other these spatial fluctuations. Moreover, we also consider the endosomal membrane much larger than the virus, so the enrichment of cholesterol in the virus has a negligible effect on the cholesterol mole fraction in the endosome. Lastly, we account for possible intrinsic curvature of the amphipathic helix domain of IFITM3 that was recently reported 23 in the ‘with sorting and curvature’ and ‘no sorting; with curvature’ scenarios. Specifically, we attribute the intrinsic curvature of ζ^>^ = −0.62 nm^=/^ to the cytosolic side because there the amphipathic helices are located23. Given all of that, the endosome cytosolic facing monolayer spontaneous-curvature is (Equation 5)

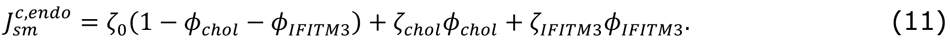

Here we also assumed for simplicity that the amphipathic helix domain has the same projected size as a lipid. The luminal-facing side of IFITM3 contains a transmembrane domain, which does induce any curvature to our knowledge. Therefore, the endosome luminal-facing monolayer spontaneous curvature is (Equation 5)

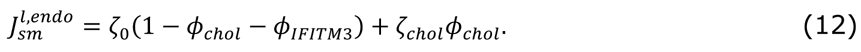

In the ‘with sorting; no curvature’ scenario, we set ζ_IJIK6L_ = 0. We also tested the possibility that the amphipathic helices of IFITM3 penetrate the hemifusion diaphragm. In such a case, there is no sorting. The monolayer spontaneous-curvature of the diaphragm cytosolic facing monolayer is equivalent to the endosomal one.

#### Pore Expansion energy barrier and first passage time

##### The magnitude of fusion-pore formation energy barrier

A membrane pore must open and expand in the hemifusion diaphragm to complete the fusion reaction117,118. The driving force for fusion pore expansion is the removal of lipids from the highly stressed region in the diaphragm and their migration to the more relaxed surrounding membrane. The energy gained by this process is given by

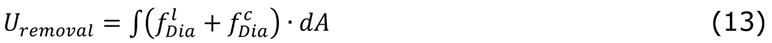

with *f_Dia,_* being the position-dependent energy density in the diaphragm monolayers (Equation 4). The superscripts *l* and *c* represent the diaphragm luminal and cytosolic facing monolayers, respectively. The integral in Equation 13 is done over the pore area of size *A*. The fusion pore expansion is resisted by the energetic cost of forming the pore rim, *U_porerim_* _#"E_ = *λ*λ, with *λ* to pore rim line tension and λ the pore rim length. The line tension depends strongly on the lipid composition. In a two-lipid component system, the effect of the relative mol fraction change, ΔΦ, is given by^119^

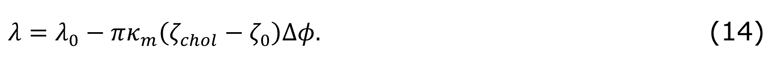

The first term in Equation 14, *λ*_<_, accounts for the contribution to line tension independent of change in lipid composition. It includes all the other lipid deformations in the pore rim. Its value is estimated as *λ*_<_ = 15 pN for a membrane containing 30% cholesterol based on theoretical considerations^117^ and experiments37. The energy needed for pore expansion is the sum of the energy cost of creating the pore rim and the energy gained by the removal of lipids (Equations 13 and 14)

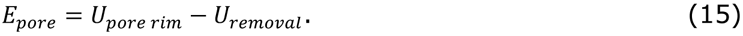

The pore-opening energy, *E^*^*_*;#._, is characterized by the critical pore radius, *p*^*^, smaller pores tend to close while larger tend to expand infinitely (Figure 3E). The energy of pore expansion at the critical size is the pore expansion energy barrier,

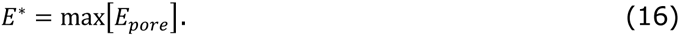

##### The dwell time in hemifusion

Thermal fluctuations determine the dynamics of the pore size; small pores flicker in size until they eventually reach the critical size beyond which they irreversibly expand120. The mean time needed for the pore size to cross the energy barrier was already described in detail in earlier works, primarily for uniformly stressed membranes^121^ but also in the specific case of pores in the hemifusion diaphragm^34^

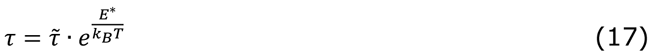

with *j* being a pre-exponent factor that depends on an unknown microscopic time constant. We evaluate the efficiency of IFITM3 in inhibiting fusion by calculating the ratio between the typical time for pore expansion with and without IFITM3, *t_ifitM3_* and *t_o_* respectively

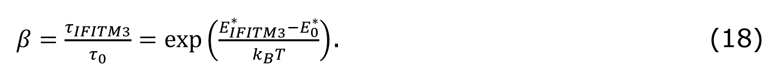

Here we assume that *t* is unchanged by slight changes in lipid composition, and the change in the diaphragm area available for fusion is negligible. We term *β* as the ‘inhibition factor’ and the difference in pore expansion energy barrier due to IFITM3 presence as Δ^∗^ = *E^*^*^∗^ −*E^*^*^∗^.

*Numerical computation:* Our computation aims to estimate the inhibition factor of IFITM3. The computation is divided into two steps. First, we calculate the equilibrium shape of the hemifusion diaphragm by minimizing Equation 6 using an iterative approach. The membranes’ mid-planes and tilt angles are gradually changed until minimal energy configuration is found.

The geometry of the fusion site is described by the junction angles (Figure 3A), *φ_v_* and *φ_c_*, the diaphragm size *R_D_*, fusion site size at the virus and endosome sides *R^endo^_r_* and *R^v^_r_*, and the distance between the virus and endosome, ℎ, is also subjected to energy minimization. The process is repeated from different initial configurations to ensure that minimal energy is genuinely found.

Next, based on the resulting equilibrium configuration and the adjusted lipid composition in the diaphragm, we calculate the energy barrier of pore expansion, Δ^∗^, (Equation 16), and the inhibition factor *β* (Equation 18). The pore is more likely to initially form close to the diaphragm rim, where the stress is maximal. However, our computations show that the critical pore size is comparable to the size of the diaphragm, making the energy barrier mostly independent of the starting position. Therefore, to facilitate the computations, we assume that the fusion pore growth starts at the center of the diaphragm and radially expands. We also assume that the diaphragm’s shape is unaffected by the growth of the pore. All computations are done using self-written MATLAB code, which is available on GitHub (https://github.com/GonenGolani/Fusion_Solver).

#### *In situ* cryo-correlative light and electron microscopy (cryo-CLEM)

*In situ* cryo-CLEM was performed on cryo-FIB milled samples based on a previously established method^42,48^, as described below:

##### Sample preparation

1.8 × 10^5^ A549 or A549-IFITM3 cells were seeded on glow-discharged 200 mesh gold grids with a holey SiO2 film with R1/4 spacing (Quantifoil) placed in a 35 mm dish and cultured in 2 ml complete growth medium (DMEM-F12, 10% FBS, 1% P/S). To facilitate grid handling, the 35 mm dish was coated with polydimethylsiloxane Sylgard 184 (Dow Corning) as described in Klein et al.^122^. The next day, cells were infected with fluorescently labeled IAV (A/WSN/1933-nDiO) using an MOI of 200. Virus stock was diluted to a final concentration of 5 × 106 PFU/ml in the infection medium (DMEM-F12, 20 mM HEPES). EM grids were blotted on filter paper (Whatman No. 1) and placed on Parafilm. 20 µl of the virus dilution was added onto the grid and incubated for 30 min at 12 °C on a cooling plate to allow viral cell attachment. EM grid was washed 5 times with infection medium and was subsequently incubated for 1 h at 37 °C, 5% CO_2_ in the complete growth medium, supplemented with 100 nM Lipi-Blue (Dojindo) to fluorescently label lipid droplets. 1 hpi, infected cells were washed 2 times with complete growth medium, grids were manually blotted on filter paper (Whatman No. 1) and a defined volume of 3 µl complete medium was added to the grid to ensure identical blotting conditions. Grid was subsequently blotted and vitrified by plunge freezing in liquid ethane using the EM GP2 automated plunge freezing device (Leica) using the following settings: 70% air humidity, 25 °C air temperature, −183 °C ethane temperature, 3.5 sec blotting from the back using a filter paper (Whatman No. 1).

##### Cryo-light microscopy (cryo-LM)

Fluorescent signal of the plunge frozen samples were acquired on the cryoCLEM wide-field microscope (Leica Microsystems)123 equipped with a 50x/0.9 NA objective. A 1.2 × 1.2 mm area of the center of each grid was mapped, acquiring a volume stack of 30 µm thickness with a Z-spacing of 300 nm using the LAS X Navigator software (Leica). Image stacks were deconvolved with AutoQuant X3 (Media Cybernetics) using a theoretical and adaptive point spread function (PSF) for 100 iterations with the following parameters: lens immersion refractive index 1; sample embedding refractive index 1.31; sample distance from coverslip of 0 nm; emission wavelength of 460 nm (for Lipi-Blue) or 525 nm (for nDiO) and appropriate settings for the used objective (NA 0.9; objective lens magnification 50x). For cryo-correlative light and electron microscopy (cryo-CLEM), lamellae were mapped again by cryo-LM after cryo-TEM and correlated with the cryo-TEM map of the lamella using the post-correlation cryo-CLEM toolbox122.

##### Cryo-focused ion beam milling (cryo-FIB)

Samples were mapped by cryo-scanning electron microscopy (cryo-SEM) on an Aquilos dual-beam cryo-focused ion beam-scanning electron microscope (cryo-FIB-SEM) (Thermo Fisher Scientific) with a cryo-stage cooled to −180 °C. The cryo-LM map of the grid was correlated to the cryo-SEM map using the MAPS Software (Thermo Fisher Scientific)124. Infected cells were selected for milling. A protective organo- metallic platinum layer was applied for 5 sec to the grid and cells were milled gradually in 5 steps with a stage angle between 15° and 18° using a Gallium ion beam125. The nominal thickness of the milled lamella at the last milling step was 150 nm. A milling pattern including micro-expansion joints was used to minimize lamella bending126.

##### Cryo-electron tomography (cryo-ET)

Cryo-ET data was acquired on a Krios cryo-TEM (Thermo Fisher Scientific) operated at 300 keV equipped with a K3 direct electron detector (Gatan) and Quanta Gatan Imaging Filter (Gatan) with an energy slit set to 20 eV. Lamellae were mapped using SerialEM^85^ at 8,700× (pixel spacing of 10.64 Å) using a defocus of −50 µm to localize multivesicular organelles which were selected for acquisition. Tilt series were acquired with SerialEM^85^ using a dose-symmetric tilting scheme^127^ with the zero angle set to 8° and a nominal tilt range of 68° to −52° with 3° increments using the following settings: Target focus of −3 µm, electron dose per record of 3 e-/Å2 and a magnification of 33,000× (pixel spacing of 2.67 Å). Automated tilt series acquisition was performed in SerialEM^85^ using the Navigator with either a realign-to-item function or scripts for Parallel Cryo Electron Tomography (PACE)128.

##### Tomogram reconstruction

Individual movie frames were split into even and odd and subsets were drift corrected using Motioncor2129. Even and odd datasets were reconstructed in parallel with identical settings in etomo as part of the IMOD software package^86^ using patch tracking for tilt series alignment. CTF estimation and 3D CTF-correction were performed in etomo. Tomogram was reconstructed using weighted back-projection with a simultaneous iterative reconstruction technique (SIRT)-like filter equivalent to 5 iterations and dose-weighting filter. Denoising was performed using the reconstructed tomograms from the even and odd datasets employing the Noise2Noise algorithm implemented in the CryoCare software package^130^ on non-binned data.

##### Volume rendering

The reconstructed and denoised tomogram volume was segmented in 3 dimensions using the Amira software (Thermo Fisher Scientific). First, a non-local means and membrane enhancement filter were applied. Membranes were automatically segmented using the Top-hat tool. The automated segmentation was manually refined by removing falsely segmented areas. In addition, missing parts of the limiting late endosomal membrane, ILVs and the viral membranes were manually segmented. The number of direct interactions between the viral particles and ILVs or the limiting late endosomal membrane were quantified.

##### Analysis of hemifusion symmetry

For each hemifusion site at the limiting late endosomal membrane or at ILVs in the endosomal lumen the length of the hemifusion diaphragm and the inner and outer angles (as defined in Figure 4A) were measured. The tomogram was resliced using the slicer tool in IMOD^86^ to position the hemifusion site on a single X-Y plane. All individual angles were measured using the ‘angle tool’ in ImageJ/FIJI80. To measure the hemifusion diaphragm length, a line profile with 10 pixel width was plotted. The average signal of the late endosomal lumen was subtracted. The difference between the zero-crossing points was determined and reported as the diaphragm length. Thus, the measurement includes the phospholipid monolayers.

#### Estimation of pore formation energy barrier based on cryo-EM data

We use the theory described above to estimate the magnitude of the fusion-pore formation energy barrier based on the cryo-EM data. The stress in the diaphragm depends only on the geometry of the fusion site and specifically on the tilt magnitude at its rim, given by

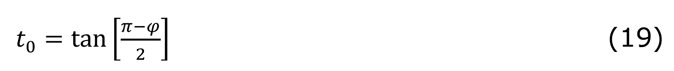

With *φ* the inner angle of the fusion site either and the virus or cytoplasmic side (Figure 5A). Since we assume that the diaphragm is axially symmetric and no significant differences were observed between opposing angles, we take the inner angle as the average 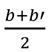 = ^>M>V^ and *φ*_!_ = 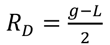 (Figure 5B). The lipid tilt propagates towards the center of the diaphragm and vanishes at its center. The tilt radial dependence can be approximated to^34^

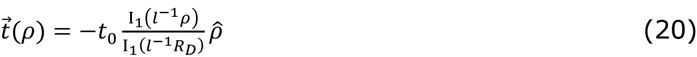

With I_\_ the modified Bessel functions of the first kind, *ρ* the radial coordinate, nm the tilt decay length. The diaphragm mid-plane radius, *R_D_*, is related to the measured diaphragm diameter (Figure 5C) by 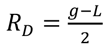. With λ the membrane thickness measured from headgroups to headgroups of opposing monolayers and the distance between head groups at the diaphragm rim. We take λ to be 6 nm based on our cryo-EM data. The change in tilt induces lipid splay, and saddle-splay is given by^34^

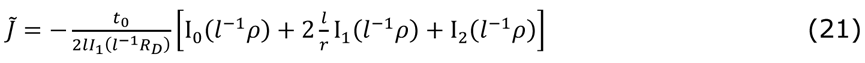

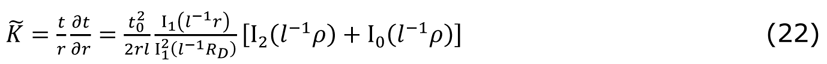

We use Equations 20–23 to calculate the tilt, splay, and saddle-splay in each monolayer separately and the membrane stress using Equation 4. Finally, each fusion site’s pore formation energy barrier is calculated using Equation 16.

#### Subtomogram averaging (STA)

To characterize the state of the influenza A fusion protein HA, which was regularly found at the hemifusion sites, we performed subtomogram averaging. In total, 30 individual putative HA densities at hemifusion sites were identified and extracted using a dipole model in Dynamo^131^ using a box size of 192 pixel (equals 51.28 nm). An initial reference model was created by averaging all subvolumes with the orientations extracted by the dipole model. Azymuth angles were randomized to evenly distribute the missing wedge of the individual subvolumes during initial model creation. The first averaging was calculated without symmetry operation (C1) using a spherical mask (Figure S14A). The resulting average of the first STA was used as a new template for the second average, using a tight mask around the central density corresponding to the HA. In the second average, a C3 symmetry operation was applied (Figure S14B) and the same numerical parameters were used. An overview of used parameters can be found in the following table:

**Numerical parameters for STA calculation in Dynamo.**

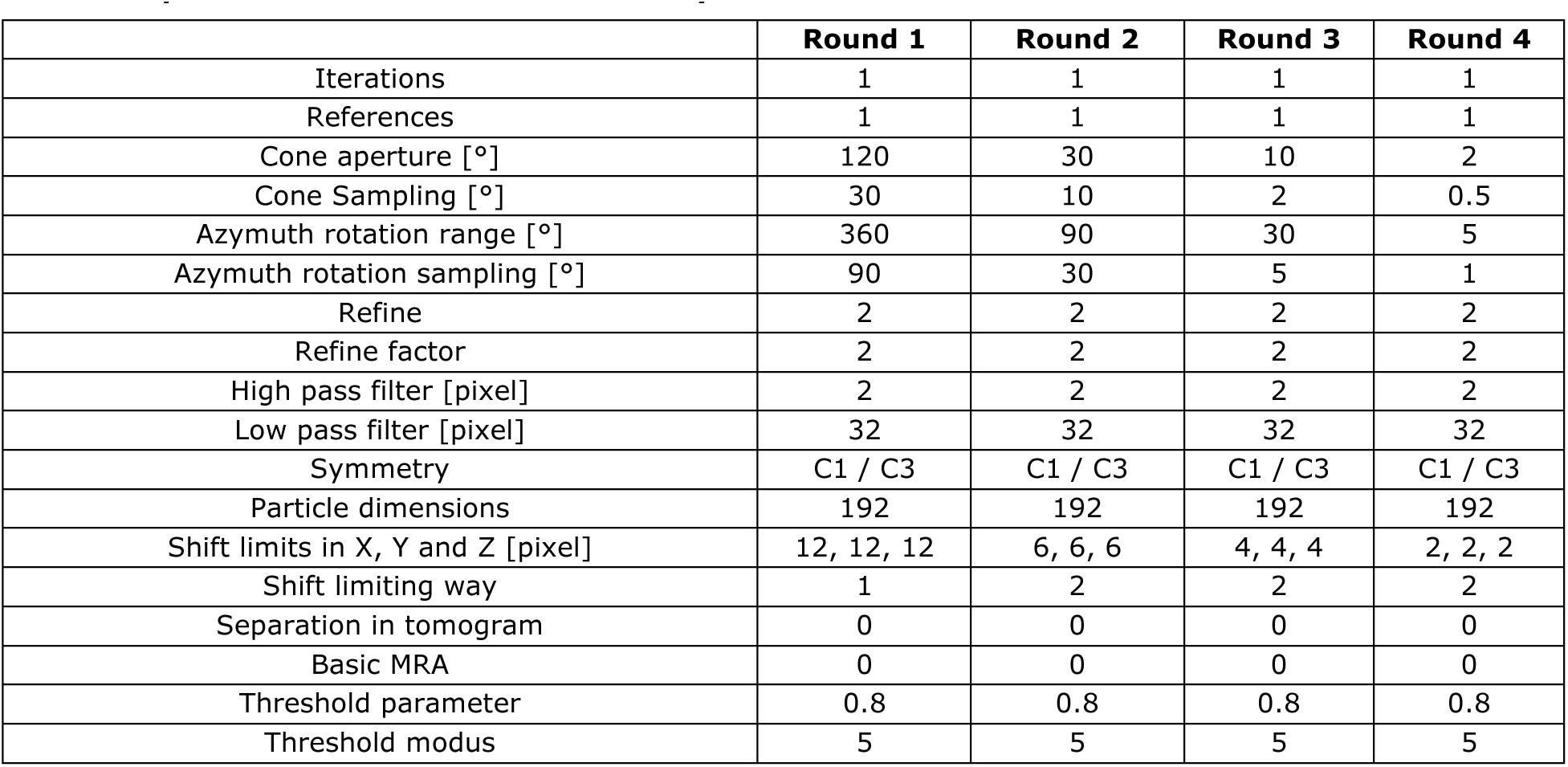

An isosurface of the final average of the STA was created using USCF Chimera132. The post-fusion structure of HA2 (PDB: 1QU1)71 was fitted to the isosurface using the ‘Fit in Map’ function in USCF Chimera. The number of atoms inside and outside of the isosurface were quantified.

### Quantification and statistical analysis

Image acquisition and quantification of ILV in late endosomes (Figure 1R–Y) was done blinded. An unpaired T-test was utilized to analyze the significance of differences in the following figure panels: Figure 1H, 1X, 1Q, 1Y, S1C, S3C, S3D, S3E, S3F. Paired T-test was utilized to analyze the significance of differences in the following figure panels: Figure 5B.

## SUPPLEMENTAL VIDEO AND EXCEL TABLE TITLES AND LEGENDS

**Supplementary Video 1 (related to** **Figure 4**). Video of the reconstuctred tomogram and 3D rendering of tomogram shown in Figure 4E and 4F.

**Supplementary Video 2 (related to Supplementary Figure 8).** Video of the reconstuctred tomogram and 3D rendering of tomogram shown in Figure S8A and S8B.

**Supplementary Video 3 (related to Supplementary Figure 9).** Video of the reconstuctred tomogram and 3D rendering of tomogram shown in Figure S9A and S9B.

**Supplementary Video 4 (related to Supplementary Figure 10).** Video of the reconstuctred tomogram and 3D rendering of tomogram shown in Figure S10A and 108B.

**Supplementary Video 5 (related to** **Figure 5**). Video of the post-fusion HA subtomogram average shown in Figure 5F–I.

**Supplementary Table 1 (related to** **Figures 1–6**). Excel sheet providing plotted values in all main and supplemental figures.

**Supplementary Table 2 (related to Figure 4 and 5).** Excel sheet providing an overview of all virus membrane contact sites.

## MATERIALS

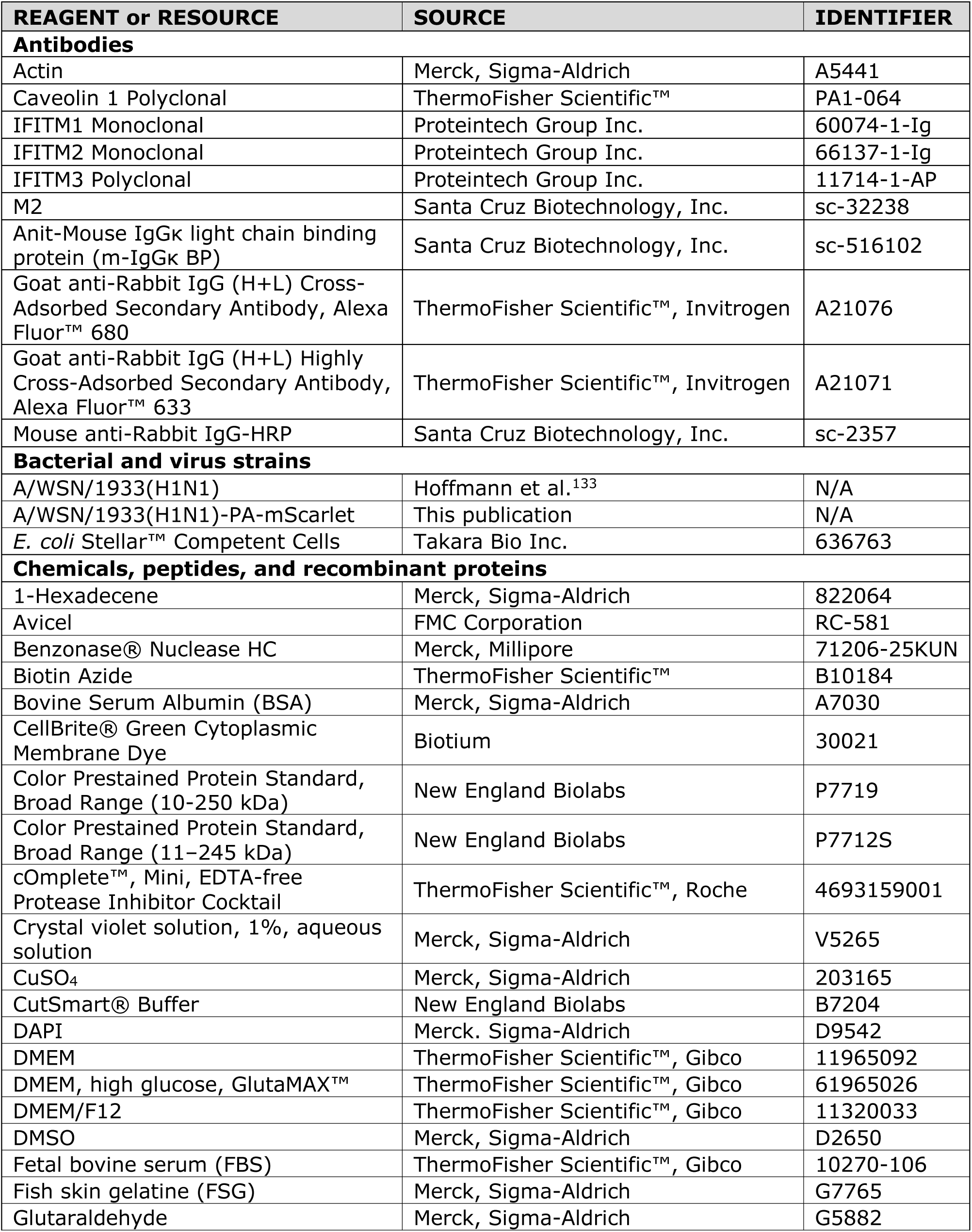
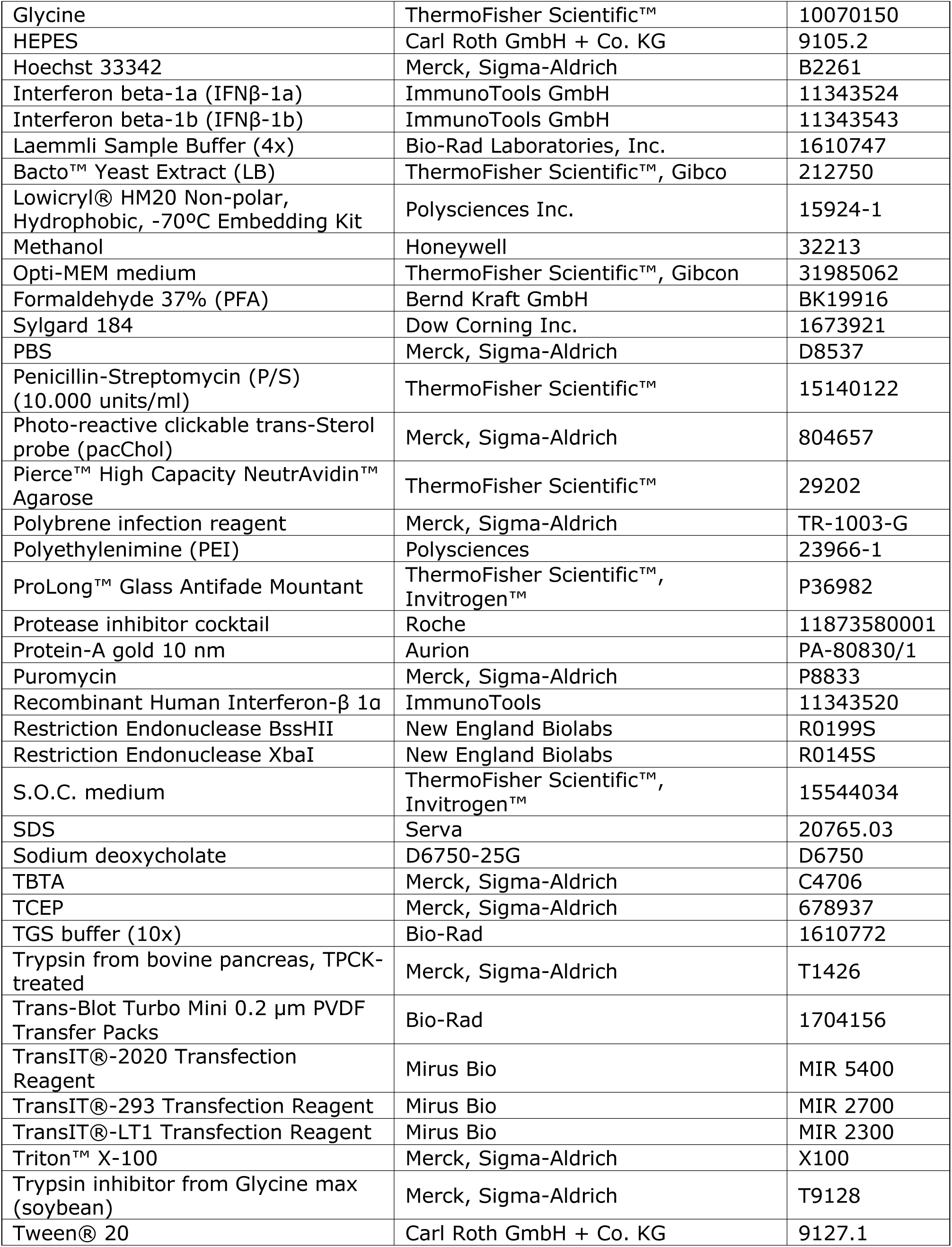
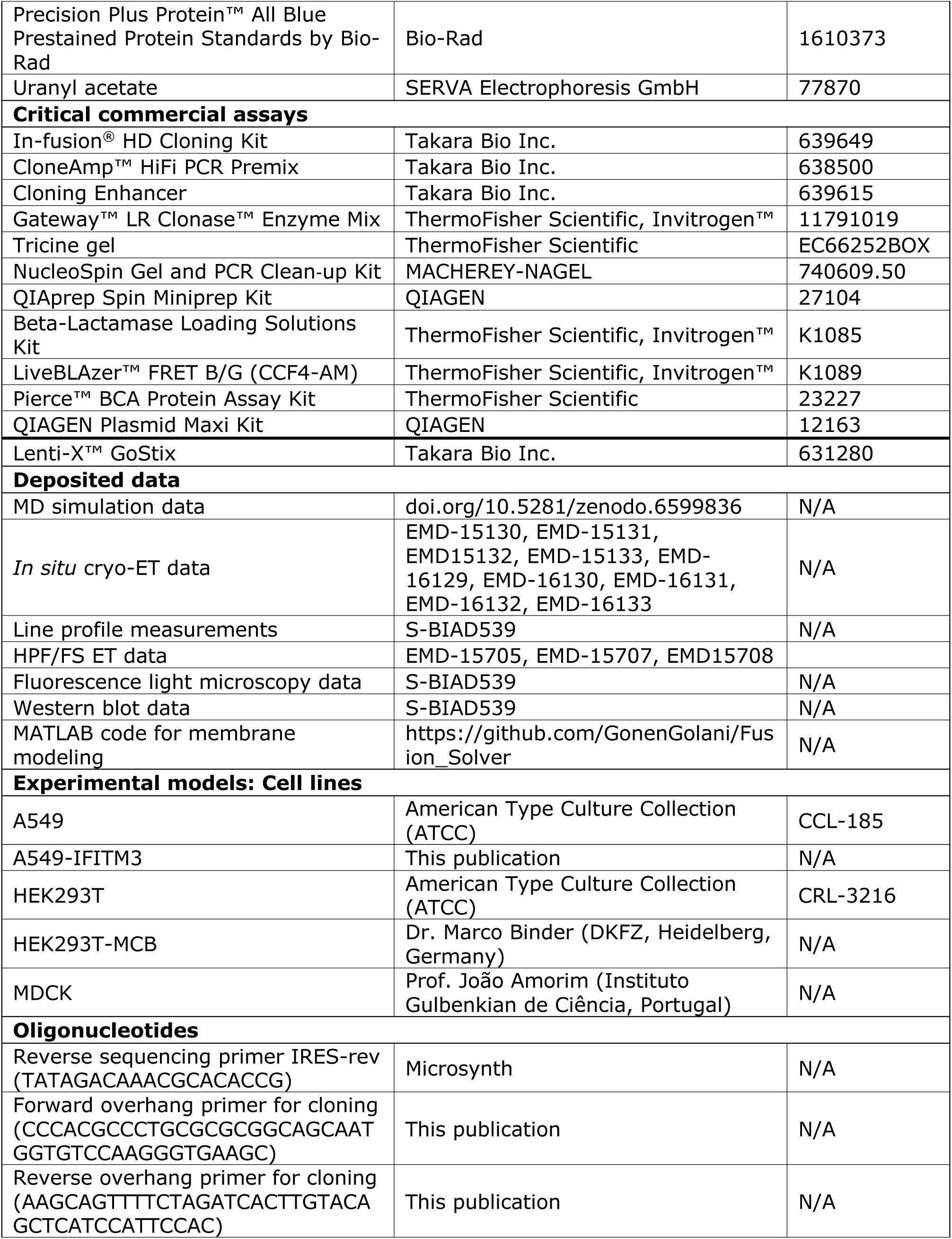
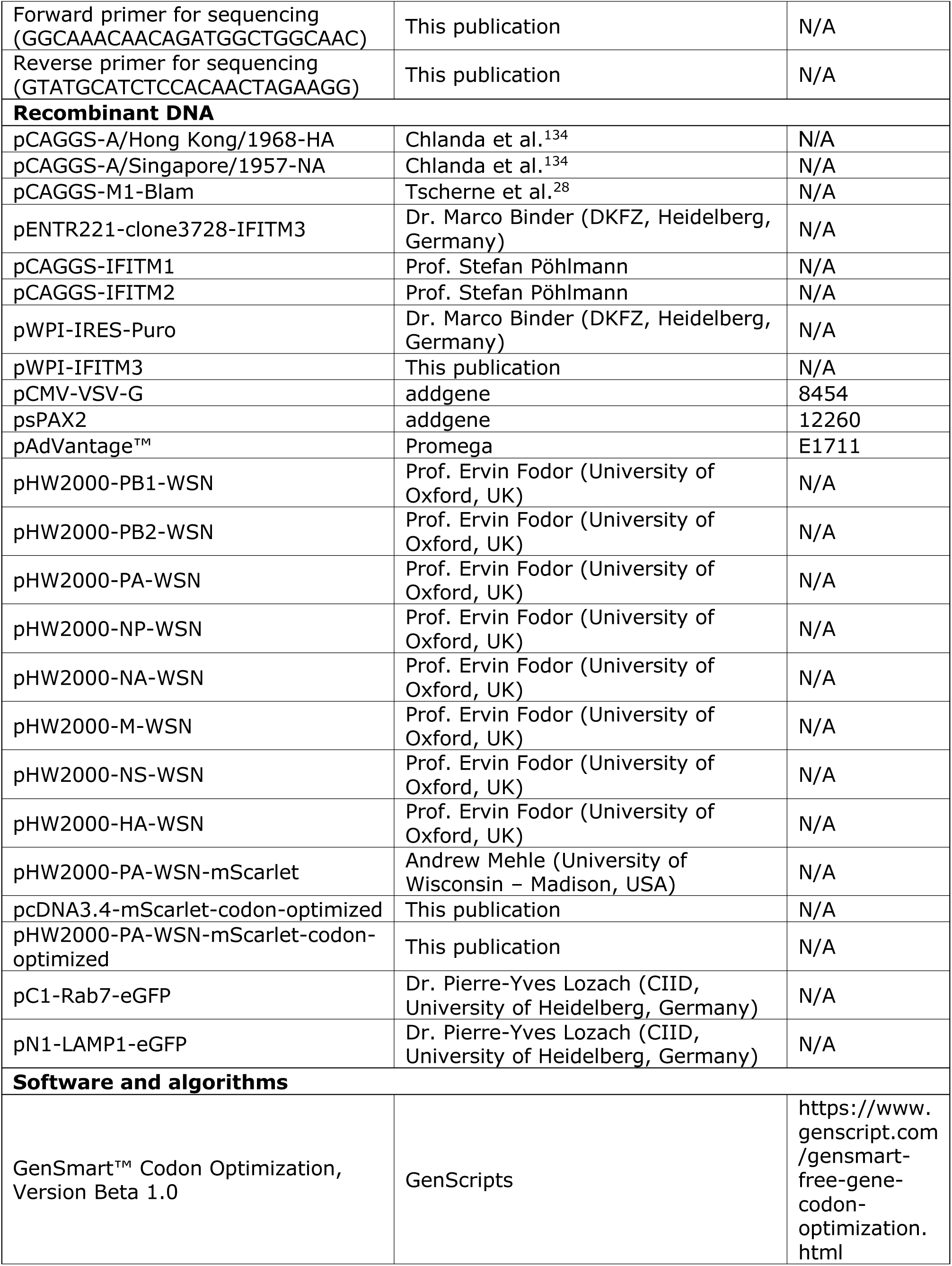
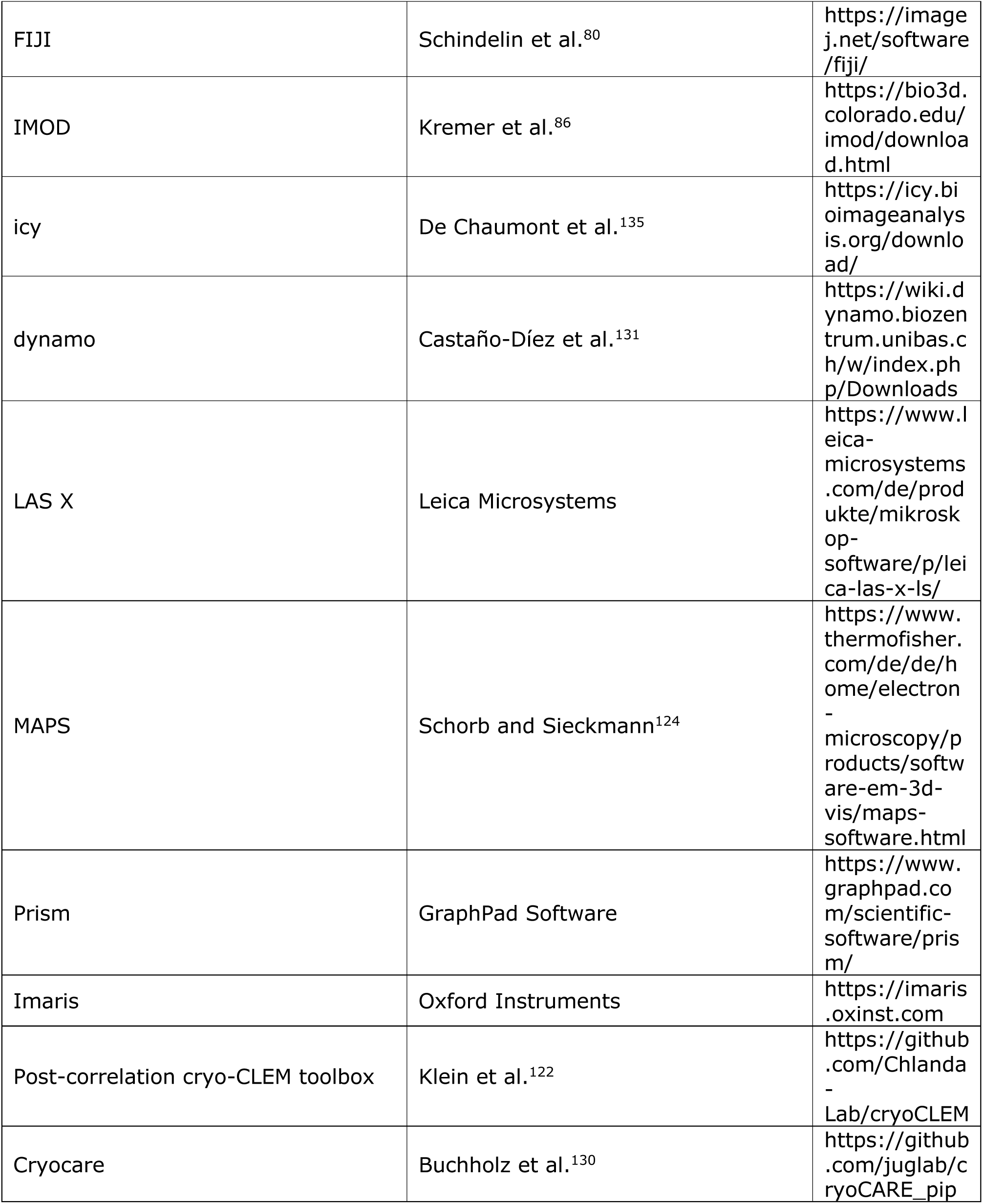
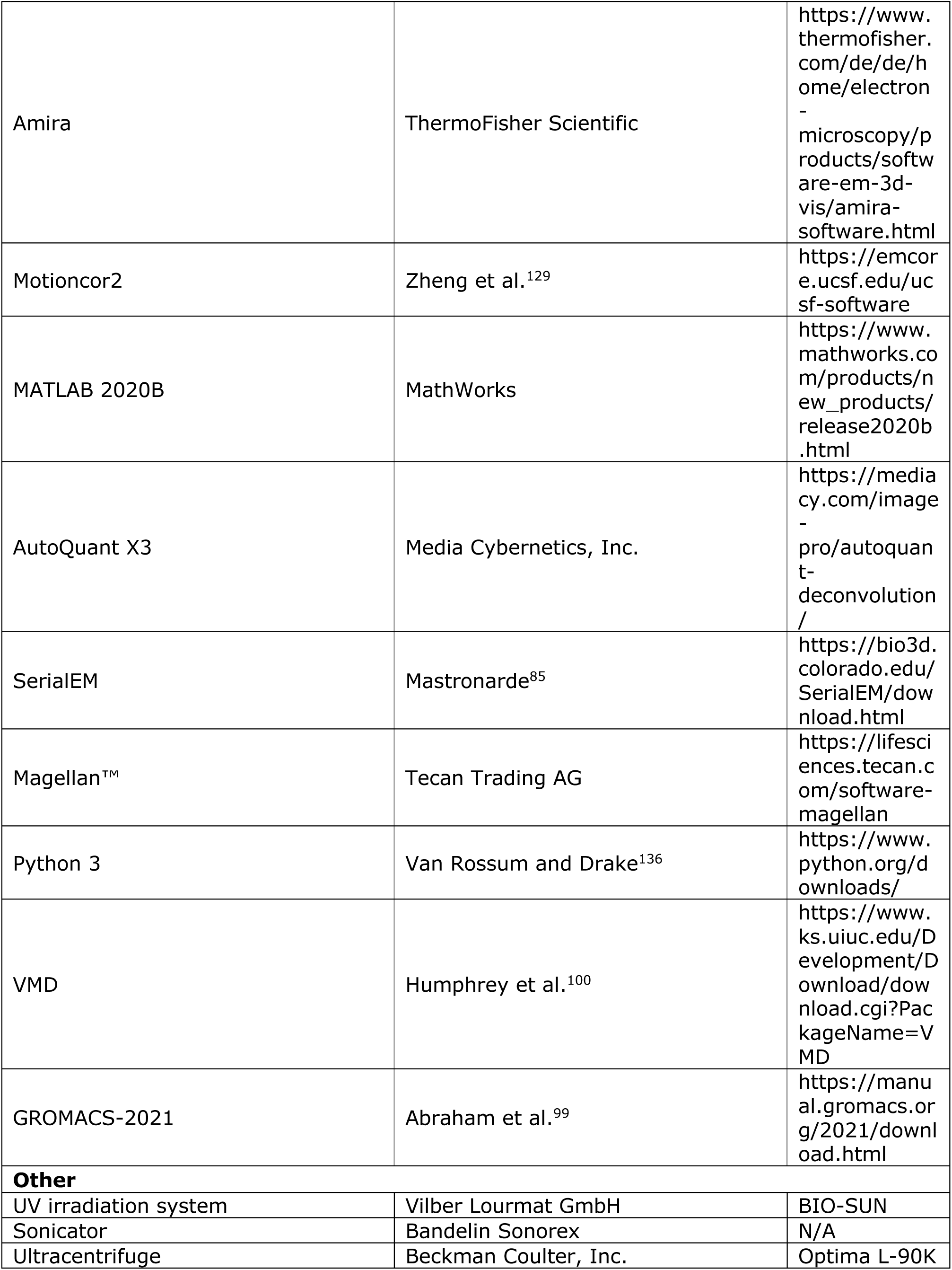
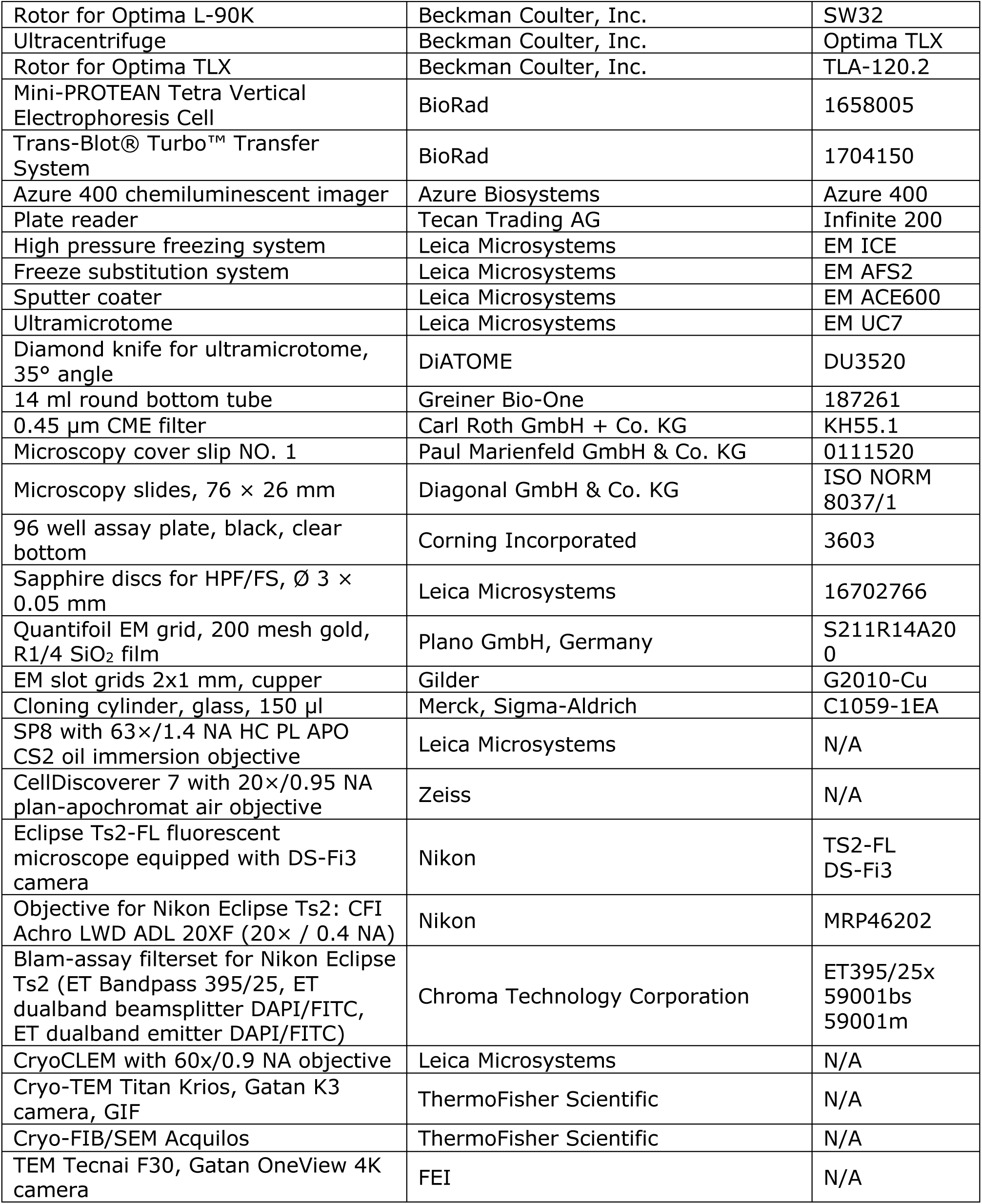

