## Supplemental Information for "IFITM3 blocks influenza virus entry by sorting lipids and stabilizing hemifusion"

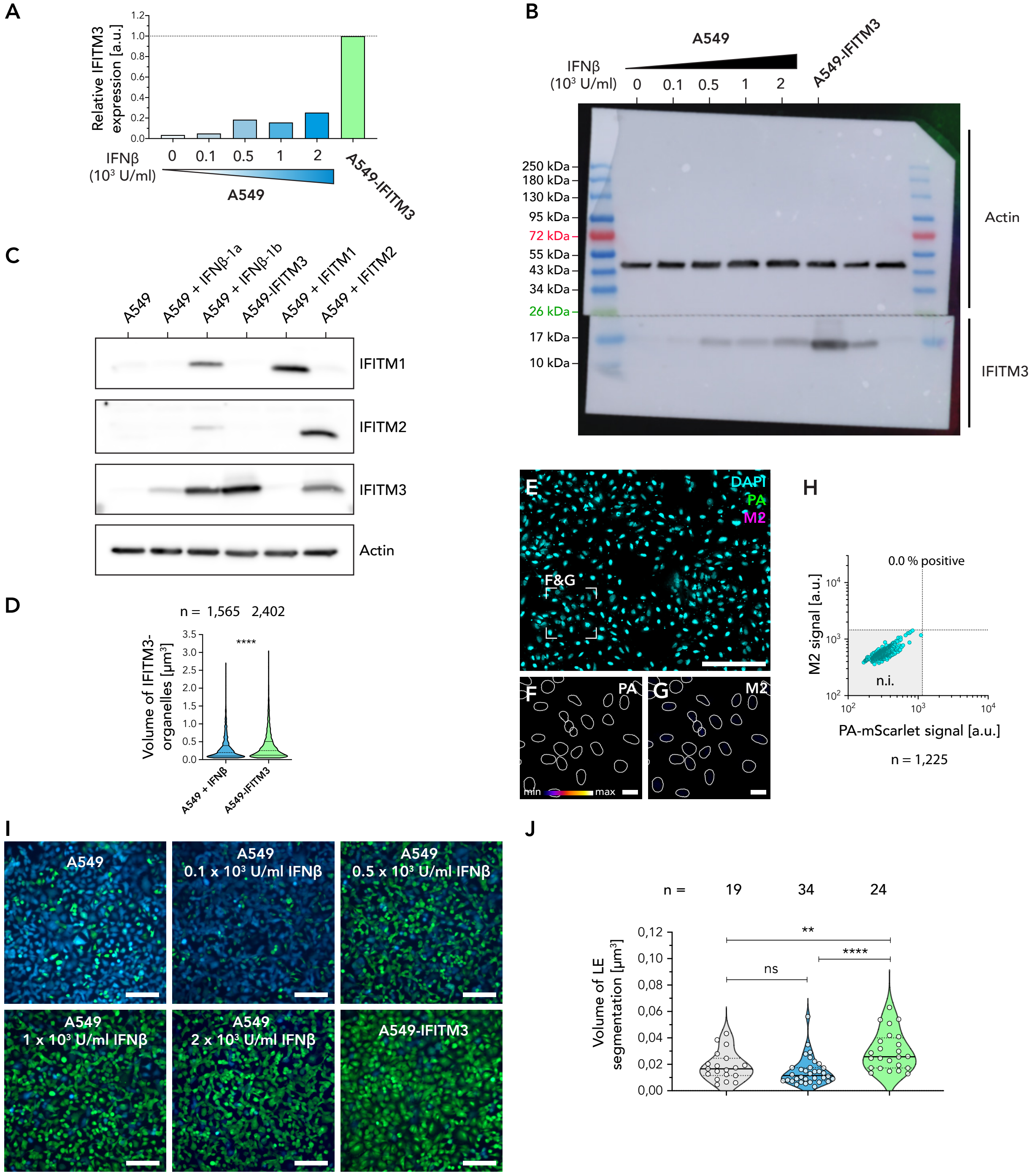


**Supplementary Figure 1 (related to Figure 1) Additional analysis of the antiviral properties of the A549-IFITM3 cell line.** **(A)** Quantification of the immunoblot analysis in Figure 1A. Relative expression levels of IFITM3 were normalized to actin. **(B)** Full image of western blot shown in Figure 1A. IFITM3 expression levels of A549-IFITM3 cells and A549 cells treated with IFNβ-1b (0, 0.1, 0.5, 1 and 2 × $10^3$ units/ml) were analyzed by immunoblot analysis. In the first and last lane of the blot, a broad range color prestained protein standard (New England BioLabs) was added. After western blotting, the membrane was cut at 26 kDa. The upper membrane section was immunolabelled for actin, the lower membrane section for IFITM3. **(C)** Immunoblot analysis of A549 (lane 1) and

A549-IFITM3 (lane 4) cells. As control, A549 cells were treated with 1 × $10^3$ units/ml INF β-1a (lane 2) or 1 × $10^3$ units/ml IFNβ-1b (lane 3) and transfected with IFITM1 (lane 5) or IFITM2 (lane 6). Actin was used as loading control. **(D)** Volume distribution of individual IFITM3 organelles based on three-dimensional segmentation of confocal microscopy data shown in Figure 1C, 1E and 1G. Data is represented as a violin plot, showing median (dotted line) and lower / upper quantiles (solid lines). An unpaired t-test was performed to evaluate the significance of differences between samples. **(E–H)** Negative control for the infection assay shown in Figure 1I–P. A549 cells were fixed and the viral protein M2 (shown in magenta) was labeled by immunofluorescence staining, and nuclei were labeled with DAPI (shown in cyan). **(E)** Composite overview image. **(F and G)** A magnified area (indicated as white square in E) for the signals of PA-mScarlet (F) and M2 are shown (G) using a fire lookup table. Individual nuclei were segmented (indicated with white lines), allowing for measuring the average signal intensity for PA-mScarlet and M2 in the segmented area. **(H)** For each segmented nucleus, the signal intensity for PA-mScarlet and M2 is shown as a scatter blot. Based on the maximum signal of this negative control, the two thresholds for non-infected cells (n.i.) of M1 ≤ 1,150 a.u. and PA-mScarlet ≤ 1,450 a.u. (dotted lines) were defined. **(I)** Fluorescent microscopy data of Blam-based cell entry assay (Figure 1Q) for A549 cells treated with increasing IFNβ-1b concentrations and A549-IFITM3 infected with M1-Blam influenza A VLPs. The signal of the non-cleaved FRET biosensor is shown in green and the signal of the proteolytically cleaved biosensor is in blue. **(J)** Quantification of segmented late endosomal volume of ET data (Figure 1S, 1U, 1W and Figure S2). Each data point represents a segmented tomogram. Scale bars: (E) 200 µm, (F and G) 20 µm, (I) 100 µm.

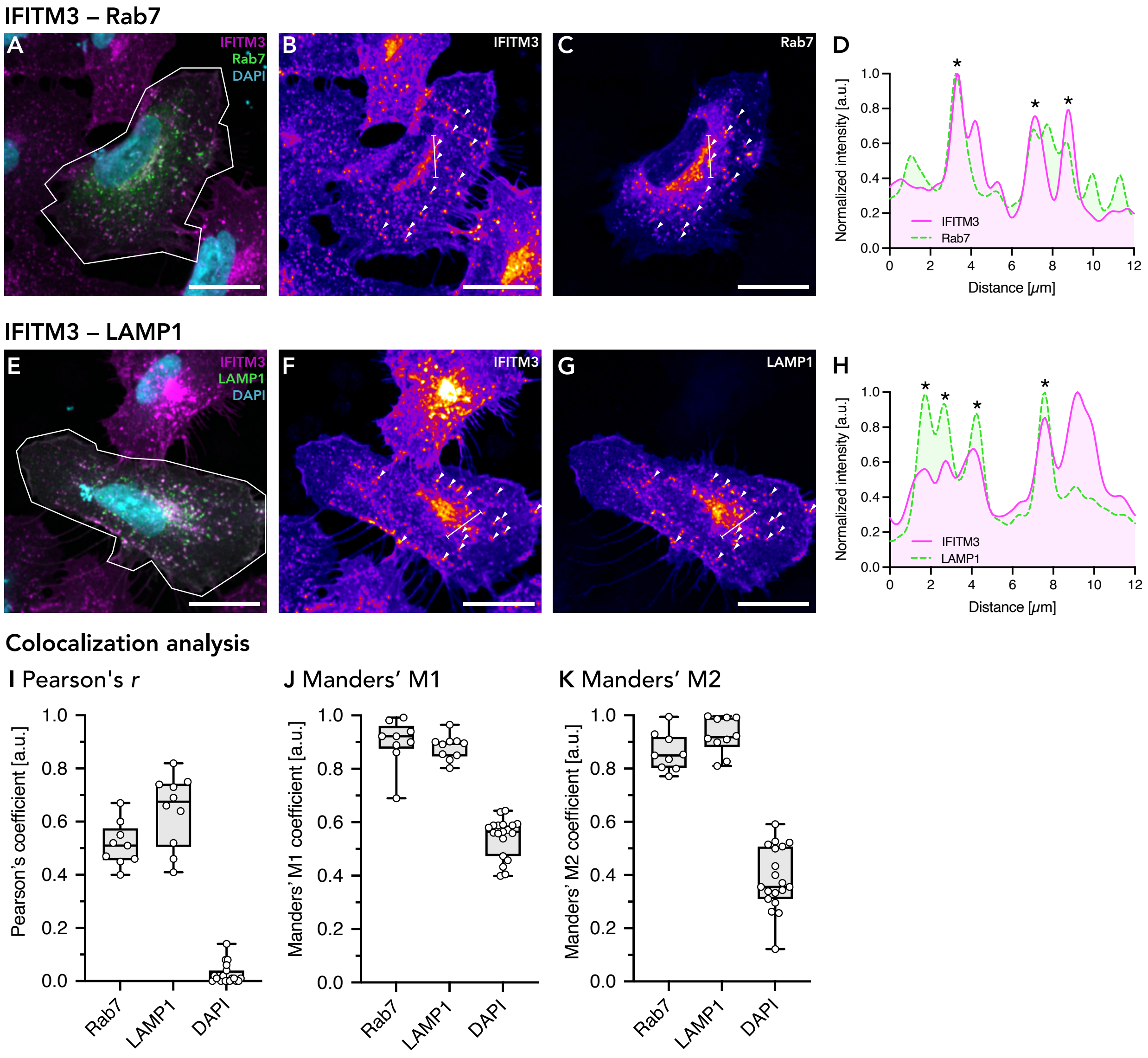


**Supplementary Figure 2 (related to Figure 1) IFITM3 colocalization with the endosomal marker Rab7, and the lysosomal marker LAMP1.** **(A–C)** Colocalization between IFITM3 and Rab7. A549-IFITM3 cells were transfected with Rab7-eGFP. IFITM3 was labeled by immunofluorescence, and nuclei were labeled with DAPI. A central slice of a 3D image volume is shown as a composite image (A) with IFITM3 in magenta, Rab7 in green, and DAPI in cyan. The region of interest (ROI) selected for colocalization analysis is indicated in white. The individual channels for IFITM3 (B) and Rab7 (C) are shown using a fire lookup table (LUT). Exemplary colocalizing features are indicated with white arrowheads. **(D)** Intensity line profiles for the IFITM3 and Rab7 signal. The position of the measurement is indicated as a white line in (B and C). Colocalizing features are indicated with an asterisk. **(E–G)** Colocalization between IFITM3 and LAMP1. A549-IFITM3 cells were transfected with LAMP1-eGFP. IFITM3 was labeled by immunofluorescence, and nuclei were labeled with DAPI. A central slice of a 3D image volume is shown as a composite image (E) with IFITM3 in magenta, LAMP1 in green, and DAPI in cyan. The region of interest (ROI) selected for colocalization analysis is indicated in white. The individual channels for IFITM3 (F) and LAMP1 (G) are shown using a fire lookup table (LUT). Exemplary colocalizing features are indicated with white arrowheads. **(H)** Intensity line profiles for the IFITM3 and LAMP1 signal. The position of the measurement is indicated as a white line (F and G). Colocalizing features are indicated with an asterisk. **(I–K)** Colocalization analysis between IFITM3 and Rab7, LAMP1 or DAPI using Pearson's correlation coefficient (I) and Manders' correlation coefficients (Manders et al. 1993) (J and K). For each sample (Rab7 or LAMP1), 3D volume data of 9 – 10 individual cells were analyzed. Scale bars: 20 µm.

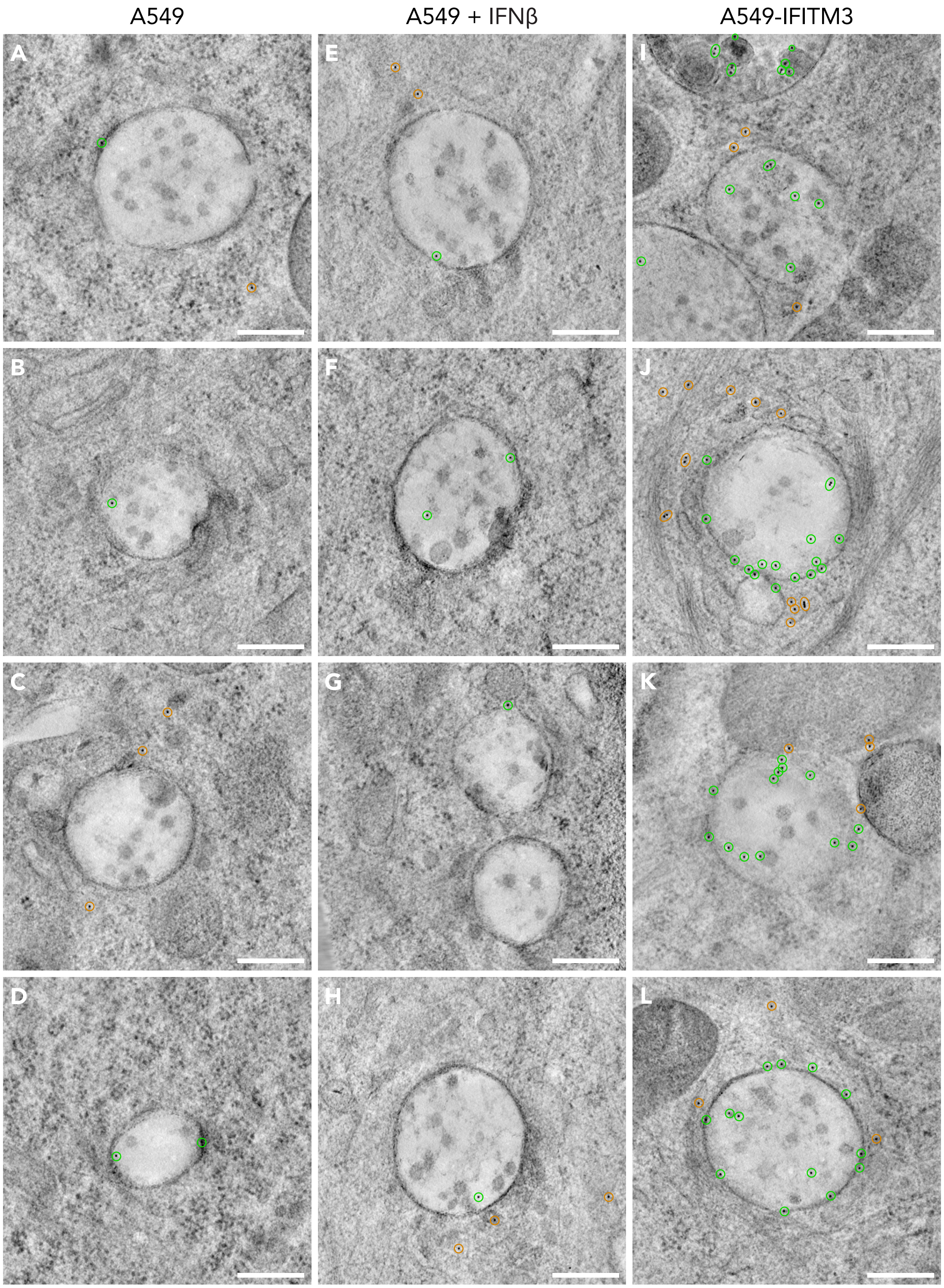


**Supplementary Figure 3 (related to Figure 1). Immuno-gold-labeling of late endosomes.** Tomograms of A549 cells (A–D), A549 cells treated with 2 × $10^3$ units/ml IFNβ-1b (E–H), and A549-IFITM3 cells (I–L) are shown. Each tomogram shows a representative late endosome featuring a typical multivesicular morphology characterized by ILVs in the late endosomal lumen. To visualize both the cellular ultrastructure and the immuno-gold-labeling against IFITM3, an average Z-projection of the whole tomogram was calculated. Gold particles are highlighted based on their localization: green circles indicate gold particles localized to late endosomes; orange circles represent gold particles localized in the cytoplasm. Scale bars: (A–L) 300 nm.

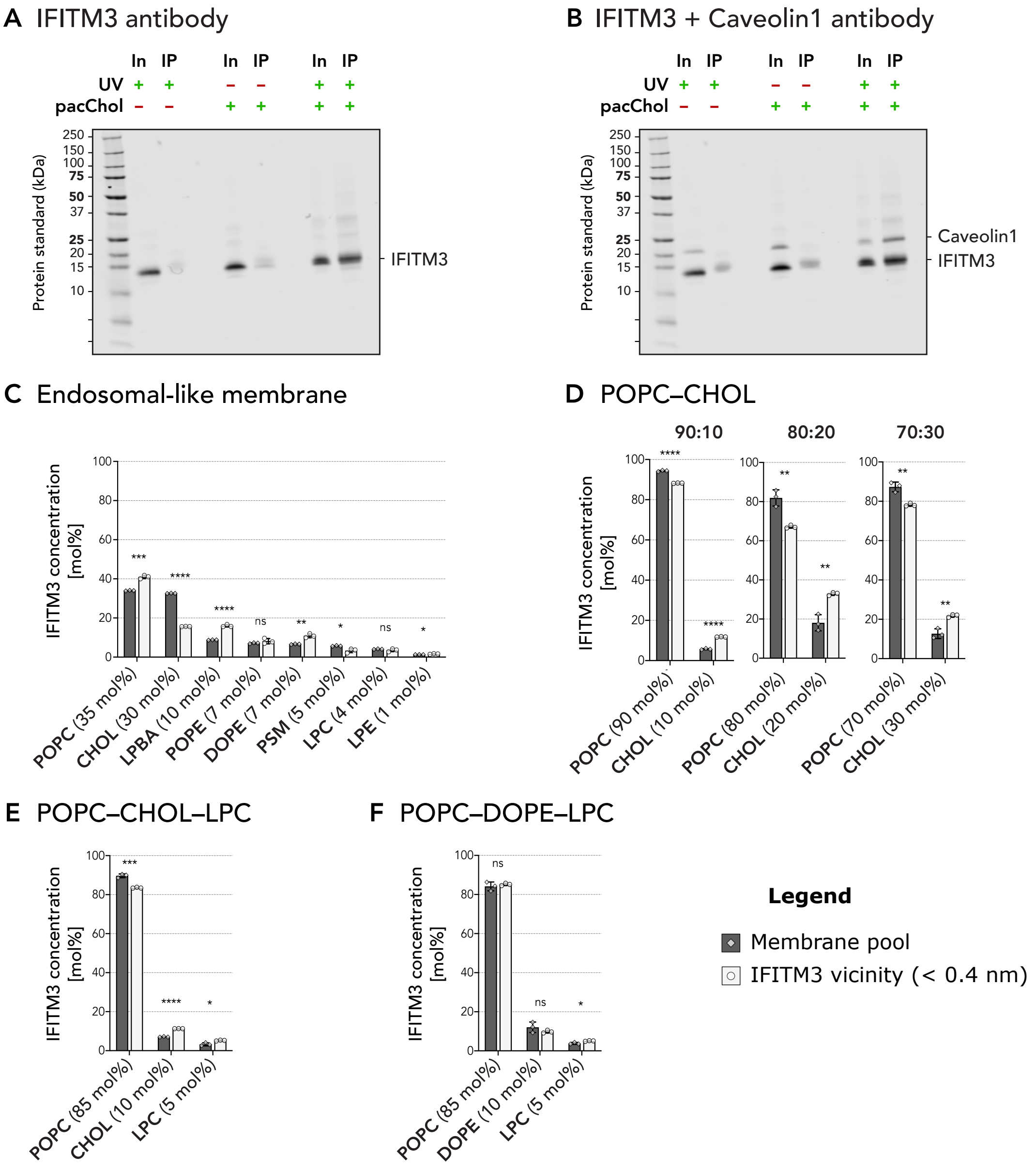


**Supplementary Figure 4 (Related to Figure 2) Additional analysis of molecular dynamics simulations. (A and B)** Full images of western-blot shown in Figure 2A. A549-IFITM3 cells were treated with a photo-reactive and clickable cholesterol analog (pacChol). After UV cross-linking and immunoprecipitation (IP), immunoblot analysis was performed. Complete cell lysate input (In) and immunoprecipitated sample (IP) were analyzed for conditions treated with UV and/or pacChol. The blot was first stained with an antibody against IFITM3 (A). The same blot was subsequently stained a second time with an antibody against Caveolin1 (B). In the first lane of the blot, the prestained protein standard 'Precision Plus' (Bio-Rad) was added. **(C–F)** Lipid concentrations in mol% in the vicinity of IFITM3 (< 0.4 nm) (white bars) and the surrounding membrane pool (dark bars) for different lipid systems: Late-endosomal like lipid system (C); binary POPC–cholesterol lipid system with cholesterol concentrations of 10, 20, and 30 mol% IFITM3 (D); ternary POPC–cholesterol–LPC lipid system (E); and ternary POPC–DOPE–LPC lipid system (F). Bars show the mean of three independent MD simulations, and the error bars represent the standard deviation (SD). All individual values are plotted. Unpaired t-tests were performed to evaluate the significance of differences between lipid concentrations in the IFITM3 vicinity and the surrounding membrane pool. Statistical significance: ns for non-significant, * for $p < 0.05$, ** for $p < 0.01$, *** for $p < 0.001$, **** for $p < 0.0001$.

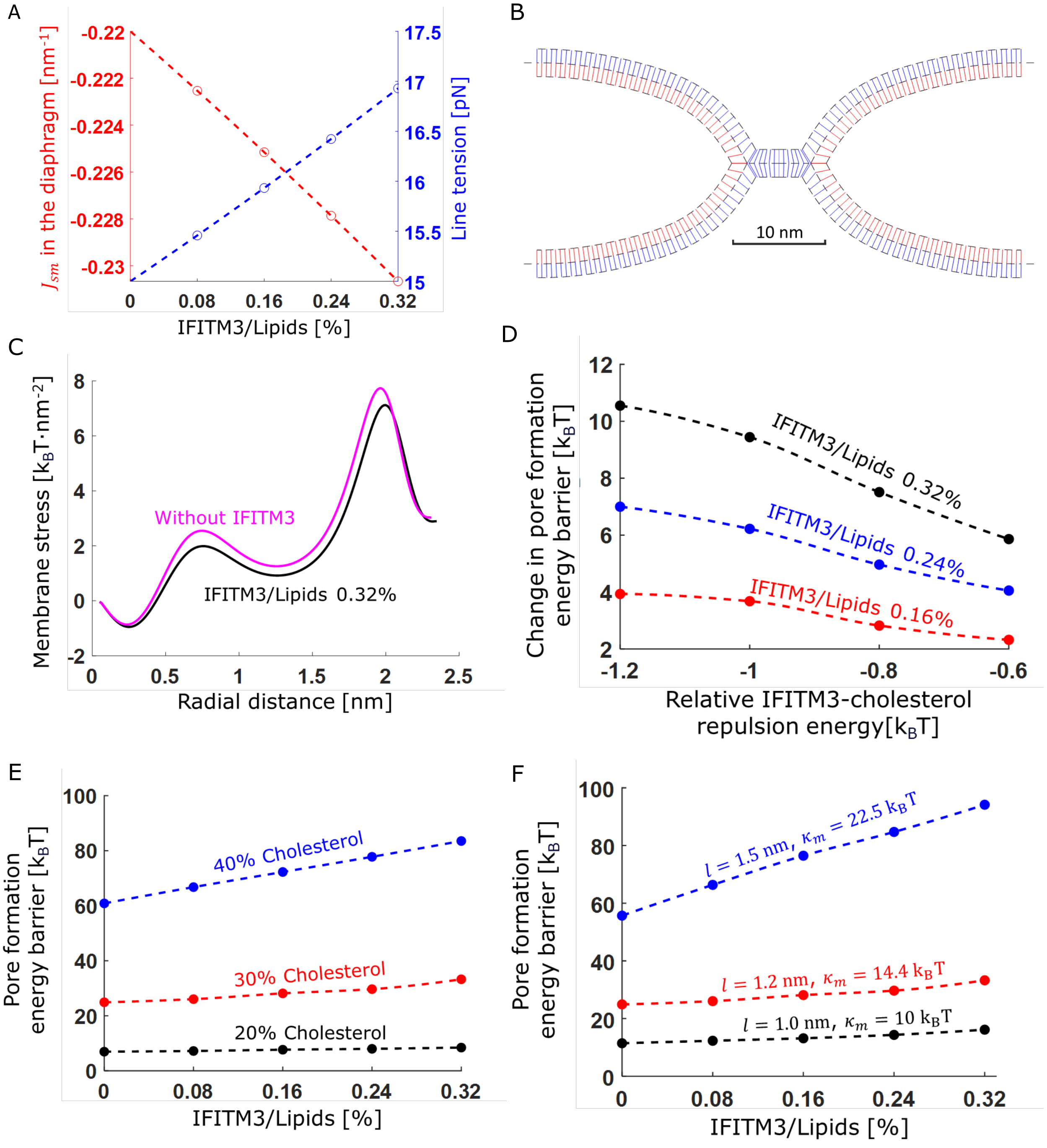


**Supplementary Figure 5 (Related to Figure 3) Additional Simulation results.** **(A)** Monolayer spontaneous curvature in the diaphragm and virus (red) and the pore line tension (blue) as a function of the ratio between lipids to IFITM3. **(B)** Hemifusion diaphragm cross-section shape after energy minimization before the addition of IFITM3. **(C)** Stress profile as a function of radial position, different cholesterol concentrations indicated by different colors. **(D)** The change in energy barrier as a function of cholesterol repulsion from IFITM3 at different IFITM3 mole fractions (indicated in legend) **(E-F)** Pore formation energy barrier as a function of IFITM3 mole fraction, **(E)** different baseline cholesterol levels, and **(F)** different tilt decay lengths, defined as $l = \sqrt{\kappa_m/\kappa_t}$. Parameters used in A-D: $\kappa_m = 14.4\ k_BT$, $\bar{\kappa}_m = -7.2\ k_BT$, $\delta_0 = 1.5\ nm$, $\zeta_{chol} = -0.5\ nm^{-1}$, $\zeta_0 = -0.1 nm^{-1}$ and $\lambda_0 = 15\ pN$, $\kappa_t = 40\ mN/m$ ($l = 1.2nm$) and 30% cholesterol mole fraction. (E) Cholesterol mole fraction indicated in plot. (F) Cholesterol 30%, monolayer bending rigidity, $\kappa_m$, indicated in plot. Since tilt rigidity is fixed, the tilt decay length changes as indicated. IFITM3 is inducing lipid sorting but no curvature in all panels.

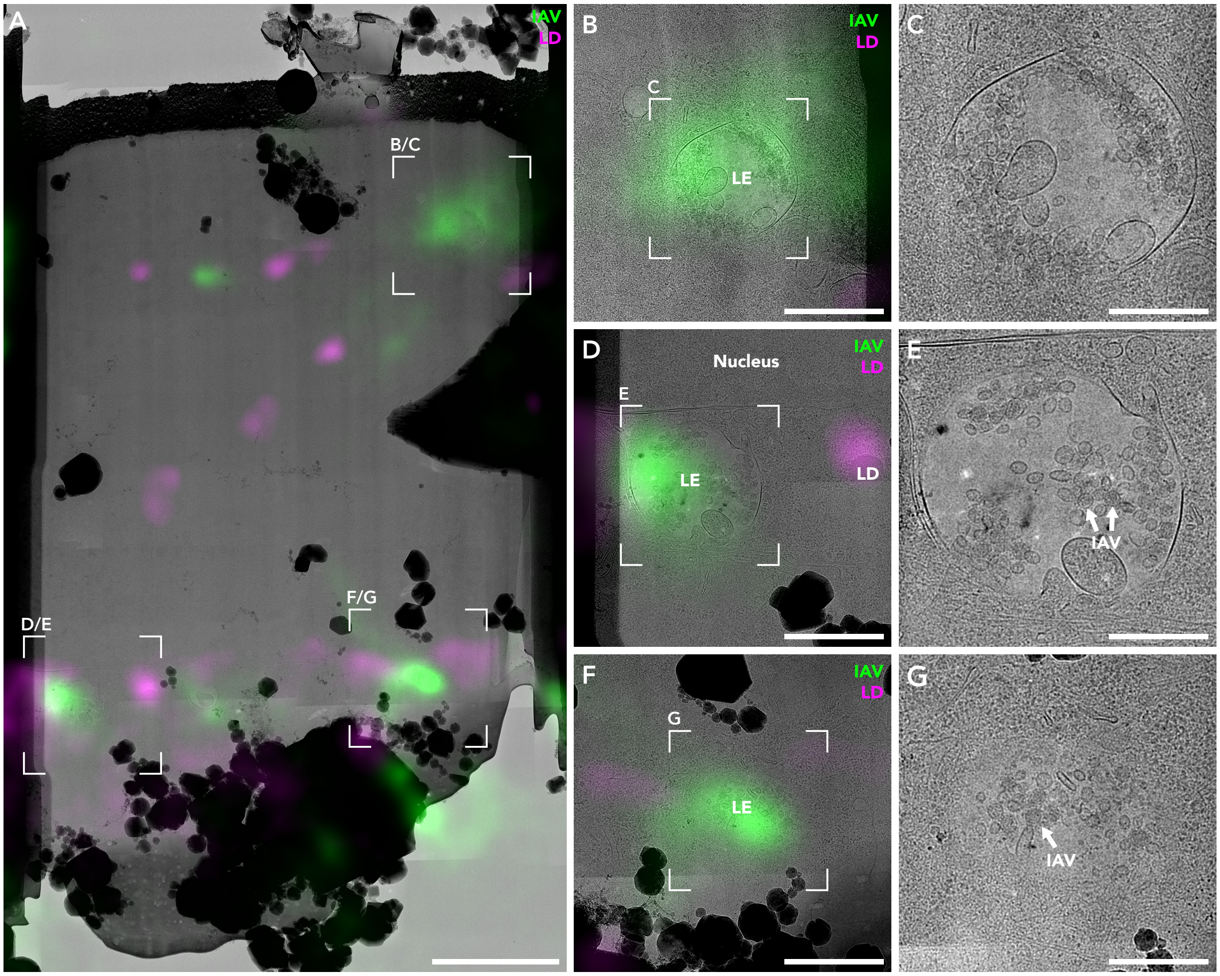


**Supplementary Figure 6 (related to Figure 4). Detailed view of *in situ* cryo-CLEM. (A)** Cryo-lamella of an A549-IFITM3 cell infected with fluorescently labeled influenza virus A/WSN/1933-nDio (MOI = 200, 1 hpi). Lipid droplets (LDs) were fluorescently labeled and used as fiducial markers for in situ cryo-CLEM. The fluorescent signal of IAV is shown in green. The LD signal is shown in magenta. **(B, D and F)** Magnified areas of the correlated cryo-lamella (A). Late endosomes (LE) with a typical multivesicular morphology were positive for fluorescent IAV signal. One correlated LD is shown in (C). **(C, D and G)** Magnified areas of the correlated cryo-lamellae (B, D and F) without overlaid fluorescence signal. Individual viral particles are indicated (white arrows). Scale bars: (A) 3 µm, (B, D, and F) 1 µm, (C, E, and G) 500 nm.

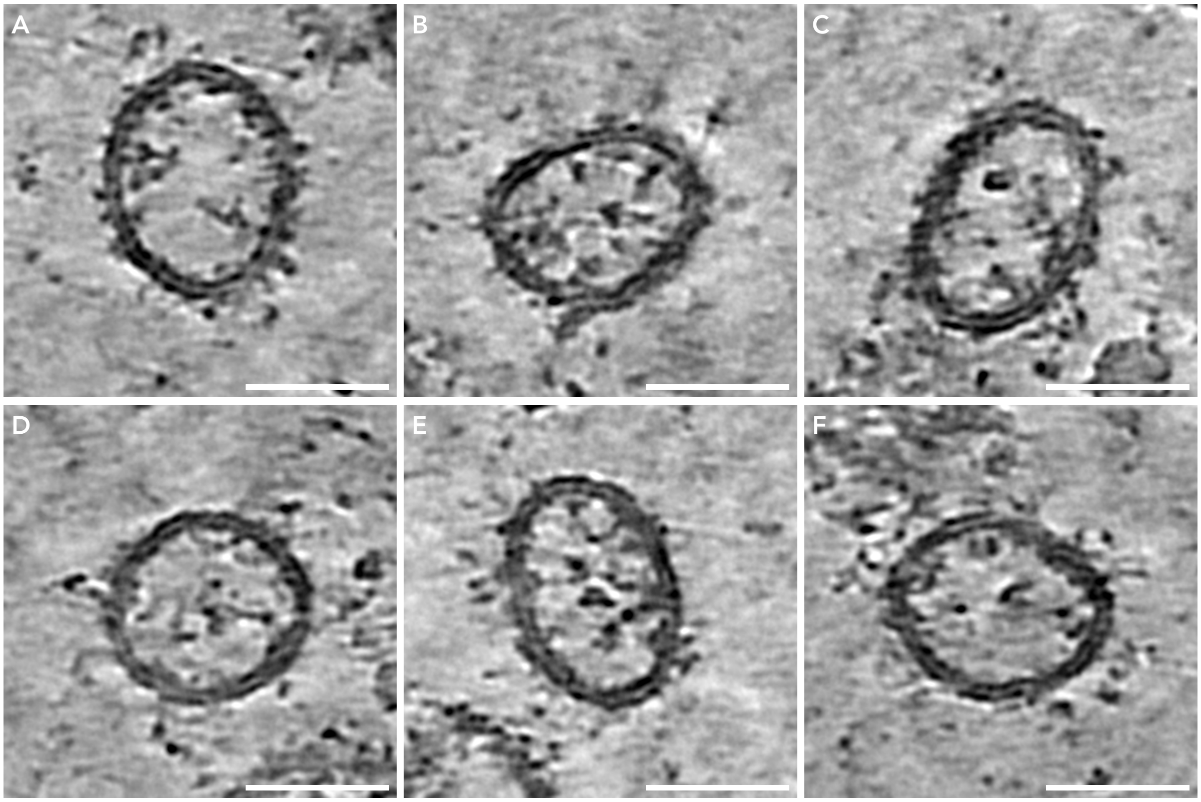


**Supplementary Figure 7 (related to Figure 4). Tomograms of ILVs in the late endosomal lumen.** Representative tomographic slices (5.34 nm thickness) of ILVs in the late endosomal lumen. ILV membranes show a typical protein coating. To increase the signal-to-noise ratio, reconstructed tomograms were SIRT-like filtered and denoised using cryoCARE. Scale bars: (A–F) 50 nm.

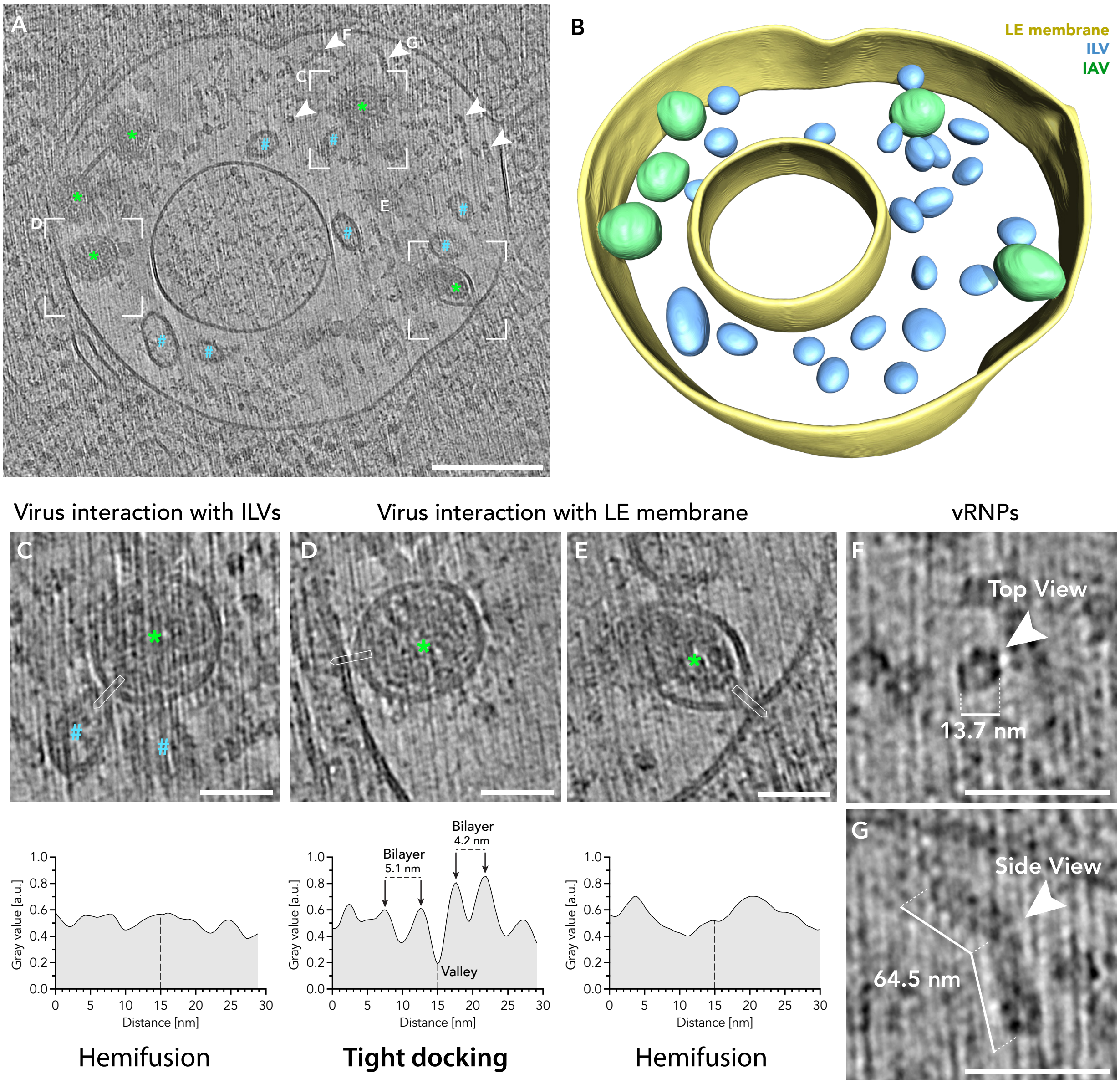


**Supplementary Figure 8 (related to Figure 4). Tomograms of ILVs in the late endosomal lumen. (A)** Tomographic slice (5.34 nm thickness) of a late endosome of A549-IFITM3 cells infected with fluorescently labeled influenza virus A/WSN/1933-nDiO (MOI = 200, 1 hpi). The late endosomal lumen incorporates ILVs (blue hashes) and IAV particles (green asterisks). vRNPs in the endosomal lumen are indicated with white arrowheads. To increase the signal-to-noise ratio, reconstructed tomograms were SIRT-like filtered and denoised using cryoCARE. **(B)** The three-dimensional volume segmentation of the tomogram shown in (A). The late endosomal membrane is shown in yellow, IAV particles in green, and ILVs in blue. **(C–E)** Magnified areas of reconstructed tomogram (A) showing one exemplary stabilized hemifusion sites between IAV particles and ILVs (C) and two contact sites between IAV particles and the limiting late endosomal membrane (D and E). Virus particles are indicated with green asterisks and ILVs with blue hashes. The contact sites were analyzed by a linear density profile. The position of the line profile is indicated with a white square. Data was acquired using a line width of 5.34 nm. The line profile is blotted (bottom column), and features are annotated. Based on the line profile, contact sites were classified as tight docking or hemifusion. **(F**

**and G)** Detailed view of vRNPs in the endosmal lumen shown as top view (F) and side view (G). See Video S2 for three-dimensional rendering. Scale bars: (A) 200 nm, (C–F) 50 nm.

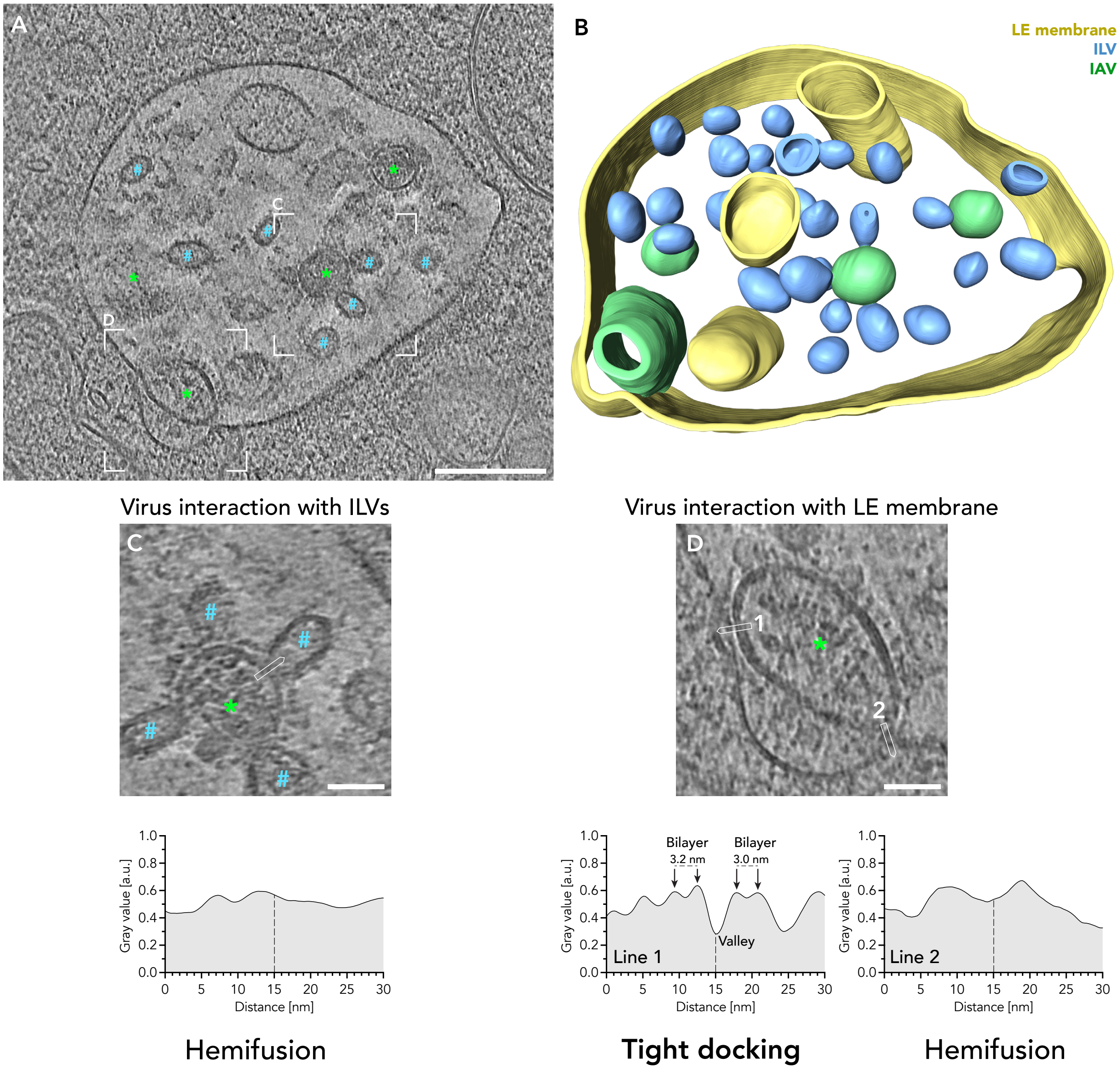


**Supplementary Figure 9 (related to Figure 4). Tomogram of ILVs in the late endosomal lumen. (A)** Tomographic slice (5.34 nm thickness) of a late endosome of A549-IFITM3 cells infected with fluorescently labeled influenza virus A/WSN/1933-nDiO (MOI = 200, 1 hpi). The late endosomal lumen incorporates ILVs (blue hashes) and IAV particles (green asterisks). To increase the signal-to-noise ratio, reconstructed tomograms were SIRT-like filtered and denoised using cryoCARE. **(B)** The three-dimensional volume segmentation of the tomogram shown in (A). The late endosomal membrane is shown in yellow, IAV particles in green, and ILVs in blue. **(C and D)** Magnified areas of reconstructed tomogram (A) showing one stabilized hemifusion site between an IAV particle and ILVs (C) and between an IAV particle and the limiting late endosomal membrane (D). Virus particles are indicated with green asterisks and ILVs with blue hashes. The contact sites were analyzed by a linear density profile. The position of the line profile is indicated with a white square. Data was acquired using a line width of 5.34 nm. The line profile is blotted (bottom column), and features are annotated. Based on the line profile, contact sites were classified as tight docking or hemifusion.See Video S3 for three-dimensional rendering. Scale bars: (A) 200 nm, (C–F) 50 nm.

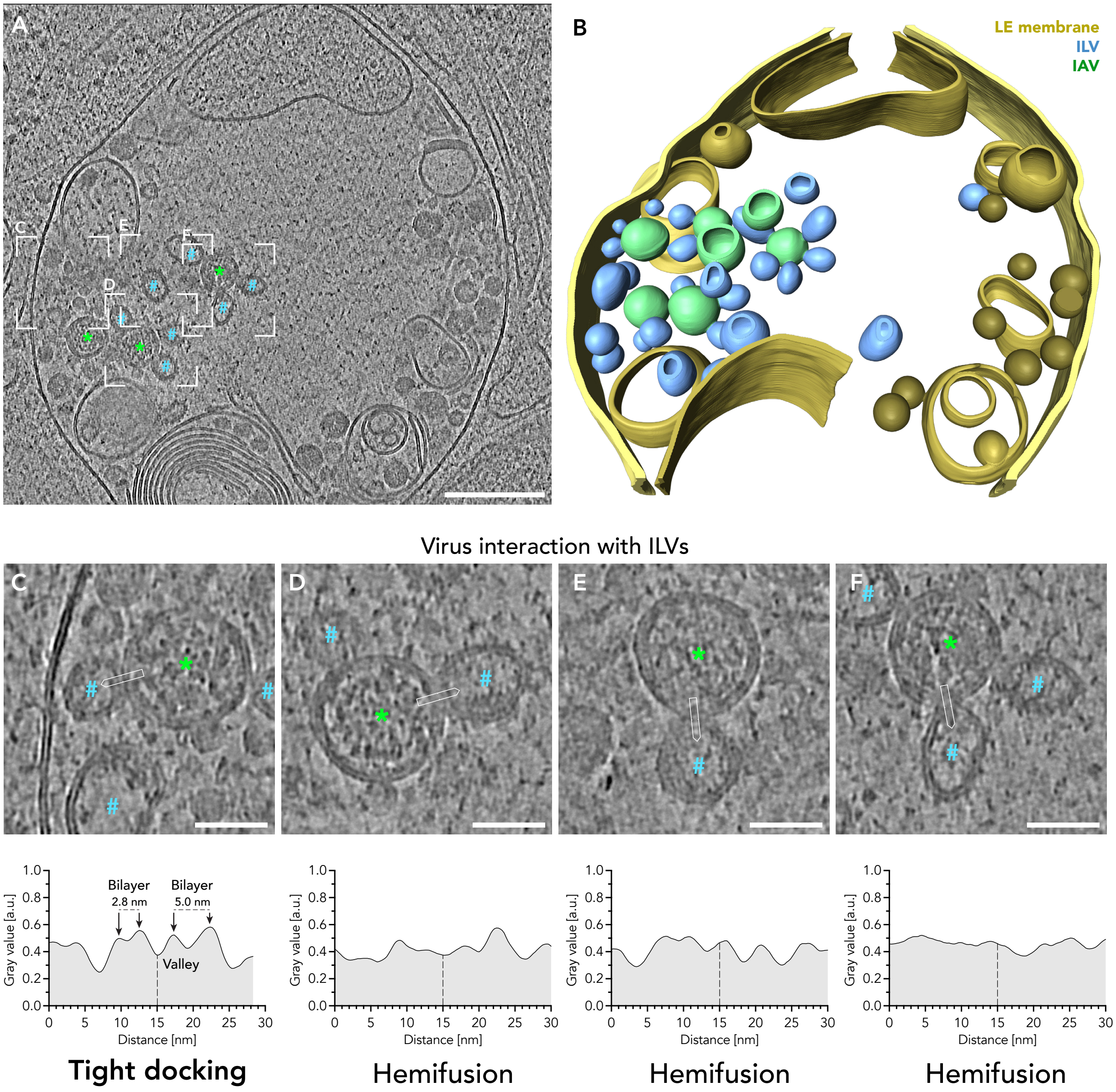


**Supplementary Figure 10 (related to Figure 4). Tomogram of ILVs in the endosomal lumen. (A)** Tomographic slice (5.34 nm thickness) of an endosome of A549-IFITM3 cells infected with fluorescently labeled influenza virus A/WSN/1933-nDiO (MOI = 200, 1 hpi). The endosomal lumen incorporates ILVs (blue hashes), IAV particles (green asterisks), and various cellular membrane degradation products. To increase the signal-to-noise ratio, reconstructed tomograms were SIRT-like filtered and denoised using cryoCARE. **(B)** The three-dimensional volume segmentation of the tomogram shown in (A). The endosomal membrane is shown in yellow, IAV particles in green, and ILVs in blue. **(C–F)** Magnified areas of reconstructed tomogram (A) showing stabilized hemifusion sites between IAV particles and ILV. Virus particles are indicated with green asterisks and ILVs with blue hashes. The contact sites were analyzed by a linear density profile. The position of the line profile is indicated with a white square. Data was acquired using a line width of 5.34 nm. The line profile is blotted (bottom column), and features are annotated. Based on the line profile, contact sites were classified as tight docking or hemifusion.See Video S4 for three-dimensional rendering. Scale bars: (A) 200 nm, (C–F) 50 nm.

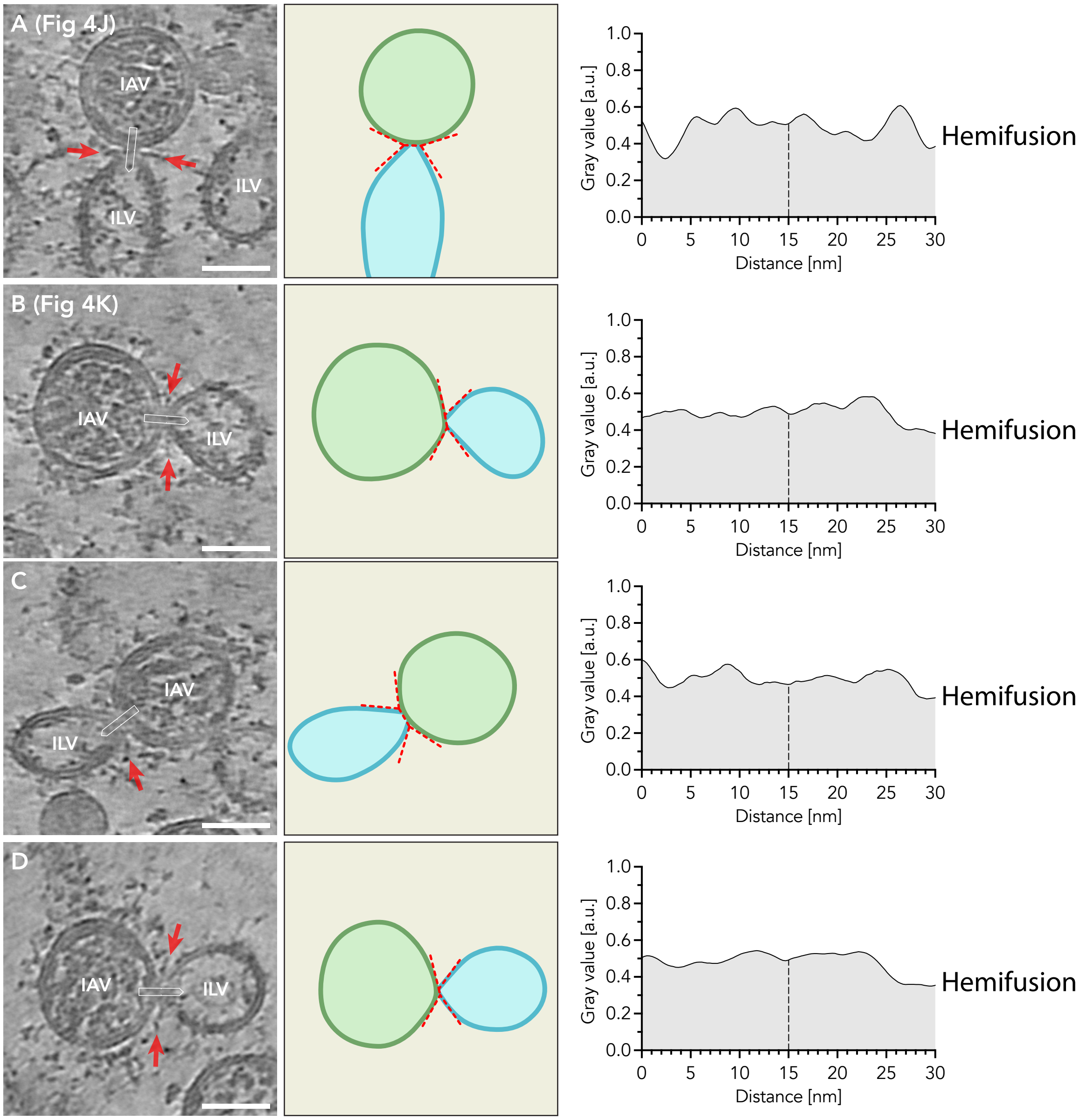


**Supplementary Figure 11 (related to Figures 4 and 5). IAV particles interact with intraluminal vesicles in the late endosomal lumen.** See page 17 for captions.

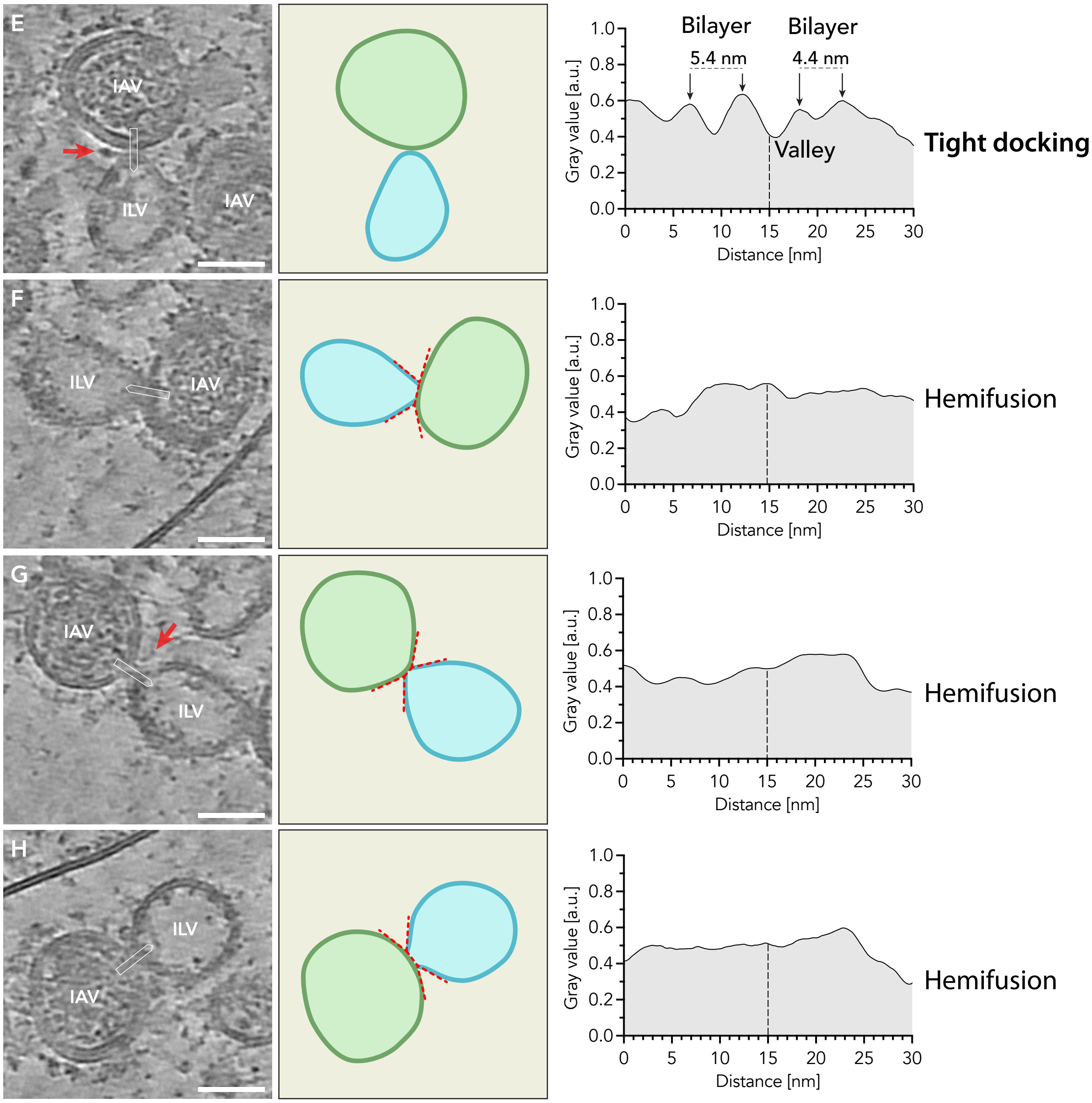


**(continued) Supplementary Figure 11 (related to Figures 4 and 5). IAV particles interact with intraluminal vesicles in the late endosomal lumen.** See page 17 for captions.

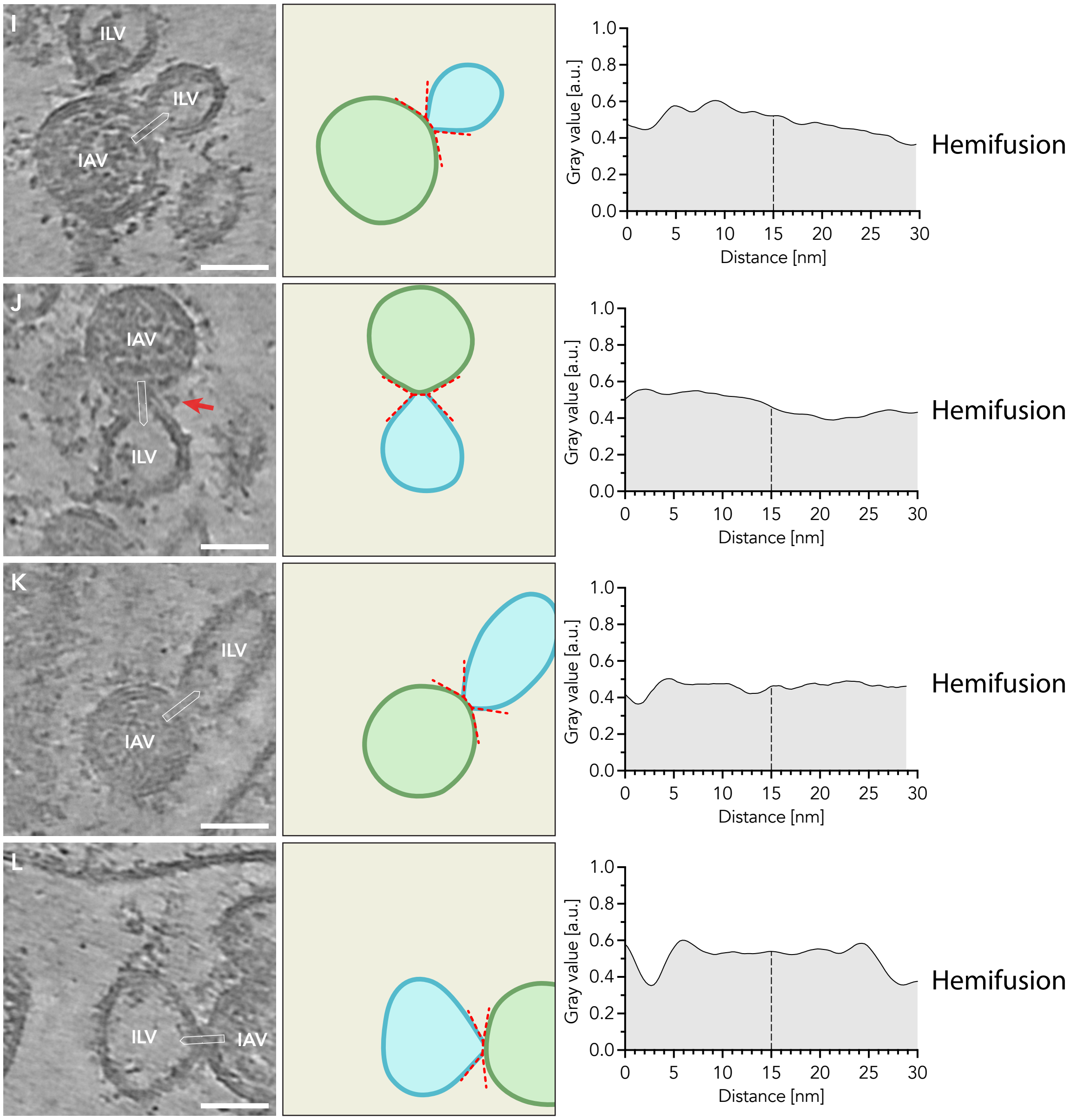


**(continued) Supplementary Figure 11 (related to Figures 4 and 5). IAV particles interact with intraluminal vesicles in the late endosomal lumen.** See page 17 for captions.

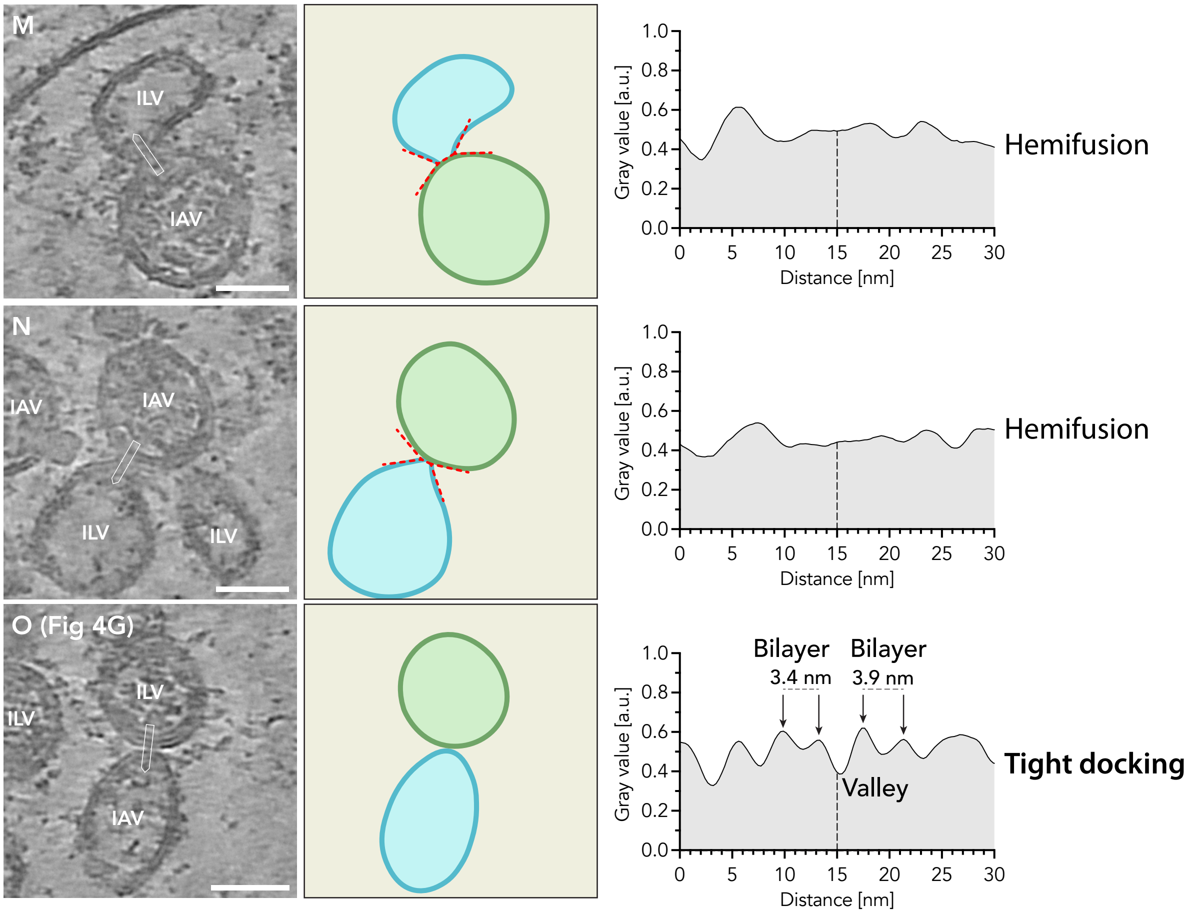


**(continued) Supplementary Figure 11 (related to Figure 4 and 5). IAV particles interact with intraluminal vesicles in the late endosomal lumen.** Gallery of all hemifusion sites between viral particles (IAV) and intraluminal vesicles (ILVs) in the late endosomal lumen of the tomogram shown in Figure 4E and 4F. Post-fusion HA glycoproteins close to the hemifusion sites are indicated with a red arrow. For each site, a schematic drawing is shown (center column), with the viral particle in green and the ILV in blue. The red dotted lines indicate the measurements for the hemifusion diaphragm lengths and the hemifusion angles (shown in Figure 5A–C). Based on this measured geometry, the stress in each hemifusion diaphragm and the fusion-pore formation energy was calculated as indicated in the schematic drawings. The contact site was analyzed by a linear density profile. The position of the line profile is indicated with a white square. Data was acquired using a line width of 5.34 nm. The line profile is blotted (right column), and features are annotated. Based on the line profile, contact sites were classified as tight docking or hemifusion. Scale bars: (A–N) 50 nm.

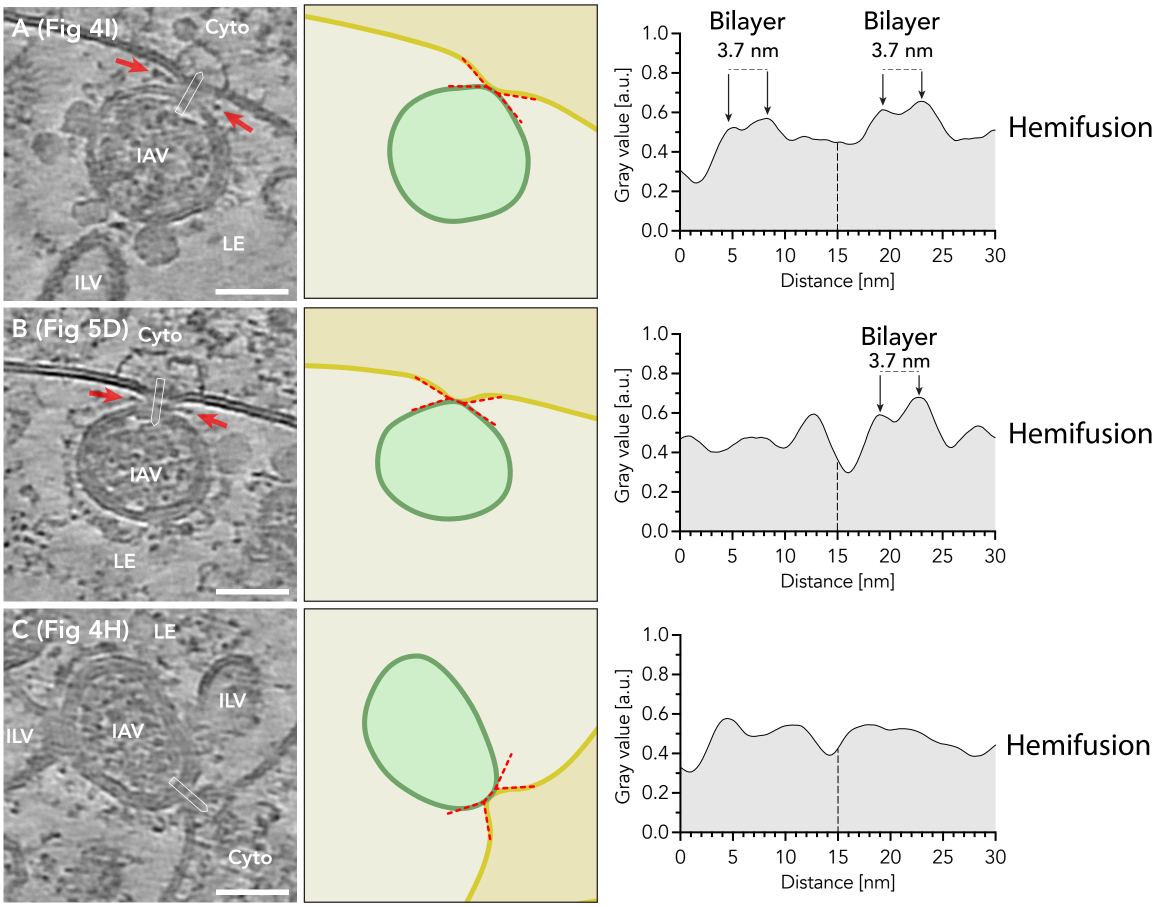


**Supplementary Figure 12 (related to Figure 4 and 5). IAV particles interact with limiting late endosomal membrane.** Gallery of all hemifusion sites between viral particles (IAV) and the limiting late endosomal membrane of the tomogram shown in Figure 4E and 4F. Post-fusion HA glycoproteins close the the hemifusion sites are indicates with a red arrow. For each site, a schematic drawing is shown on the right with the viral particle in green and the limiting late endosomal membrane in yellow. The red dotted lines indicate the measurements for the hemifusion diaphragm lengths and the hemifusion angles (shown in Figure 5A–C). Based on this measured geometry, the stress in each hemifusion diaphragm and the fusion-pore formation energy was calculated as indicated in the schematic drawings. Scale bars: (A–N) 50 nm. The contact site was analyzed by a linear density profile. The position of the line profile is indicated with a white square. Data was acquired using a line width of 5.34 nm. The line profile is blotted (right column), and features are annotated. Based on the line profile, contact sites were classified as tight docking or hemifusion. Scale bars: (A–N) 50 nm.

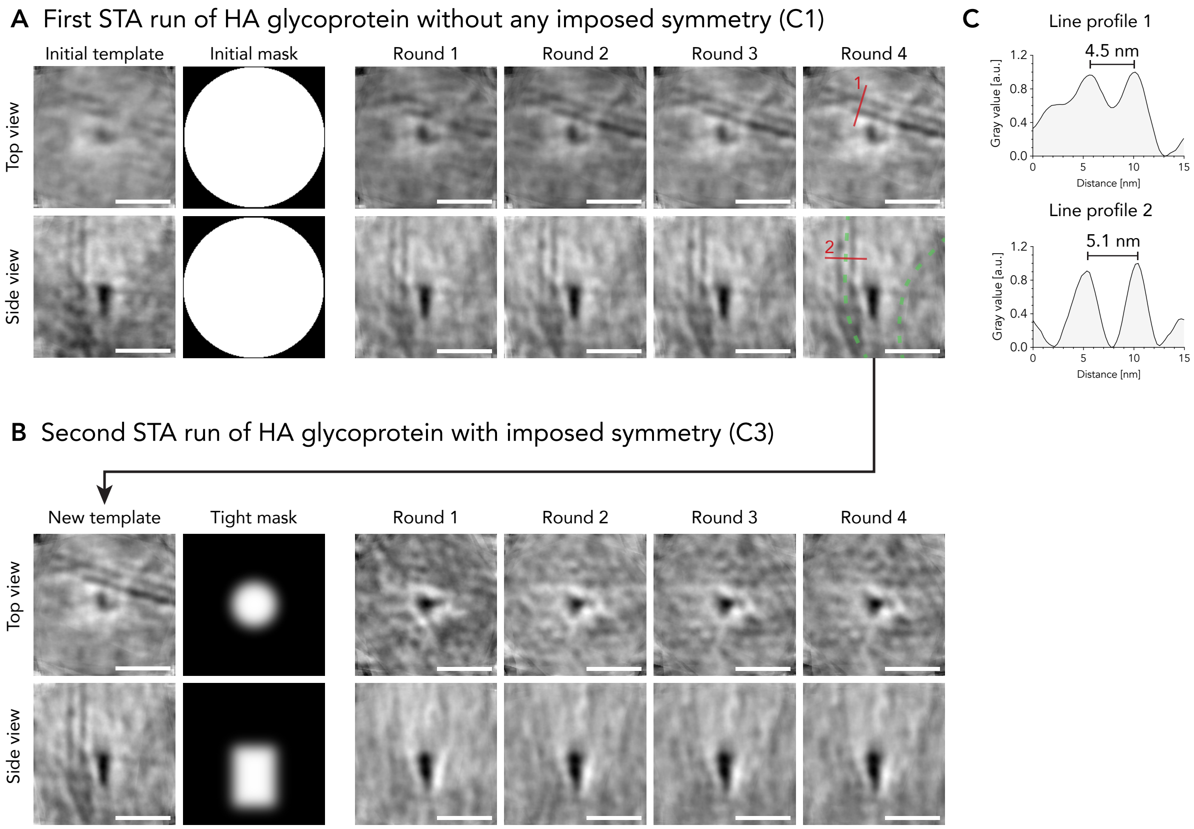


**Supplementary Figure 13 (related to Figure 5). Subtomogram average iterations.** **(A)** First subtomogram average (STA) run. No symmetry was imposed (C1). The initial template was generated by averaging all picked particles using a dipole-model which provides initial orientation of each particle. As a mask, a generic sphere was used. The particles were averaged over 4 rounds. The result of each round is shown as central slice through the averaged volume as a top and side view. In both the top and side view of the result of the last round, in addition to the central density of the post-fusion HA, the phospholipid bilayer of the viral and late endosomal membrane (green dotted lines) is resolved. The phospholipid monolayer distance was measured at the two indicated sites (red lines). **(B)** Plotted line profiles (A). For each profile the two maxima (representing the center of each phospholipid monolayer) were determined, and the distance was measured as indicated. **(C)** Second STA run, refining the results of the first STA run (A). The final result of the first run was used as new template. To refine the resolution of the post-fusion HA, a tight mask was used. As HA is a trimeric protein, a three-fold symmetry (C3) was imposed. The particles were averaged over 4 rounds. The result of each round is shown as central slice through the averaged volume as a top and side view. Scale bars: (A and B) 10 nm.

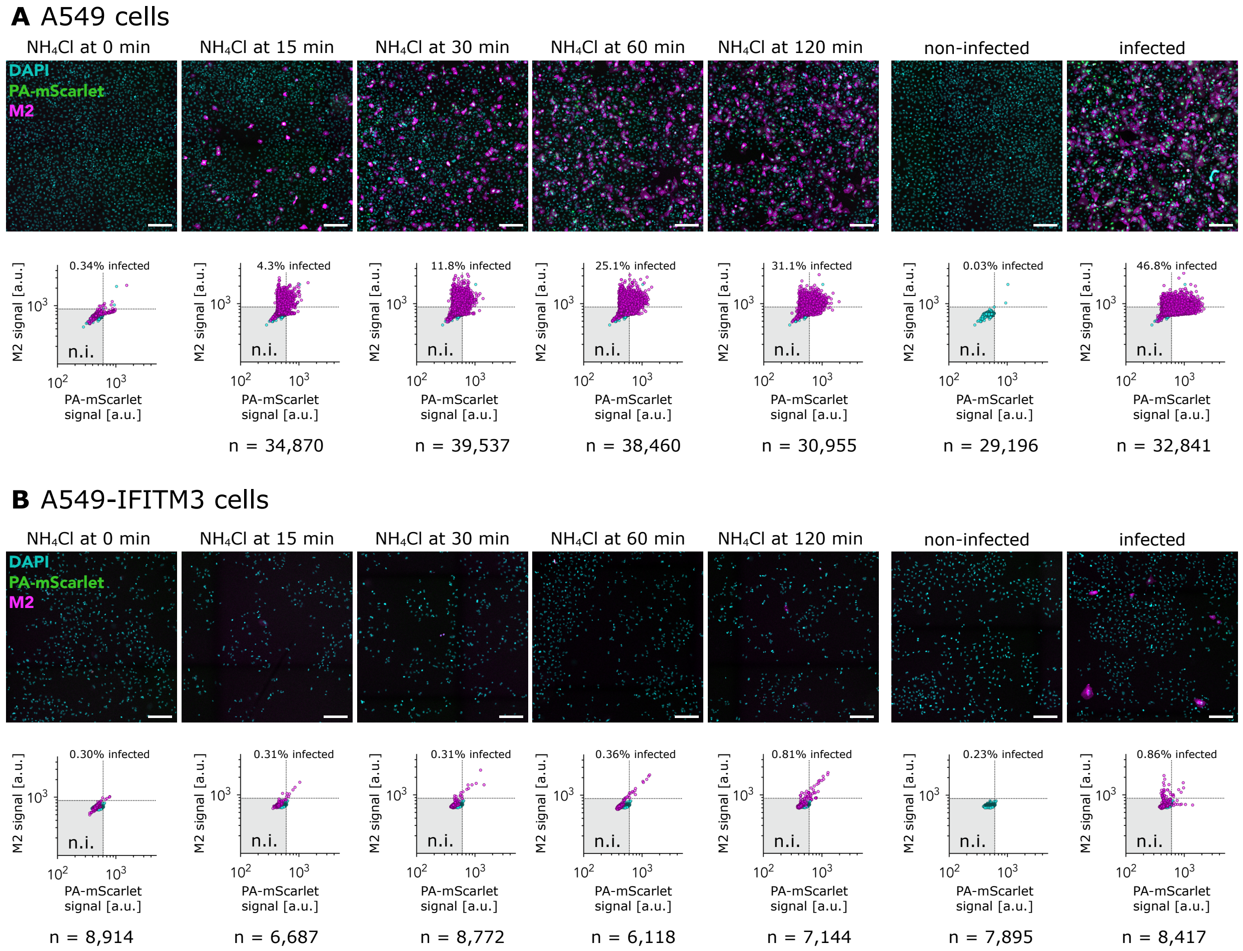


**Supplementary Figure 14 (related to Figure 6). IAV penetration assay by $NH_4Cl$ add-in time course.** A549 cells (A), and A549-IFITM3 cells (B) were infected with a fluorescent reporter virus A/WSN/1933-PA-mScarlet (MOI = 3). At different time points post infection (0 mpi, 15 mpi, 30 mpi, 60 mpi and 120 mpi) cells were treated with 50 mM $NH_4Cl$, inhibiting viral membrane fusion by neutralization of the endosomal system. Cells were fixed 12 hpi, M2 was labeled by immunofluorescence, and nuclei were fluorescently labeled by DAPI. For each sample a large area was acquired using an automated fluorescence microscopy Celldiscoverer 7 (Zeiss). Nuclei were automatically segmented and the average signal for M2 and PA-mScarlet was measured in the segmented area. For A549 cells, between 29,196 and 39,537 cells were segmented, for A549-IFITM3 cells, between 6,118 and 8,914 cells. The top row shows an exemplary section of the sample with DAPI in cyan, PA-mScarlet in green and M2 in magenta. In the bottom row, a scatter plot for the PA-mScarlet and M2 signal is shown. Thresholds for defining infected cells were manually set based on the non-infected controls, as indicated with dotted lines. For each condition, the percentage of infected cells is indicated.

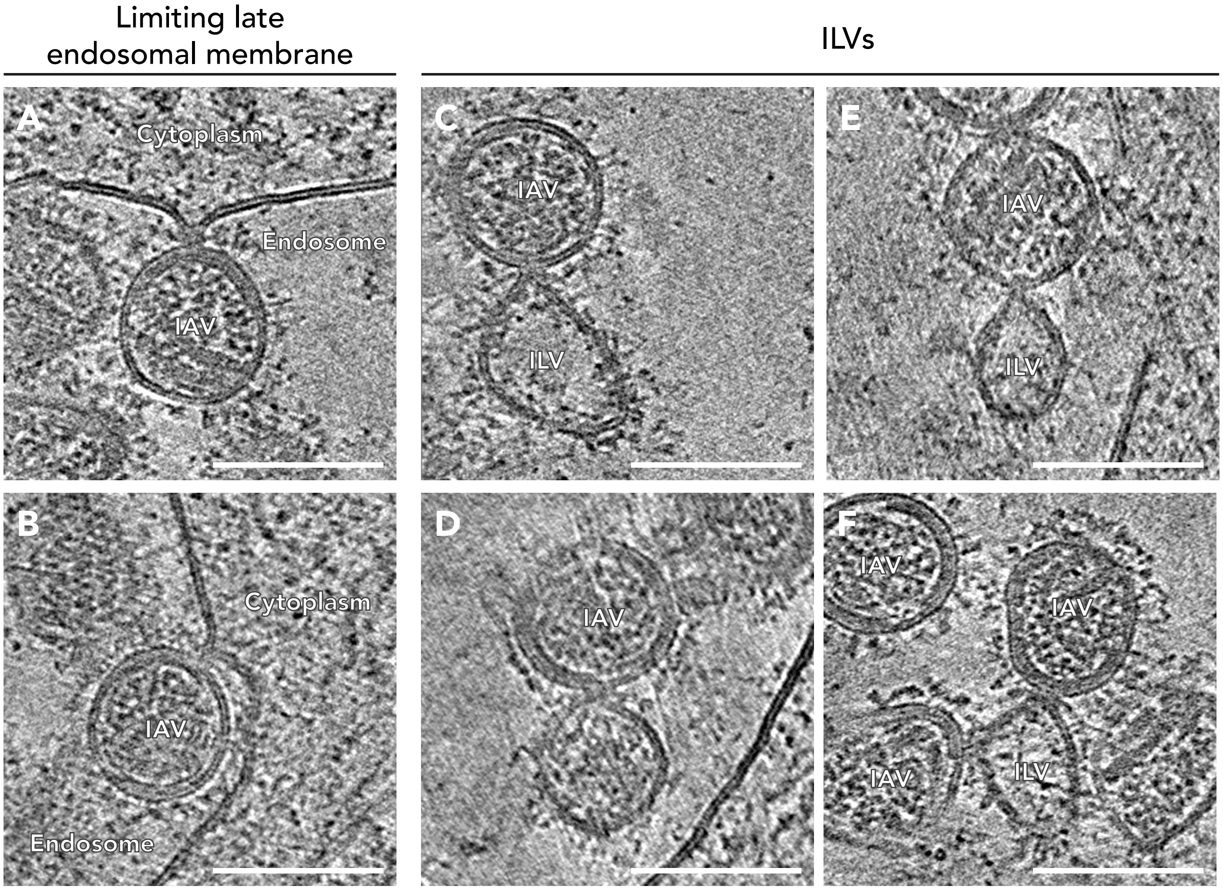


**Supplementary Figure 15 (related to Figure 6). Gallery of hemifusion sites during IAV membrane fusion in A549 cells.** A549 cells were infected with A/WSN/1933-PA-mScarlet (MOI = $3\times10^4$) and plunge frozen between 18 and 31 mpi. **(A and B)** IAV particles in hemifusion state at the limiting late endosomal membrane. **(C – F)** IAV particles in hemifusion state at ILV membranes in the endosomal lumen. Scale bars: 100 nm.
